# Mechanism-Guided Engineering of Fluorinase Unlocks Efficient Nucleophilic Biofluorination

**DOI:** 10.1101/2025.07.28.666932

**Authors:** Michaela Slanska, Daniel Christoph Volke, Isabel Pardo, Antonin Kunka, Naveen Banchallihundi Krishna, Anuj J. Shetty, Likith Muthuraj, Gladstone Sigamani, Roopa Lalitha, Alexander Kai Büll, Martin Marek, Pravin R. Kumar, Jiri Damborsky, Pablo I. Nikel, Zbynek Prokop

## Abstract

The fluorinase enzyme, the only known biocatalyst forming stable carbon–fluorine bonds, operates with extremely low efficiency, catalyzing one reaction every 2–12 minutes. This severely limits its utility for sustainable biofluorination, and its sluggish activity remains poorly understood. We suppressed its aggregation through directed mutagenesis and elucidated the kinetic mechanism using a novel mathematical framework that fits complex kinetic and oligomerization data. This analysis revealed that >80% of enzyme molecules are inactive under standard conditions due to two dead-end pathways. The designed W50F+A279R mutant preferentially formed hexamers and displayed enhanced catalytic efficiency in this oligomeric state. When coupled with mechanism-based optimization of the reaction medium, including enzymatic removal of the inhibitory product, the catalytic turnover rate reached 12.5 ± 2.1 min⁻¹, representing ∼60-fold increase compared with previously reported turnover rates of the wild-type enzyme. Our work provides a mechanistic blueprint for fluorinase enhancement and a generalizable mathematical framework for analyzing kinetics of multimeric enzymes.

## Introduction

Fluorinated organic compounds (organofluorines) have become indispensable in many industries and an integral part of everyday life in the developed world (**Fig. 1a**). In the pharmaceutical industry, fluorine often improves drug absorption, distribution, and excretion, as well as metabolic stability, efficacy, and selectivity^1,2^. Consequently, of all pharmaceuticals approved by the United States Food and Drug Administration between 2018 and 2025 (398 molecules), 24% are fluorinated structures (**Fig. 1b**)^3,4^. The number is even higher for blockbuster drugs, with 36% small-molecule drugs fluorinated as of 2023^5,6^. For instance, atorvastatin/Lipitor, a fluorinated drug used to treat hypercholesterolemia, was once the best-selling drug on the market^7,8^. Fluorine is also integral to the agrochemical industry; its incorporation into modern agrochemicals contributes to their efficacy and economic viability^9^. Of all pesticides approved by the International Organization of Standardization (ISO) between the years 2018 and 2025 (97 products), 59% are fluorinated^10^ (**Fig. 1b**). Furthermore, the biggest increase (>50%) in the fluorochemical demand is attributed to fluoropolymers, which have applications in practically all industrial sectors^11^. One such example is polytetrafluoroethylene^12^, which has revolutionized the material industry and is now used worldwide as part of cookware (Teflon), protective clothing (Goretex), and many other applications^13^.

**Fig. 1:**
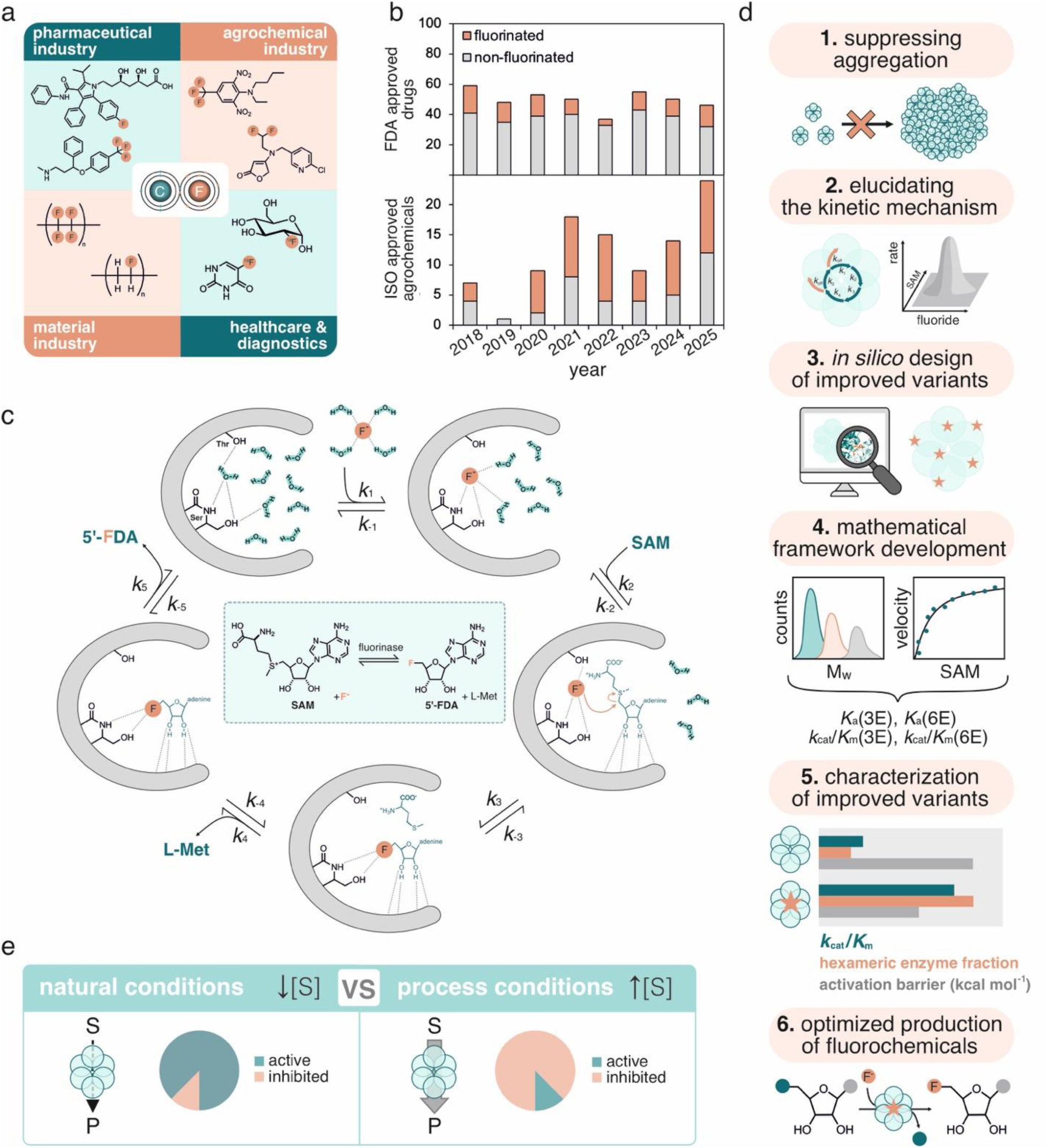
Mechanism-driven engineering of fluorinase. (**a**) Examples of organofluorines used in various industries (pharmaceutical industry – atorvastatin, fluoxetine; agrochemical industry – benfluralin, flupyradifurone; material industry – polyfluoroethylene, polyvinylfluoride; healthcare and diagnostics – 2-[^18^F]fluoro-2-deoxy-D-glucose, [^18^F]-5-fluorouracil). (**b**) Fluorinated and non-fluorinated pharmaceutical drugs and agrochemicals approved by the FDA and the ISO, respectively, in 2018–2025. (**c**) Reaction scheme and mechanism of fluorinase^37^. *k*_1_ - hydrated fluoride binds to the binding site, and some hydrogen bonds to water molecules are replaced by contacts with residues; *k*_2_ – specific binding of SAM, the remaining water is forced out of the binding site; *k*_3_ – S_N_2 nucleophilic substitution; *k*_4_ – release of Met; *k*_5_ – release of 5’-FDA. SAM – *S*-adenosyl-L-methionine, L-Met – L-methionine, 5’-FDA – 5’-fluoro-5’-deoxyadenosine. (**d**) Workflow of rational engineering of fluorinase for increased production yields of organofluorines. (**e**) Limitations of fluorinase under natural conditions (low substrate concentrations) and process conditions (high substrate concentrations). Under natural conditions, fluorinase is present mostly in its active form, but has very low turnover rates due to undersaturation by the substrates. The substrate concentrations need to be increased to increase the turnover rate under process settings, however, resulting in substrate inhibition.

Despite the widespread use of organofluorines, forming the carbon-fluorine bond remains challenging^1^. Current approaches rely exclusively on chemical methods, using non-renewable reagents and substrates, operating under extreme conditions, and often involving harmful chemicals^11,13^. Finding an efficient biosynthetic route is of high interest to achieve safer and more environmentally friendly production of these chemicals. Unfortunately, organofluorines are scarce in nature^14,15^, and so are the enzymatic pathways to their production. Out of roughly five thousand identified halometabolites to date, only thirteen natural fluorometabolites have been reported^16,17^. This is attributed to fluorine’s properties, e.g., its high redox potential (–2.87 eV, below that of hydrogen peroxide, –1.8 eV)^18^, which renders the haloperoxidase-type mechanism typical for chloride’s and bromide’s incorporation impossible for fluorine. Fluorine’s high electronegativity, higher than any other atom’s in the periodic table, precludes oxidation strategies involving the formation of halonium ions (X^+^) or halide radicals (X·)^19^. Furthermore, although the fluoride ion (F^−^) can act as a strong nucleophile, the ion is tightly hydrated and thus practically inert in aqueous environments^20^.

The fluorinase enzyme (5’-fluoro-5’-deoxyadenosine synthase, FlA)^21–27^ is the only known natural biocatalyst capable of forming stable carbon-fluorine bonds. Fluorinases are relatively rare enzymes, and they have been isolated in a few bacterial species and an Archaea. The fluorinase catalyzes the S_N_2 reaction between *S*-adenosyl-L-methionine (SAM) and F^−^ to generate 5’-fluoro-5’-deoxyadenosine (5’-FDA) and L-methionine (L-Met) (**Fig. 1c**)^21^. This enzyme is pivotal in developing biotechnological organofluorine synthesis (biofluorination), an unrealized frontier in modern biocatalysis. However, the catalytic efficiency reported for fluorinases is incredibly low, with turnover numbers (*k*_cat_) ranging between 0.083 and 0.41 min^−1 22,26^. The reason for this low efficiency is unknown, which hinders enzyme engineering efforts. Furthermore, fluorinase has poor solubility and is prone to aggregation, which constrains transient kinetic studies. Although some efforts have aimed to use the fluorinase as a gateway for fluorine incorporation into living cells^28–30^, these attempts have been hindered by the limitations described above.

Herein, we engineered the archetypal fluorinase FlA1 from *Streptomyces* sp. strain MA37 (**Fig. S1**) to enhance its solubility, enabling elucidation of its kinetic mechanism, uncovering the rate-limiting steps and two inhibitory pathways underlying the enzyme’s low catalytic efficiency. We further developed a novel mathematical framework to integrate complex kinetic information with oligomerization data and leveraged these insights and computational modelling to rationally design improved fluorinase variants. Additionally, we used our mechanistic understanding and knowledge of oligomerization dynamics to optimize the reaction medium for enhanced biocatalysis (**Fig. 1d**). Notably, we discovered that at substrate concentrations relevant to process-scale conditions, the majority of the enzyme becomes inhibited and thus catalytically inactive (**Fig. 1e**). Our findings demonstrate that a combined optimization of both the biocatalyst sequence and process parameters represents a promising strategy to improve the efficiency of biofluorination. Considering the highly conserved sequences among bacterial fluorinases (> 85% sequence similarity, **Fig. S1**), the results reported here can likely be extrapolated to other fluorinases apart from FlA1. Furthermore, the novel mathematical framework described here can be used to investigate oligomerization-dependent kinetics of other multimeric enzymes.

## Reducing Aggregation Enables Transient Kinetic Analysis

Transient kinetic experiments, needed for the determination of the kinetic mechanism of the fluorinase, require high amounts of protein in preparations where the enzyme concentration is precisely known. However, initial biochemical characterization revealed a high aggregation propensity of the fluorinase FlA1 from *Streptomyces* sp. strain MA37 (**Fig. S2**). Only 29 ± 9% of FlA1 was produced in soluble form when overexpressed in *Escherichia coli*, complicating protein purification and reducing the overall yield. Correspondingly, size-exclusion chromatography revealed significant aggregation in the purified fluorinase sample. To elucidate the cause of the aggregation, we tested its dependence on external factors by following static light scattering (SLS), which increases with increased particle size^31^. **Fig. 2a** displays aggregation monitored during a 10-h incubation at varying temperatures, revealing a clear concentration and temperature dependence of the scattering signal. At 37 °C, the aggregation was very rapid. Although decreasing the temperature to 30 °C and 25 °C decelerated precipitation, significant protein loss still occurred in a matter of minutes.

**Fig. 2:**
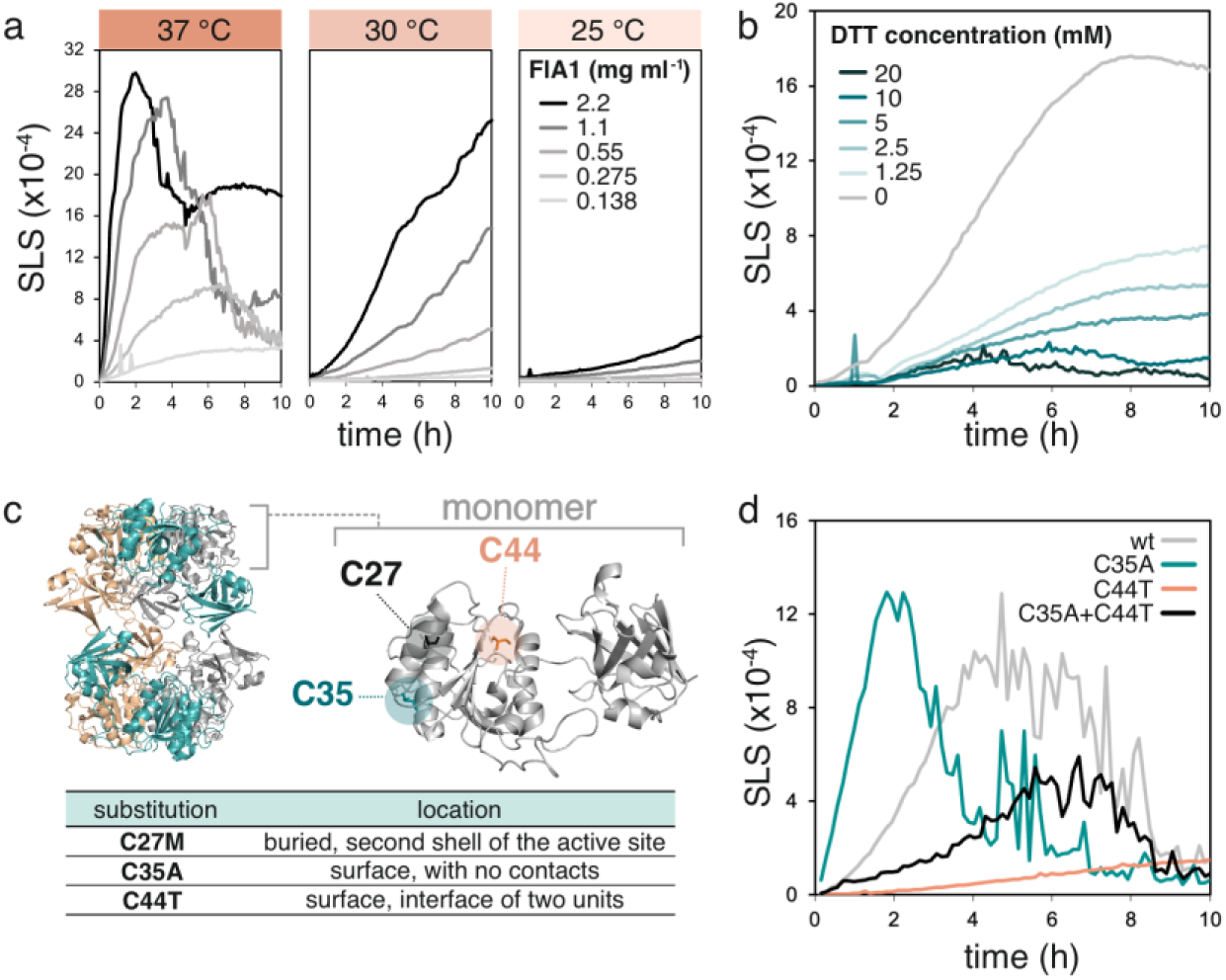
Engineering of fluorinase variants with suppressed aggregation and improved solubility. (**a**) Aggregation of FlA1 followed by static light scattering (SLS) at 266 nm at varying temperatures and enzyme concentrations in 30 mM HEPES, 75 mM KF (pH 7.8). (**b**) Effect of dithiothreitol (DTT) on the aggregation of FlA1 (1.6 mg ml^-1^) followed by SLS at 266 nm at 30 °C in 30 mM HEPES, 75 mM KF (pH 7.8). (**c**) The structure of FlA1 with highlighted cysteine residues and substitutions based on evolutionary analysis conducted using HotSpot Wizard^32^. (**d**) Aggregation of FlA1 variants (0.3 mg ml^-1^) with substituted cysteines followed by SLS at 266 nm at 37 °C in 30 mM HEPES, 75 mM KF (pH 7.8).

We found that increasing concentrations of the reducing agent dithiothreitol substantially reduced the rate and amplitude of the scattering signal (**Fig. 2b**). This indicated that the cysteine residues in the fluorinase sequence are likely responsible for FlA1’s high aggregation propensity. The monomeric form of fluorinase contains three cysteine residues, i.e., C27, C35, and C44 (**Fig. 2c**). We examined their spatial arrangement within the quaternary structures and found that, although their positions in the crystal structure do not favour disulfide bridge formation, such interactions may be possible within oligomeric assemblies. We targeted the C35 and C44 residues for mutagenesis because they are solvent-exposed, unlike C27, making them more likely to influence aggregation propensity. Their positioning further from the active site also minimizes potential disruption of catalytic activity. The *in-house* tool HotSpot Wizard^32^ was used to analyze the spatial location of cysteine residues and to predict amino acids most suitable for substitution at these positions based on their frequencies in homologous sequences (**Fig. 2c; Table S1**). Based on this analysis, three new fluorinase variants were constructed: C35A, C44T, and C35A+C44T.

The oligomeric state, which is crucial for fluorinase activity, was preserved in all the variants (**Fig. S3a**). It was found that variant C35A led to a complete loss of activity, but C44T and C35A+C44T only slightly decreased it (**Fig. S3b**). Importantly, the mutation C44T showed a significant deceleration of aggregation (**Fig. 2d**). At the temperature used for kinetic measurements (30 °C), the scattering signal of the C44T variant remains low and stable (**Fig. S4**), indicating its suitability for kinetic analysis. Accordingly, this FlA1 mutant was used in subsequent kinetic experiments to elucidate the fluorinase reaction mechanism.

## Two Inhibitory Pathways and Enzyme Undersaturation Revealed by Kinetic Modeling

The reason for the low catalytic efficiencies of fluorinases is mostly unknown, which prevents rational engineering of better variants applicable in the industrial-scale production of fluorochemicals. To fill the knowledge gap, we characterized FlA1 kinetics by steady-state and pre-steady-state (burst) experiments in both forward and reverse directions, globally fitted the data to a proposed kinetic model (**Fig, 3a, Figs. S5-S6**), and validated parameter reliability via FitSpace analysis (**Fig. S7**, **Table S2**); the enzyme remained hexameric under all conditions (**Fig. S8**).

The nucleophilic substitution step leading to the carbon-fluorine bond formation was rapid (*k*_+3_ = 102 min^−1^). Fluoride is a weak nucleophile in aqueous environments but a strong nucleophile in aprotic solvents^33^. While the substitution reaction is virtually impossible in water, the enzyme adopted a desolvation strategy to create a local aprotic environment in which the fluoride ion can effectively attack the C5’ of SAM. Mechanistically, the active site initially facilitates fluoride entry via transient interactions with water molecules, allowing F^−^ to access and bind within the catalytic pocket, where it is stabilized by interactions with catalytic residues (**Fig. 1c**). Upon SAM binding, these water molecules are displaced, resulting in a largely aprotic environment that promotes nucleophilic attack. In addition, the positively charged sulfonium center of SAM further activates the C5′ carbon, electrostatically favoring attack by the fluoride ion (**Fig. 1c**).

The rate-limiting step in the fluorination reaction, assuming sufficient fluoride concentration, is the release of 5’-FDA (*k*_+5_ = 17.1 min^−1^). Additionally, 5’-FDA acts as a substrate for the reverse reaction (defluorination) and displays tight binding to the apoenzyme (*k*₋₅ = 219 μM^−1^ min^−1^, *K*_d,5’-FDA_ = 78 nM), decreasing the fraction of fluorinase operating in the forward direction.

Furthermore, the kinetic analysis of C44T identified several additional factors limiting the catalytic performance of FlA1. Notably, the enzyme exhibits low affinity for fluoride (*K*_d,F⁻_ = *k*₋₁/*k*₊₁ = 151 mM). While this limitation can be overcome *in vitro* by applying high fluoride concentrations, such conditions are not feasible in an *in vivo* context due to fluoride toxicity^34^. In natural environments, fluoride concentrations are low (200 mg kg^−1^ in soil^35,36^), supporting only modest catalytic activity (**Fig. 3b**), but crucially, avoiding the formation of inactive enzyme-substrate complexes (**Fig. 3c**). For industrial applications, significantly higher concentrations of substrates would be required to achieve useful productivity (**Fig. 3b**). However, under these process conditions, inhibitory pathways become dominant, with over 80% of FlA1 sequestered in unproductive states. Consequently, even under optimized substrate conditions, fluorination turnover number remains low, approximately 24-fold slower than the rate-limiting step of 5’-FDA release. This indicates substantial untapped catalytic potential and suggests that the effective turnover is suppressed by additional inhibitory effects.

**Fig. 3:**
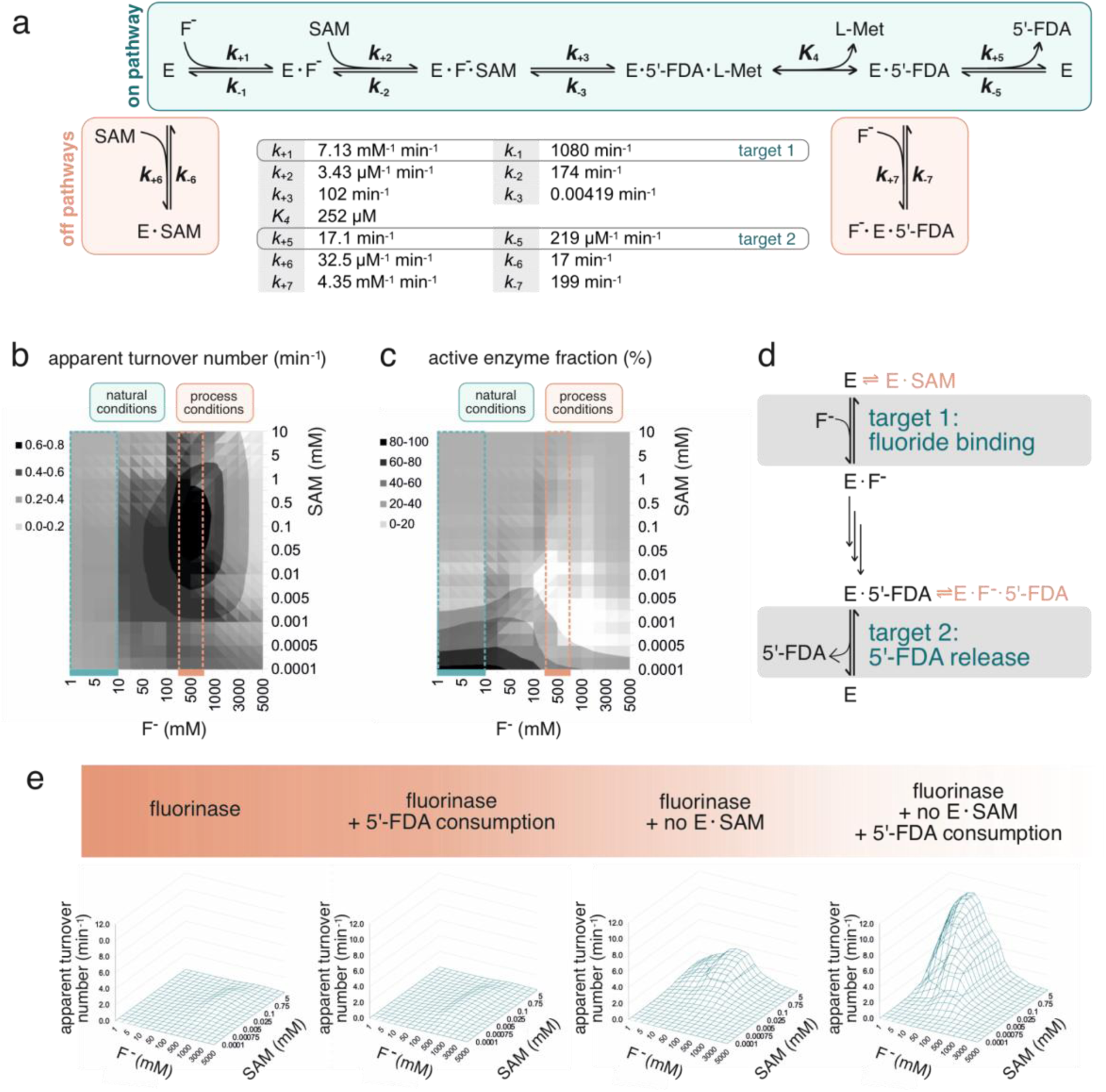
Kinetic mechanism of fluorinase. (**a**) Ordered sequential mechanism of FlA1+C44T with two inhibitory pathways, which trap the enzyme in unproductive states. E – enzyme (FlA1+C44T), SAM – *S*-adenosyl-L-methionine, L-Met – L-methionine, 5’-FDA – 5’-fluoro-5’-deoxyadenosine. The kinetic constants of each step in the catalytic cycle are displayed. The mechanism was derived from experiments at 30 °C in 30 mM HEPES, 150 mM NaCl, and pH 7.8. (**b**) Dependency of the rate of 5’-FDA formation on F^-^ and SAM concentrations. (**c**) Dependency of the fraction of active enzyme species (E, E·F⁻, E·F⁻·SAM, E·5’-FDA·L-Met, E·5’-FDA) on F^-^ and SAM concentrations. (**d**) Targets for the engineering of more efficient fluorinases. (**e**) Predicted increase in fluorination turnover number due to enhanced 5’-FDA consumption and suppression of inhibition by SAM (no E·SAM). The simulation data shown in panels **b**, **c**, and **e** were generated using KinTek Explorer^61^ based on the kinetic mechanism and determined kinetic constants (panel **a**). The simulated parameter, apparent turnover number, reflects product formation per unit of enzyme and time under defined conditions, without assuming kinetic saturation.

One of the inhibitory pathways is a previously described competitive inhibition by SAM^37^, which binds the free enzyme to form the E⋅SAM complex. Steered molecular dynamics and umbrella sampling simulation studies revealed that in the presence of SAM in the active site, F^−^ cannot reach the active site residues but instead forms a stable hydrogen bond with non-catalytic amino acids T155 and S269 (**Fig. S9**). The second off-pathway is the unproductive binding of fluoride to the enzyme–product complex E⋅5’-FDA to form F^−^⋅E⋅5’-FDA. A metadynamics study of the E⋅5’-FDA and F^−^⋅E⋅5’-FDA complexes revealed that this inhibition occurs by a bridging interaction between enzyme residues, F^−^ and 5’-FDA (**Figs. S10-S11**). Metadynamics is an atomistic enhanced-sampling molecular dynamics method that applies a history-dependent bias along selected collective variables to overcome local free-energy minima. The bias, constructed from Gaussian potentials, accelerates transitions between minima and enables efficient sampling of rare events within accessible simulation timescales.

To increase the overall catalytic efficiency of fluorinases, the amount of enzyme trapped in the unproductive states E·SAM and F⁻·E·5’-FDA needs to be decreased (**Fig. 3d**). Eliminating unproductive SAM binding directly is challenging, as reducing its affinity for the free enzyme (off-pathway) would also impair its productive binding to the E·F⁻ complex (on-pathway). As an alternative solution to decrease E·SAM formation, we propose engineering the enzyme for increased affinity toward fluoride. Given that SAM binding to the apoenzyme competes with fluoride binding, this emerges as a promising target to address both major limitations (i) the low affinity towards fluoride itself and (ii) the competitive formation of the unproductive E·SAM complex. **Fig. 3e** shows that eliminating formation of the E·SAM complex would significantly increase the apparent turnover number of the fluorination reaction, reaching 3.6 min^−1^ at optimal substrate concentrations. In combination with 5’-FDA consumption in downstream reactions, an apparent turnover number of up to 10 min^−1^ appears achievable.

## Mutations Design Guided by Fluoride Binding Simulations and Quantum Chemical Calculations

The kinetic analysis revealed two inhibitory pathways that inactivate fluorinase: (i) SAM competitively binds the free enzyme, blocking F⁻ access, and (ii) F⁻ binds to the enzyme-product complex (E·5’-FDA), trapping the enzyme in another inactive complex (F⁻·E·5’-FDA). Enhancing initial F⁻ binding to the apoenzyme emerged as a promising approach to improve the enzyme’s catalytic efficiency. This would increase the overall rate of the on-pathway leading to 5’-FDA formation and limit the competing inhibition of the apoenzyme by unproductive binding of SAM. To identify mutations favouring fast fluoride binding, we conducted a comprehensive *in silico* investigation of the fluoride-fluorinase interaction. The simulations were performed in an aqueous environment, consistent with the native conditions under which the enzyme operates, and to mimic the conditions of the kinetic studies. Computer modelling delineated F⁻ diffusion into the catalytic site. Subsequently, smooth global biased simulations were used to generate data processed via a grid-based interaction profiling method to predict mutational hotspots.

The stepwise diffusion process, obtained from Gaussian Accelerated Molecular Dynamics simulations, is shown in **Fig. 4a**. The diffusion pathway is visualized via F⁻ clusters colored by potential of mean force (PMF) values, mapping the free energy landscape of ion movement. The most critical step was the dehydration of F⁻, requiring hydrophilic surface residues to strip water molecules while guiding the ion. Consistently, four water molecules surrounded F⁻ in a defined geometry (**Fig. S12, Movie S1**). The regular hydrogen-bonding interactions between solvent water molecules and F⁻ ions illustrate the weak nucleophilic character of fluoride outside the enzyme microenvironment. Simulations identified W50, Y77, P78, A79, A255, and N278 as key facilitators of dehydration. The backbone atoms of P78, A79, and A255, and the sidechain of N278 were observed to interact with F⁻, forming stable hydrogen bonds. These residues acted as guides to move the F⁻ towards the active site. As F⁻ diffused further, the backbone of Y77 interacted with F⁻, directing it towards the catalytic region where it interacts with T80 and S158 (**Fig. S13, Movie S1**). W50 was identified as a key residue that hinders both the binding of the fluoride ion (F⁻) to the active site and the release of products following the reaction. Umbrella sampling simulations with 0.5 Å windows revealed that F⁻ can diffuse into the apo-enzyme (**Fig. 4b**). However, in the presence of SAM, F⁻ adopts a non-catalytic pose due to stabilizing interactions with residues T155 and S269 (**Fig. S9**). To modulate F⁻ and SAM binding without compromising the structural integrity of the fluorinase, a conservative W50F mutation was introduced. This substitution aimed to reduce the steric hindrance posed by W50 while preserving essential interactions and hydrophobicity.

**Fig. 4:**
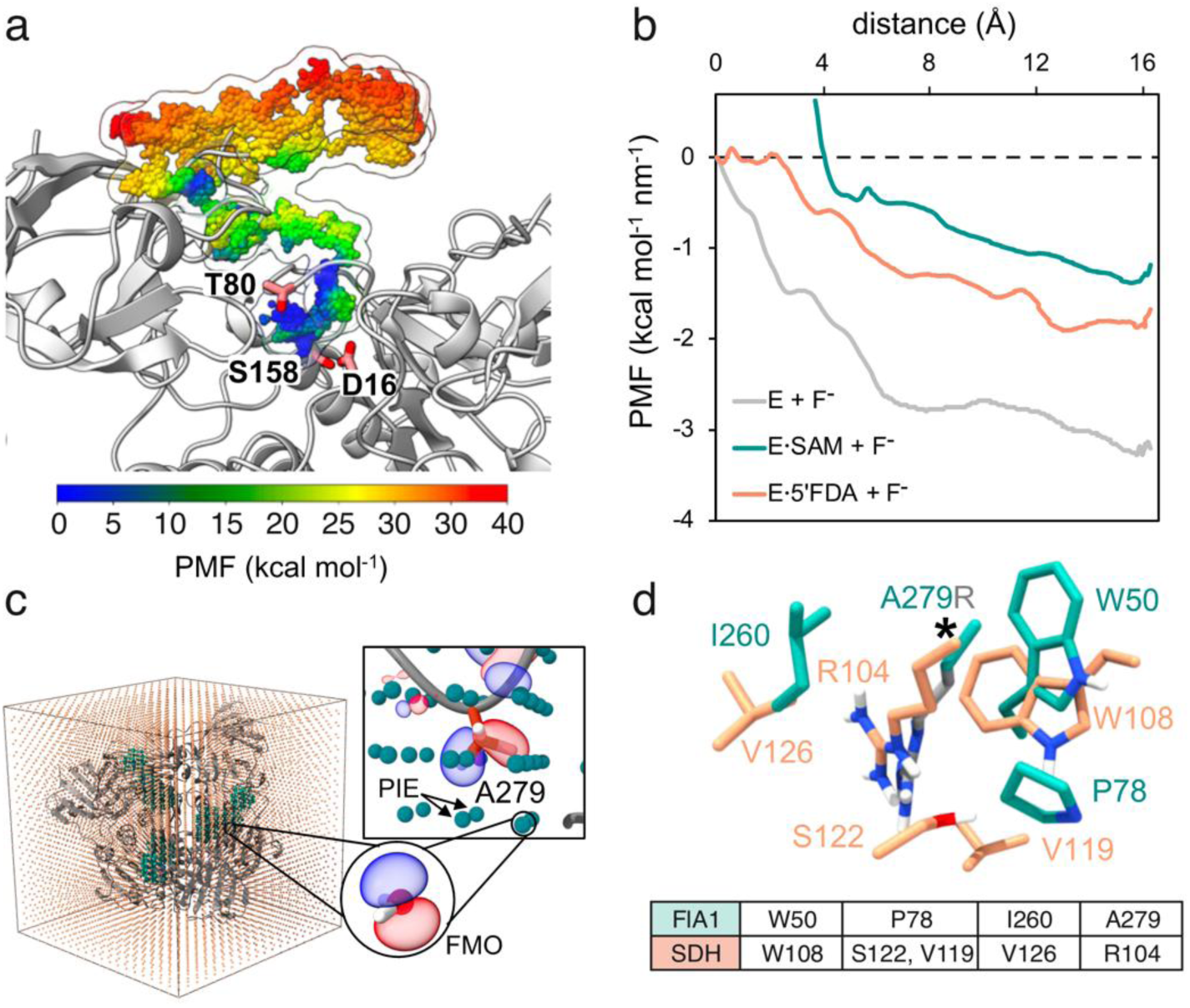
Mechanistic insights into fluoride diffusion and grid-based mutant design. (**a**) Gaussian accelerated molecular dynamics simulation showing F⁻ diffusion from solvent into the enzyme through a specific entry point, following an energetically favorable pathway to the active site. The pathway is visualized using F⁻ clusters colored by potential of mean force values (PMF), calculated as a function of two collective variables: the distance between F⁻ and S158 (CV1) and the distance between F⁻ and T80 (CV2), using second-order cumulant expansion. The enzyme is shown as a grey ribbon, with key active-site residues depicted as sticks. PMF values (kcal mol^-1^) are color-coded: blue-green for low-energy zones (favorable, 0-15 kcal mol^-1^), green-yellow for moderate energy zones (metastable, 15-25 kcal mol^-1^), and orange-red for high-energy zones (unfavorable, 25-40 kcal mol^-1^). (**b**) PMF profiles of F⁻ diffusion into the active site from umbrella sampling for three enzyme states: apoenzyme (E+F⁻), enzyme–SAM complex (E·SAM+F⁻), and enzyme–5’-FDA complex (E·5’-FDA+F⁻). (**c**) Conformational ensemble of enzyme–F⁻ complexes from diffusion studies mapped onto a 3D grid (2.0 Å spacing), capturing pair interaction energy (PIE) values between amino acid side chains and placed molecular probes. The residue A279 is identified as an energetic hotspot based on high cumulative PIE values. Residues are shown as orange sticks, with interacting probes depicted as teal spheres and orbital surfaces visualized. (**d**) 3D overlay of the A279 region in FlA1 with the corresponding region from succinate dehydrogenase (SDH, PDB ID 1YQ3) based on probe-derived matching. FlA1 residues are shown as teal sticks; corresponding mutation side chains as grey sticks; residues from the database structure as peach sticks. The asterisk indicates A279 as a mutation hotspot with a high PIE value matched to a substitution A279R.

To guide mutation design, enzyme–substrate conformations from the F⁻ diffusion pathway, were analyzed. The conformational ensemble of FlA1–F⁻ complexes was aligned and projected onto a 3D grid with 2.0 Å resolution. At each grid point, molecular probes representing amino acid side chains or water molecules were placed, and pair interaction energy (PIE) values were computed using the Fragment Molecular Orbital (FMO) method (**Fig. 4c**). The residue A279 emerged as a distinct interaction hotspot, showing a high PIE value indicative of strong stabilizing potential. Independently, this residue was identified as a hotspot in a directed evolution campaign by Sun *et al.*^38^, suggesting the potential effect of mutations at this position. Notably, probes interacting near A279 mirrored the interaction network of known active-site motifs in avian succinate dehydrogenase (SDH, PDB ID 1YQ3), particularly involving residues W108, S122/V119, V126, and R104 (**Fig. 4d**). The spatial proximity of these residues arises from hexamer formation of FlA and was therefore considered in the design of mutations for experimental validation. Inspired by the functional role of R104 in SDH, the A279R mutation was proposed to introduce a similar positively charged side chain. The W50F and A279R substitutions were individually introduced into the wild-type fluorinase via site-directed mutagenesis. Subsequently, a double mutant, W50F+A279R, was constructed to evaluate potential synergistic effects.

## Engineering Fluorinase for Enhanced Catalysis via Hexamer Formation and Inhibition Suppression

The wild-type fluorinase, its variant with suppressed aggregation (C44T) and its variants designed for tailored fluoride binding (W50F, A279R, and W50F+A279R) were kinetically and structurally characterized by (i) a conventional steady-state analysis that revealed the presence of multiple oligomeric enzyme forms, (ii) a detailed analysis of the oligomeric states, which revealed the influence of substrates on oligomerization, and (iii) a global numerical analysis, providing a detailed understanding of fluorinase structural dynamics and catalytic performance. The refined results of this complex analysis are summarized in **Fig. 5**, with detailed descriptions of the individual methodological components provided in the **Supplementary texts S27 and S28**.

**Fig. 5:**
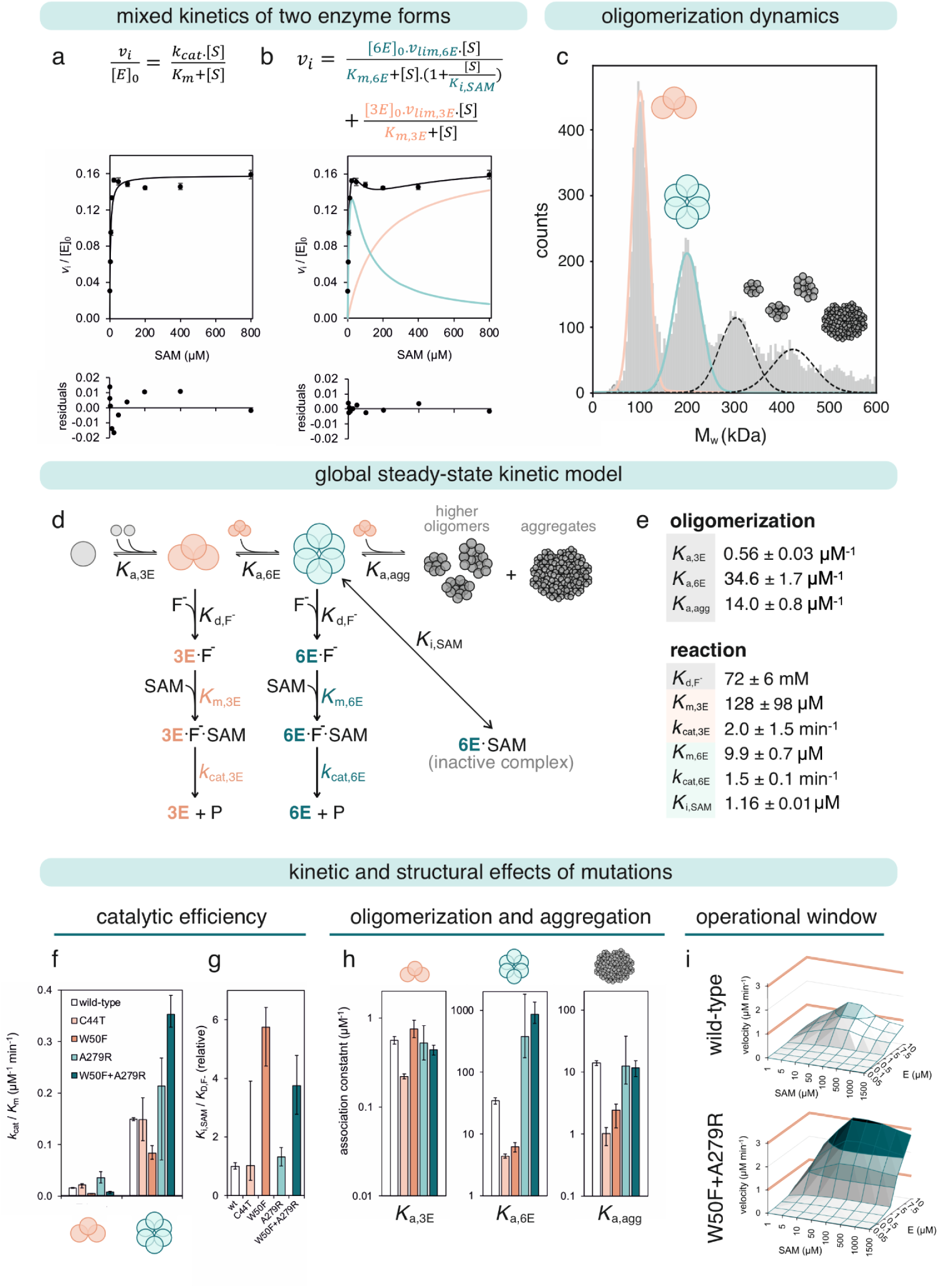
Catalytic performance and oligomerization dynamics of fluorinase and its mutants. Analytical fitting of the steady-state kinetic data of wild-type FlA1. (**a**) The best fit for the hyperbolic Michaelis-Menten mechanism, and (**b**) a model accounting for two enzyme species. The kinetic data were collected with 1.56 to 800 *μ*M of SAM, 1 *μ*M FlA1, 75 mM KF in 50 mM HEPES, pH 7.8 at 37 °C. Each data point represents the mean of three independent replicates; error bars indicate the standard deviation. (**c**) Mass photometry analysis of fluorinase oligomerization. The histograms of the binding event counts were fitted to the sum of five Gaussian distributions. The data were recorded for the wild-type FlA1 at a concentration of 500 nM in 30 mM HEPES buffer, pH 7.5, 20°C. (**d**) The minimal steady-state kinetic model is defined collectively by the results from analytical fitting, mass photometry analysis, and observations of the active site titration. (**e**) Kinetic parameters of wild-type FlA1 with standard errors (s.e.) estimated by nonlinear regression. Due to the correlation between these individual parameter values, the secondary analysis used their ratio, *k*_cat_/*K*_M_. (**f**) Consistent comparison for both trimer and hexamer catalytic efficiency across all tested variants. (**g**) The active enzyme fraction involved in the reaction is defined by *K*_i,SAM_/*K*_d,F⁻_ ratio, where K_i,SAM_ is the equilibrium constant for the dissociation of the inhibitory E⋅SAM complex, and *K*_d,F⁻_ is the equilibrium constant for the dissociation of an active E⋅F- complex. (**h**) The equilibrium association constants for monomer to trimer transition *K*_a,3E_, formation of a hexamer *K*_a,6E_, and formation of higher oligomers *K*_a,agg_ leading to protein aggregation. Error bars in (**f-h**) represent the confidence intervals (lower and upper limits) of the parameters determined by confidence contour analysis with a Chi^2^ threshold of 0.98 (**Table S4**). (**i**) The dependence of the initial reaction velocity of wild-type fluorinase and the W50F+A279R variant on the concentrations of *S*-adenosyl-L-methionine (SAM) and enzyme (E) at the standard concentration of fluoride 75 mM. The simulation data were generated in KinTek Explorer using the steady-state kinetic model (Fig. 6d) and the parameter estimates presented in **Table S4**.

The kinetic behavior of fluorinase and its mutants was first examined using a conventional steady-state analysis of initial velocities. Although the initial fit to the hyperbolic Michaelis-Menten model appeared to capture the general trend of the data, a closer examination of the residuals revealed significant systematic deviations, indicating that the model did not fully capture the observed kinetics (**Fig. 5a**). Substrate inhibition was indicated at SAM concentrations up to 200 μM (**Fig. S20b**), while higher concentrations showed a closer fit to the hyperbolic model (**Fig. S20c**), suggesting a mixed kinetic profile. A refined model accounting for two enzyme species with distinct kinetic characteristics provided an accurate explanation of the observed data (**Fig. 5b**). This finding was consistent with prior studies^39^ showing distinct kinetics for separated trimeric and hexameric forms of fluorinase. To differentiate the contributions of co-existing trimeric and hexameric forms to the mixed kinetics, mass photometry was used to characterize the oligomerization states of fluorinase (**Fig. 5c, Fig. S14**). By employing an advanced hybrid analytical approach that integrates kinetic data with quantitative data on trimer and hexamer fractions, more reliable estimates of individual kinetic parameters were further refined (**Fig. S20d**). Still, the conventional analytical fitting remains inadequate for fully capturing the dynamics of the kinetic and structural behavior of fluorinase.

We applied global numerical fitting, integrating reaction kinetic data and structural data from mass photometry, to address this limitation (**Fig. 5d and Fig. S21**). Numerical analysis (i) retains all raw data without any loss of information, e.g., information on the active enzyme fraction excluded during conventional initial velocity estimation (**Fig. S19a**), (ii) eliminates the reliance on simplifying assumptions, such as steady-state approximations, and (iii) improves the accuracy and reliability of parameter estimates, particularly under non-linear conditions. In the initial phase, the global numerical analysis revealed that the oligomeric behavior data recorded for the free enzyme did not align with the catalytic data, a discrepancy undetectable by previous conventional analysis. These results prompted us to conduct a comprehensive study of fluorinase’s oligomeric behavior, comparing the enzyme’s free state with its structural changes in the presence of substrates. The results confirmed that the presence of substrates significantly affects oligomerization behavior (**Figs. S17-S18**). Parallel analysis of kinetic data alongside structural data, both recorded under identical reaction conditions, including the presence of substrates, enabled the global analysis to yield accurate and highly consistent results. This approach provided high-quality estimates of key kinetic parameters (**Fig. 5e, Table S4**), supported by robust statistical validation (**Fig. S23**). A detailed description of the global numerical analysis is provided in **Supplementary Text S28**.

Both conventional and numerical approaches consistently demonstrated that the hexamer exhibits significantly higher catalytic efficiency (*k*_cat_/*K*_m_) than the trimer, with an approximately tenfold increase (**Fig. 5f**). The enhanced efficiency of the hexamer is mainly due to a lower Michaelis constant (*K*_m_), rather than changes in the turnover number (*k*_cat_) (**Fig. 5e**). This finding is consistent with previous studies on fluorinase engineering, which demonstrated that the isolated trimeric and hexameric forms of the enzyme differ primarily in their *K*_m_ values, 206 ± 34 and 3.7 ± 0.7 μM, respectively, while exhibiting nearly identical *k*_cat_ values of 0.20 ± 0.01 and 0.22 ± 0.01 min^−1 39^.

Mutations, particularly A279R, further increased the hexamer’s catalytic efficiency, with the most pronounced effect observed in the double mutant W50F+A279R (**Fig. 5f**). Additionally, the W50F mutation increases the fraction of active enzymes involved in the reaction, indicated in active site titration data (**Fig. S19**) and characterized by nearly a sixfold increased *K*_i,SAM_/*K*_d,F-_ ratio (**Fig. 5g**). The positive impact of W50F on the active enzyme fraction was also mostly preserved in the double mutant W50F+A279R. Under standard conditions, the wild type enzyme is undersaturated and the fraction of active enzyme is lowered, mainly due to a high dissociation constant for the enzyme-fluoride complex (*K*_d,F-_ = 72 ± 6 mM) relative to typical fluoride concentrations used (75 mM). Additionally, the improper order of SAM binding (characterized by the equilibrium dissociation constant *K*_i,SAM_), further decreases the fraction of active enzyme in the reaction. Subsequent analysis of fluoride kinetics (**Fig. S22**) indicated that an increase in the active enzyme fraction primarily results from the mitigation of the substrate inhibitory effect of SAM, as fluoride *K*_m_ showed minimal differences between the wild-type enzyme and W50F+A279R mutant. This conclusion is supported by global numerical analysis, which yielded *K*_d,F-_ the values of 72 ± 6 mM and 71 ± 16 mM, and *K*_i,SAM_ values of 1.16 ± 0.08 μM and 10.6 ± 2.2 μM for the wild type and W50F+A279R, respectively.

The proposed mutations also had a significant impact on the structural behavior of fluorinase (**Fig. 5h**). While no notable effect was observed on trimer formation (*K*_a,3E_), both mutations, W50F and A279R, significantly influenced hexamer formation (*K*_a,6E_) and subsequent assembly into higher-order oligomers (*K*_a,agg_). The A279R mutation enhanced hexamer formation, making this variant, along with the double mutant W50F+A279R, predominantly hexameric (**Fig. S18**). In contrast, W50F reduced hexamer formation but mitigated aggregation propensity similar to C44T. The W50F+A279R double mutant efficiently formed the catalytically active hexamer, comparable to A279R alone, while maintaining an aggregation propensity similar to the wild type. The C44T mutation, which was designed to suppress aggregation, significantly decreased *K*_a,agg_ but also reduced trimer (*K*_a,3E_) and hexamer (*K*_a,6E_) formation efficiency, suggesting a trade-off between suppressed aggregation and oligomerization.

The dynamic nature of fluorinase behavior, influenced by the interplay of oligomerization and functional properties, makes predicting enzyme performance under varying conditions challenging. To address this, we used the global numerical model (**Fig. 5d**) to simulate enzyme behavior across different conditions, enabling a comparison of the operational space and performance optima for the wild type and the best-performing double mutant (**Fig. 5i**). While the wild-type enzyme exhibited a narrow activity range, with a pronounced decline in activity due to substrate inhibition at higher substrate concentrations, the W50F+A279R mutant demonstrated a broader operational window and achieved twice the activity of the wild type. These findings underscore the highly dynamic nature of fluorinase and its mutants, highlighting the potential for further optimization of catalytic performance through targeted mutagenesis.

## Metadynamics Reveals Structural Basis of Enhanced Fluorination and Hexamerization

Molecular modelling of the wild-type fluorinase and its mutants was carried out to understand the effects of introduced substitutions. Representative states along the minimum reaction path obtained by QM/MM metadynamics are illustrated in **Fig. S24**, following a stepwise mechanism with reactant, transition state, and product. In the reactant state, SAM is positioned in the active site, and the fluoride ion is stabilized through interactions with the catalytic residues T80 and S158, aligning for nucleophilic attack (**Movie S2**). QM/MM simulations were conducted using three simultaneous quantum regions (3 x 64 atoms) centered on the active sites. Two collective variables (CVs) were defined: CV1, representing the distance between the sulfur and C5′ of SAM, and CV2, representing the distance between F⁻ and C5′ of SAM.

The reaction free energy profiles shown in **Fig. 6a–d** describe the transition from the reactant state (CV1 ≈ 1.8 Å, CV2 ≈ 2.8–3.0 Å) to the product state (CV1 ≈ 3.4–4.0 Å, CV2 ≈ 1.2–1.4 Å), with energy barriers visualized by a gradient from purple (low) to red (high). Among the systems studied, the wild-type enzyme displays the highest activation barrier at 7.1 kcal mol^−1^. The A279R variant shows a moderate reduction in the barrier to 6.3 kcal mol^−1^, while the W50F mutant exhibits an intermediate profile with a barrier of 6.8 kcal mol^−1^. Notably, the double mutant A279R+W50F demonstrates the lowest energy barrier at 5.7 kcal mol^−1^, indicating the most favorable and efficient catalysis.

**Fig. 6:**
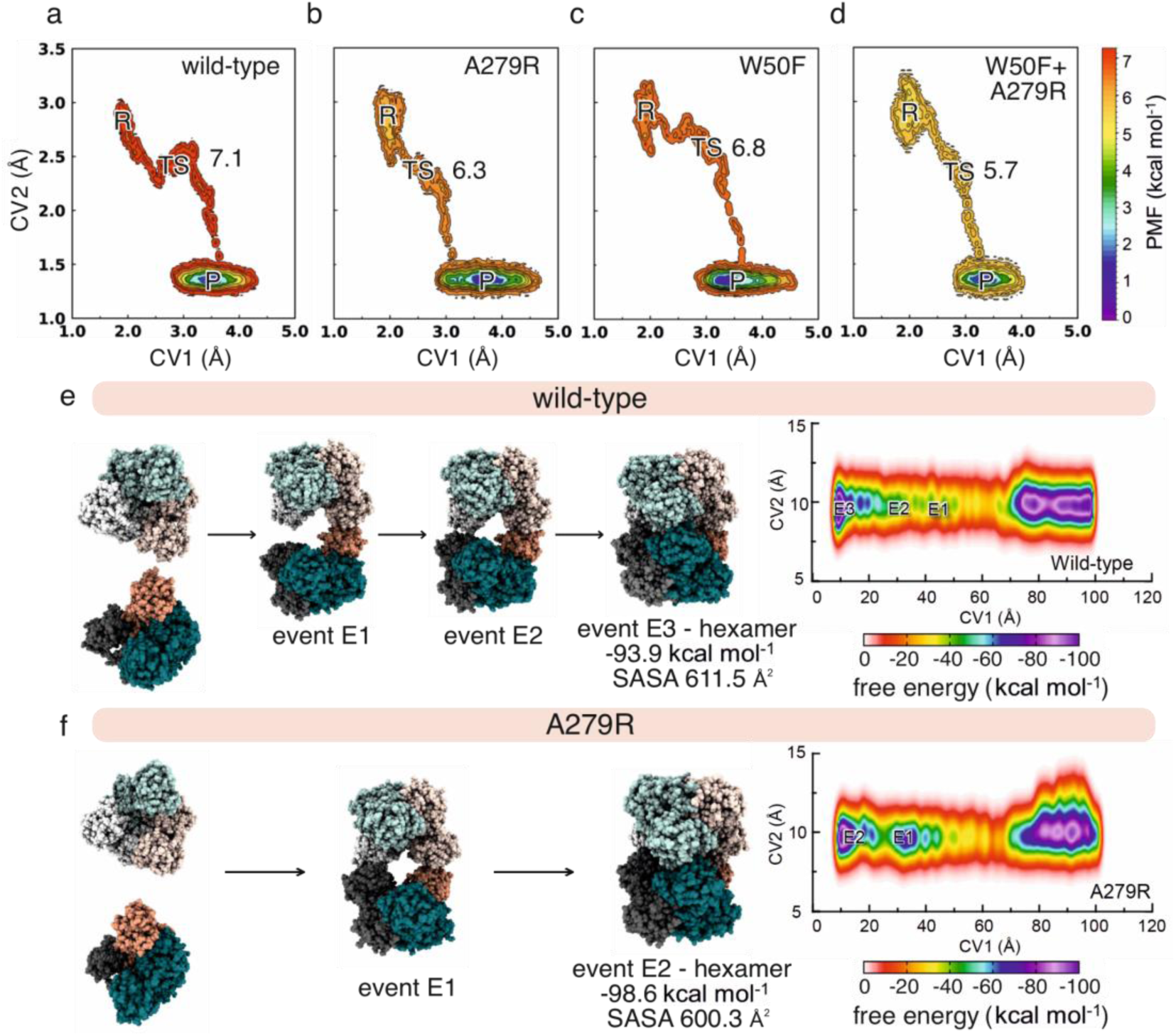
Computational analysis of the effect of mutations on fluorinase activity and oligomerization. Potential of mean force free energy landscapes depicting the catalytic reaction for the wild type (**a**) and mutants A279R (**b**), W50F (**c**), and W50F+A279R (**d**), plotted along two collective variables CV1 and CV2. CV1 represents the distance between the sulphur atom of SAM and C5’ atom of SAM, and CV2 represents the distance between F⁻ and SAM-C5’. The substrate (S), the transition state (TS) and product (P) are labelled. Activation energy barriers are indicated: wild type 7.1 kcal mol^-1^, A279R 6.3 kcal mol^-1^, W50F 6.8 kcal mol^-1^, and W50F+A279R 5.7 kcal mol^-1^, demonstrating reduced reaction barriers in engineered mutants compared to the wild-type enzyme (**Movie S2**). (**e**) Metadynamics studies on hexamer complex formation in wild-type FlA1 highlighted three events in the free energy landscape with the occurrence of events E1 followed by E2 leading to the formation of hexamer (E3, −93.9 kcal mol^-1^) and a solvent accessible surface area (SASA) value of 611.5 Å^2^. (**f**) Metadynamics studies on hexamer complex formation in A279R highlighted two events in the free energy landscape with the occurrence of event E1 leading to the formation of hexamer (E2, −98.6 kcal.mol^-1^) and SASA value of 600.3 Å^2^. In wild type and A279R, arrows indicate hexamer formation pathways. The A279R attains a hexamer state that is more compact and energetically more favorable than the wild type.

Next, we conducted well-tempered metadynamics simulations to explain the structural basis for favorable formation of the hexamer by A279R mutant (**Fig. 6e-f**, **Movie S3, Figs. S25, S28**). We observed that both wild type and A279R adopted distinct trimer conformations during assembly (**Supplementary Text S29**). A279R associated more readily, indicating a kinetically favorable pathway, while the wild type displayed delayed hexamer formation with prolonged intermediates (**Fig. S26**). Solvent accessible surface area analysis indicates the A279R attains a more compact and stable conformation (600.3 Å²) compared to the wild type (611.5 Å²). The enhanced stability in A279R is attributed to stronger inter-chain interactions. Specifically, R279 of N1 forms salt bridges with D51 and E53 of N2, stabilizing the trimer (**Fig. S27**). Additional stabilizing contacts between R279 and residues W50, I260, and N278, together with further inter-chain salt bridges, reinforce both the dimeric and trimeric interfaces. Importantly, the effect of the A279R mutation on promoting the formation of more active hexamers is orthogonal to the effect of W50F on increasing the fraction of catalytically active enzyme, providing a plausible explanation for the additive effects determined with the combined mutant W50F+A279R.

## Model-Guided Protein and Medium Engineering Enhances the Fluorination Efficiency by Two Orders of Magnitude

To test the optimal process conditions predicted by the kinetic model, the activity of wild-type fluorinase and W50F+A279R mutant was assayed under varying high substrate concentrations via end-point measurements after 2 h (**Fig. 7b, Fig. S29**). To circumvent the adverse effect of the product 5’-FDA (substrate for the reverse reaction – defluorination, *K*_d,5’-FDA_ = 78 nM, **Fig 3a**), we hypothesized that an enzyme capable of converting the fluorodeoxynucleoside into a downstream fluorometabolite could relieve FlA1 from the 5’-FDA-mediated effect. To this end, we selected an *S*-adenosyl-L-homocysteine nucleosidase from *E*. *coli* (MtnN), which hydrolyses 5’-FDA into adenine and 5’-fluoro-5’-deoxyribose, to be included in the enzyme reaction (**Fig. 7a**). In these enzymatic cascades, the apparent turnover number increased substantially: when 5’-FDA was consumed, the wild type enzyme showed values up to 5 min^−1^, while the mutant W50F+A279R exceeded 8.3 min^−1^ (**Fig. 7b**). Kinetic measurements, using the optimal substrate concentrations, 0.5 mM SAM and 500 mM NaF, confirmed high apparent turnover numbers of 3.7 ± 0.5 min^−1^ for the wild type and 12.5 ± 2.1 min^−1^ for the W50F+A279R mutant (**Fig. 7c**, **Fig. S31**). This result exceeds previously reported activities of the wild-type FlA1 enzyme, which ranged from 0.16 to 0.26 min^−1 22,26,39^, under low substrate concentrations (0.8 mM SAM and 0.075 M NaF) that mimic natural conditions but are suboptimal for industrial applications. At the same time, this dual enzymatic cascade produced 5’-fluoro-5’-deoxyribose stoichiometrically. This fluorinated sugar has multiple applications as a precursor of pharmaceutical antiviral and anticancer drugs^40^ and as a biomedical tracer^41^. Apart from this work, the strategy of using MtnN to consume 5’-FDA could be used to advance other previously reported fluorinase engineering efforts^27,38,42^.

**Fig. 7:**
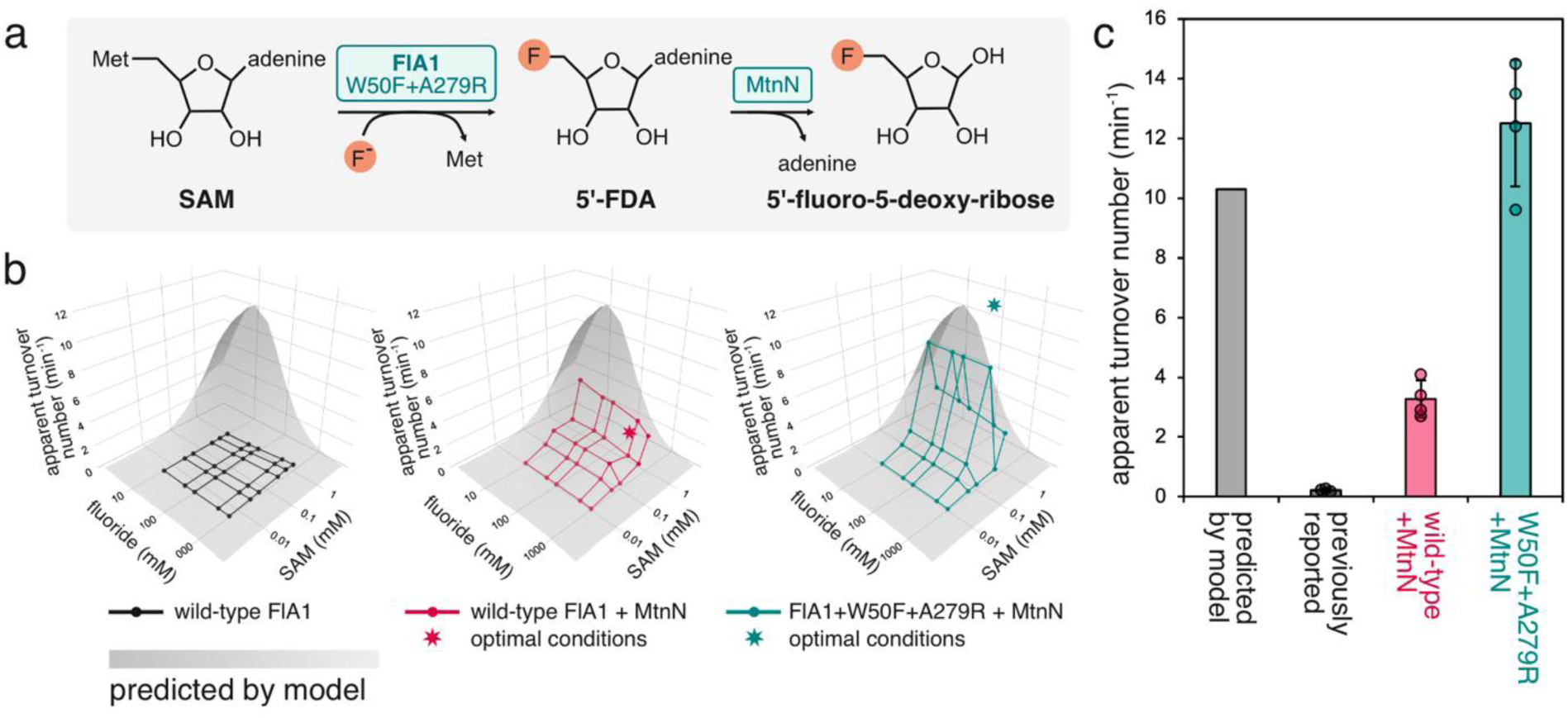
Protein and medium optimization for effective *in vitro* biofluorination. (**a**) Optimization of the fluorinase reaction was achieved by rational engineering of FlA1 (W50F+A279R) and by downstream coupling to *S*-adenosyl-L-homocysteine nucleosidase (MtnN), consuming 5’-FDA (substrate of the reverse reaction of FlA1). (**b**) Experimental validation of model-predicted apparent turnover numbers of FlA1 across varying concentrations of SAM and fluoride. The model prediction (grey, Fig. 3e) is based on the kinetic mechanism of FlA1+C44T (Fig. 3a) and simulates an idealized scenario in which inhibition by SAM is eliminated and 5′-FDA is efficiently consumed. The overlaid experimental datapoints (**Fig. S29**) were collected with 0.5 *μ*M FlA1 wild type or FlA1+A279R+W50F, 0.5 *μ*M MtnN, 0.005-1 mM SAM, 10-1000 mM NaF in 30 mM HEPES, pH 7.8 at 37 °C, and represent the end-point after 2 h incubation. The stars indicate maximum apparent turnover numbers calculated from initial rates (**Fig. S31**) measured at 500 μM SAM and 500 mM NaF. (**c**) Comparison of the maximum apparent turnover number predicted from the kinetic model with the previously reported values for wild-type FlA1 under suboptimal conditions (300-800 μM SAM, 75-200 mM fluoride) ^22,26,39^, and with the maximum experimentally achieved values following medium optimization (wild type FlA1 and FlA1+W50F+A279R, 500 μM SAM, 500 mM fluoride, 5’-FDA consumption by MtnN). The error-bars represent the standard deviation calculated from three independent experiments.

## Discussion

Fluorination plays a critical role in modern society, as organofluorines are widely used in pharmaceuticals, agrochemicals, and materials science^43–45^. The discovery of fluorinases—catalytically inefficient enzymes capable of catalyzing carbon–fluorine bond formation—revealed Nature’s strategy for selective biofluorination^21^. Fluorinated natural products are extremely rare in biology because fluoride is scarce and chemically inert in most environments^20,46^. Metabolic engineering and biocatalysis efforts^47–49^, while impressive, continue to be hampered by the limited activity of fluorinase. Recently, Li and co-workers established an asymmetric photoenzymatic approach for introducing fluorinated motifs into olefins, advancing enzymatic methods for the biocatalytic valorization of fluorinated feedstocks^50^. Previous efforts to engineer fluorinases have demonstrated success in broadening substrate specificity^38,42^, enhancing catalytic efficiency^27^, and modifying the oligomerization state^39^. Feng *et al.* employed site-directed and saturation mutagenesis around the active site, generating 85 mutants and achieving a 2-fold increase in catalytic efficiency. These results highlight the challenges of fluorinase engineering and underscore the need for further optimization to enable their sustainable use in synthesizing valuable organofluorines.

Understanding a reaction mechanism and its rate-limiting steps provides an excellent starting point for enzyme engineering^51^. To decipher the kinetic mechanism of fluorinase, we undertook a comprehensive analysis that involved collecting both enzymatic kinetics and oligomerization data. This dual focus was essential, as fluorinase activity is closely linked to its oligomeric state, and highly dynamic changes in oligomerization affect catalytic function^39^. A challenge was the development of a mathematical framework capable of simultaneously fitting the kinetic and oligomerization datasets. This task is non-trivial, due to the varying catalytic activities of distinct oligomeric species and the rapid, often transient, shifts in oligomerization equilibria that occur over time. We created an integrative mathematical model that captures the interplay between enzymatic activity and oligomerization dynamics (**Fig. 5**). The resulting methodology offers a robust and generalizable approach for analyzing the kinetics of multimeric enzymes. Beyond fluorinase, this framework is broadly applicable and can be adapted to the study of other oligomeric proteins, such as aldolase^52^ triosephosphate isomerase^53^, glutamate dehydrogenase^53^, (*S*)-carbonyl reductase^54^, catalase^55^, and vanillyl-alcohol oxidase^55^. We anticipate that the routine inclusion of oligomerization data will become a standard aspect of enzymology research by the increasing availability of advanced biophysical techniques, e.g., mass photometry, analytical ultracentrifugation, native mass spectrometry, and fluorescence correlation spectroscopy, which enable a real-time characterization of protein complexes.

Previous mechanistic studies discovered the key role of electronic contributions in the reaction mechanism of fluorinase^56^, and recognized fluoride desolvation, 5’-FDA release, and inhibition by SAM as potential bottlenecks in the mechanism^37^, but a kinetic characterization has not been reported to date. Through rational engineering of the fluorinase aggregation propensity, yielding the C44R variant, we were able to obtain sufficient amounts of soluble protein for transient kinetic experiments, which revealed multiple bottlenecks limiting its catalytic performance. The enzyme has a very low affinity for fluoride (*K*_d,F-_ = 151 mM), requiring high substrate concentrations that are incompatible with microbial systems due to toxicity^57,58^. The chemical step of carbon–fluorine bond formation is relatively fast; the rate-limiting step is the slow release of the product 5’-FDA. Two inhibitory off-pathways were identified: (i) blocking fluoride entry by competitive binding of SAM to the free enzyme, consistent with previous findings^37^, and (ii) a newly described unproductive binding of fluoride to the enzyme–product complex E·5’-FDA. Molecular dynamic simulations provided detailed insights into fluoride desolvation, enzymatic catalysis and assembly of oligomers (**Movies S1-S3**). Leveraging this knowledge, we engineered the double point mutant W50F+A279R, which preferentially formed hexamers and displayed enhanced catalytic efficiency in this oligomeric state. Additionally, the effect of the W50F mutation was largely preserved in the double mutant W50F+A279R, ensuring mitigation of the substrate inhibition by SAM. Kinetic characterization of the engineered mutants, combined with our mathematical framework, enabled the prediction of optimal reaction conditions, namely, 500 mM NaF and 0.5 mM SAM, with MtnN to remove 5’-FDA. Under these conditions, biofluorination achieved an apparent turnover number of 12.5 ± 2.1 min^−1^, corresponding to an approximately 60-fold improvement over previously reported values^22,26,39,57^.

The limited catalytic performance historically attributed to fluorinase arises largely from *in vitro* conditions that deviate from the natural biochemical context where the enzyme has evolved^22,26,39,57^. Under standard assay setups, elevated concentrations of fluoride and SAM promote the accumulation of unproductive enzyme–substrate complexes and reveal two kinetically trapped pathways, rendering most enzyme molecules inactive. By contrast, in natural settings, low fluoride availability restricts turnover but minimizes inhibitory interactions^59,60^. Our kinetic modeling and structural analysis support this interpretation, showing that rational mutations combined with optimized medium composition can restore substantial activity. Importantly, the enzyme evolved under environmental fluoride concentrations far below those required to achieve high turnover^59^. This evolutionary constraint shaped a catalytic mechanism optimized for selectivity and control, not for maximal catalytic constant *k_cat_*, which becomes detrimental under process conditions. These findings underscore the importance of matching assay design to physiological parameters in enzyme characterization and engineering.

In summary, our work represents a substantial advance in enzymatic fluorination by explaining long-standing limitations related to extremely low catalytic efficiency. Through the integration of mechanistic insights, transient kinetics, structural modelling, and targeted mutagenesis, we have not only engineered a functionally enhanced fluorinase but also optimized process conditions for efficient *de novo* biofluorination. Importantly, we have also established a generalizable strategy for optimizing the biocatalysis of oligomeric enzymes. A central element of this strategy has been a mathematical framework that offers a broadly applicable tool for the rigorous analysis and design of oligomeric enzyme systems ^52–55^.

## Acknowledgements

The authors acknowledge the support of the RECETOX Research Infrastructure (No. LM2023069) and CZECRIN (No. LM2023049), funded by the Ministry of Education, Youth and Sports (MEYS) of the Czech Republic. This project was also supported by the European Union’s Horizon 2020 Research and Innovation Programme under grant agreements CETOCOEN (No. 857560) and CLARA (No. 101136607). Additional funding was provided through the ESIF-MEYS Johannes Amos Comenius Programme under the CLARA project (No. CZ.02.01.01/00/23_029/0008437), co-financed by the European Union and MEYS. The project also received support from the National Institute for Cancer Research (EXCELES, No. LX22NPO5102), funded by the European Union – Next Generation EU. ZP acknowledges support from the Czech-Swiss grant provided by the Czech Science Foundation (No. 25-15784L). MM acknowledges support from the Czech Science Foundation (No. GX25-17329X). PIN gratefully acknowledges financial support from The Novo Nordisk Foundation through grants NNF20CC0035580, *LiFe* (NNF18OC0034818), *TARGET* (NNF21OC0067996), FM·*Pseudomonas* (NNF24OC0091501), and NovoF (NNF23OC0083631), and the European Union’s Horizon2020 Research and Innovation Programme under grant agreements No. 814418 (*SinFonia*) and No. 101082049 (*TOLERATE*).

## Author contributions

MS conducted aggregation studies and kinetic experiments, constructed enzyme mutants, prepared graphical materials, and contributed to manuscript writing; DCV characterized enzyme kinetics, acquired experimental data, purified and isolated enzyme variants, and designed production media for optimized 5’-FDA synthesis; IP constructed enzyme mutants and characterized fluorinase activity *in vitro*; AK determined the oligomeric states of enzyme variants; NBK designed and conducted the majority of metadynamics and steered molecular dynamics simulations, enriched the mutation dataset, interpreted results, and contributed to manuscript writing; AJS designed and performed metadynamics studies, significantly contributed to the experimental design of computational work, and participated in manuscript writing; LM designed and conducted the majority of QM/MM studies, interpreted the results, and contributed to manuscript writing; GS performed the majority of 7D grid-based AI experiments for mutation design and reviewed the manuscript; RL conducted 7D grid-based AI experiments for mutation design and participated in manuscript review; AKB supervised the determination of oligomeric states; MM supervised molecular biology experiments and contributed to the development of non-aggregating mutants; JD designed enzyme mutants, interpreted experimental results, and contributed to manuscript writing; PRK designed and interpreted the 7D grid-based AI experiments and conducted QMD studies, was involved in the overall design of computational experiments, performed computational analyses, interpreted results, contributed to manuscript writing, and reviewed the final version; PIN conducted data analysis, interpreted results, contributed to the manuscript, coordinated experimental efforts, and managed the EU project that provided essential funding; ZP performed kinetic analyses, interpreted results, contributed to manuscript writing, and coordinated experiments.

## Supplementary Materials

Materials and Methods

Supplementary Text

Figs. S1 to S31

Tables S1 to S4

## References and Notes

1. Furuya, T., Kamlet, A. S. & Ritter, T. Catalysis for fluorination and trifluoromethylation. Nature 473, 470–477 (2011).

2. Müller, K., Faeh, C. & Diederich, F. Fluorine in Pharmaceuticals: Looking Beyond Intuition. Science 317, 1881–1886 (2007).

3. Research, C. for D. E. and. Novel Drug Approvals at FDA. FDA https://www.fda.gov/drugs/development-approval-process-drugs/novel-drug-approvals-fda (2024).

4. Ali, S. & Zhou, J. Highlights on U.S. FDA-approved fluorinated drugs over the past five years (2018–2022). European Journal of Medicinal Chemistry 256, 115476 (2023).

5. Mei, H., et al. Fluorine-Containing Drugs Approved by the FDA in 2018. Chemistry – A European Journal 25, 11797–11819 (2019).

6. The Bestselling Drug Products of 2023: Ranking All 152 Blockbusters | Solt DB. https://www.living.tech/data-visual/the-bestselling-drug-products-of-2023-ranking-all-152-blockbusters.

7. Li, J. J. Discovery of Lipitor. in Triumph of the Heart: The Story of Statins (ed. Li, J. J.) 0 (Oxford University Press, 2009). doi:10.1093/oso/9780195323573.003.0010.

8. Wang, J., et al. Fluorine in Pharmaceutical Industry: Fluorine-Containing Drugs Introduced to the Market in the Last Decade (2001–2011). Chem. Rev. 114, 2432–2506 (2014).

9. Jeschke, P. Current Trends in the Design of Fluorine-Containing Agrochemicals. in Organofluorine Chemistry 363–395 (2021). doi:10.1002/9783527825158.ch11.

10. Compendium of Pesticide Common Names. http://www.bcpcpesticidecompendium.org/.

11. Martinelli, L. & Nikel, P. I. Breaking the state-of-the-art in the chemical industry with new-to-Nature products via synthetic microbiology. Microb Biotechnol 12, 187–190 (2019).

12. Plunkett, R. J. Tetrafluoroethylene polymers. (1941).

13. Harsanyi, A. & Sandford, G. Organofluorine chemistry: applications, sources and sustainability. Green Chem. 17, 2081–2086 (2015).

14. O’Hagan, D. & B. Harper, D. Fluorine-containing natural products. Journal of Fluorine Chemistry 100, 127–133 (1999).

15. Chan, K. K. J. & O’Hagan, D. The Rare Fluorinated Natural Products and Biotechnological Prospects for Fluorine Enzymology. in NATURAL PRODUCT BIOSYNTHESIS BY MICROORGANISMS AND PLANTS, PT B (ed. Hopwood, D.) 219–235 (Academic Press/Elsevier, SAN DIEGO, 2012). doi:10.1016/B978-0-12-394291-3.00003-4.

16. Ludewig, H. et al. Halogenases: structures and functions. Current Opinion in Structural Biology 65, 51–60 (2020).

17. Gribble, G. W. A recent survey of naturally occurring organohalogen compounds. Environ. Chem. 12, 396–405 (2015).

18. O’Hagan, D. & Deng, H. Enzymatic Fluorination and Biotechnological Developments of the Fluorinase. Chem. Rev. 115, 634–649 (2015).

19. Neumann, C. S., Fujimori, D. G. & Walsh, C. T. Halogenation strategies in natural product biosynthesis. Chem Biol 15, 99–109 (2008).

20. Dong, C., et al. Crystal structure and mechanism of a bacterial fluorinating enzyme. Nature 427, 561–565 (2004).

21. O’Hagan, D., Schaffrath, C., Cobb, S. L., Hamilton, J. T. G. & Murphy, C. D. Biosynthesis of an organofluorine molecule. Nature 416, 279–279 (2002).

22. Deng, H., et al. Identification of Fluorinases from Streptomyces sp MA37, Norcardia brasiliensis, and Actinoplanes sp N902-109 by Genome Mining. ChemBioChem 15, 364–368 (2014).

23. Wang, Y., Deng, Z. & Qu, X. Characterization of a SAM-dependent fluorinase from a latent biosynthetic pathway for fluoroacetate and 4-fluorothreonine formation in Nocardia brasiliensis. F1000Res 3, 61 (2014).

24. Huang, S., et al. Fluoroacetate biosynthesis from the marine-derived bacterium Streptomyces xinghaiensis NRRL B-24674. Org. Biomol. Chem. 12, 4828–4831 (2014).

25. Sooklal, S. A., De Koning, C., Brady, D. & Rumbold, K. Identification and characterisation of a fluorinase from Actinopolyspora mzabensis. Protein Expr Purif 166, 105508 (2020).

26. Pardo, I., et al. A Nonconventional Archaeal Fluorinase Identified by In Silico Mining for Enhanced Fluorine Biocatalysis. ACS Catal. 12, 6570–6577 (2022).

27. Feng, X., Cao, Y., Liu, W. & Xian, M. Identification of Two Novel Fluorinases From Amycolatopsis sp. CA-128772 and Methanosaeta sp. PtaU1.Bin055 and a Mutant With Improved Catalytic Efficiency With Native Substrate. Front. Bioeng. Biotechnol. 10, (2022).

28. Haas, R. & Nikel, P. I. Challenges and opportunities in bringing nonbiological atoms to life with synthetic metabolism. Trends Biotechnol 41, 27–45 (2023).

29. Cros, A., Alfaro-Espinoza, G., De Maria, A., Wirth, N. T. & Nikel, P. I. Synthetic metabolism for biohalogenation. Curr Opin Biotechnol 74, 180–193 (2022).

30. Calero, P., et al. A fluoride-responsive genetic circuit enables in vivo biofluorination in engineered Pseudomonas putida. Nat Commun 11, 5045 (2020).

31. Salvatore, D., Croguennec, T., Bouhallab, S., Forge, V. & Nicolai, T. Kinetics and Structure during Self-Assembly of Oppositely Charged Proteins in Aqueous Solution. Biomacromolecules 12, 1920–1926 (2011).

32. Sumbalova, L., Stourac, J., Martinek, T., Bednar, D. & Damborsky, J. HotSpot Wizard 3.0: web server for automated design of mutations and smart libraries based on sequence input information. Nucleic Acids Res 46, W356–W362 (2018).

33. Nolte, C., Ammer, J. & Mayr, H. Nucleofugality and Nucleophilicity of Fluoride in Protic Solvents. J. Org. Chem. 77, 3325–3335 (2012).

34. Ma, H., et al. Effects of fluoride on bacterial growth and its gene/protein expression. Chemosphere 100, 190–193 (2014).

35. Fuge, R. & Andrews, M. J. Fluorine in the UK environment. Environ Geochem Health 10, 96–104 (1988).

36. Mikkonen, H. G., et al. Environmental and anthropogenic influences on ambient background concentrations of fluoride in soil. Environmental Pollution 242, 1838–1849 (2018).

37. Zhu, X., Robinson, D. A., McEwan, A. R., O’Hagan, D. & Naismith, J. H. Mechanism of Enzymatic Fluorination in Streptomyces cattleya. J. Am. Chem. Soc. 129, 14597–14604 (2007).

38. Sun, H., et al. Directed Evolution of a Fluorinase for Improved Fluorination Efficiency with a Non-native Substrate. Angewandte Chemie International Edition 55, 14277–14280 (2016).

39. Kittilä, T., et al. Oligomerization engineering of the fluorinase enzyme leads to an active trimer that supports synthesis of fluorometabolites in vitro. Microb Biotechnol 15, 1622–1632 (2022).

40. Parker, W. B. Enzymology of Purine and Pyrimidine Antimetabolites Used in the Treatment of Cancer. Chem. Rev. 109, 2880–2893 (2009).

41. Onega, M., et al. An enzymatic route to 5-deoxy-5-[18F]fluoro-D-ribose, a [18F]-fluorinated sugar for PET imaging. Chem. Commun. 46, 139–141 (2010).

42. Yeo, W. L., et al. Probing the molecular determinants of fluorinase specificity. Chem. Commun. 53, 2559–2562 (2017).

43. O’Hagan, D. Understanding organofluorine chemistry. An introduction to the C–F bond. Chem. Soc. Rev. 37, 308–319 (2008).

44. Ogawa, Y., Tokunaga, E., Kobayashi, O., Hirai, K. & Shibata, N. Current Contributions of Organofluorine Compounds to the Agrochemical Industry. iScience 23, 101467 (2020).

45. Inoue, M., Sumii, Y. & Shibata, N. Contribution of Organofluorine Compounds to Pharmaceuticals. ACS Omega 5, 10633–10640 (2020).

46. Carvalho, M. F. & Oliveira, R. S. Natural production of fluorinated compounds and biotechnological prospects of the fluorinase enzyme. Critical Reviews in Biotechnology 37, 880–897 (2017).

47. Sirirungruang, S., et al. Engineering site-selective incorporation of fluorine into polyketides. Nat Chem Biol 18, 886–893 (2022).

48. Walker, M. C. & Chang, M. C. Y. Natural and engineered biosynthesis of fluorinated natural products. Chem Soc Rev 43, 6527–6536 (2014).

49. Thuronyi, B. W., Privalsky, T. M. & Chang, M. C. Y. Engineered Fluorine Metabolism and Fluoropolymer Production in Living Cells. Angew Chem Int Ed Engl 56, 13637–13640 (2017).

50. Li, M., Yuan, Y., Harrison, W., Zhang, Z. & Zhao, H. Asymmetric photoenzymatic incorporation of fluorinated motifs into olefins. Science 385, 416–421 (2024).

51. Bornscheuer, U. T., et al. Engineering the third wave of biocatalysis. Nature 485, 185–194 (2012).

52. Katebi, A. R. & Jernigan, R. L. Aldolases Utilize Different Oligomeric States To Preserve Their Functional Dynamics. Biochemistry 54, 3543–3554 (2015).

53. Mishra, S. K., Sankar, K. & Jernigan, R. L. Altered Dynamics upon Oligomerization Corresponds to Key Functional Sites. Proteins 85, 1422–1434 (2017).

54. Li, Y., et al. Oligomeric interactions maintain active-site structure in a noncooperative enzyme family. The EMBO Journal 41, e108368 (2022).

55. Yancheva, Y. D., Kaya, S. G., Belyy, A., Fraaije, M. W. & Tych, K. M. Impact of Ligand-Induced Oligomer Dissociation on Enzyme Diffusion, Directly Observed at the Single-Molecule Level. Nano Lett. 25, 2373–2380 (2025).

56. Gutiérrez-Sánchez, N., Kästner, J., Mendizábal, F. & Miranda-Rojas, S. Insights into the Catalytic Mechanism and Selectivity of S-Adenosyl Methionine (SAM)-Dependent Fluorinase toward Carbon–Halogen Bond Formation. J. Chem. Inf. Model. 65, 6998–7012 (2025).

57. Ma, L., et al. Biological fluorination from the sea: discovery of a SAM-dependent nucleophilic fluorinating enzyme from the marine-derived bacterium Streptomyces xinghaiensis NRRL B24674. RSC Advances 6, 27047–27051 (2016).

58. Calero, P., Gurdo, N. & Nikel, P. I. Role of the CrcB transporter of Pseudomonas putida in the multi-level stress response elicited by mineral fluoride. Environmental Microbiology 24, 5082–5104 (2022).

59. Carvalho, M. F. & Oliveira, R. S. Natural production of fluorinated compounds and biotechnological prospects of the fluorinase enzyme. Crit Rev Biotechnol 37, 880–897 (2017).

60. Petkowski, J. J., Seager, S. & Bains, W. Reasons why life on Earth rarely makes fluorine-containing compounds and their implications for the search for life beyond Earth. Sci Rep 14, 15575 (2024).

61. Johnson, K. A., Simpson, Z. B. & Blom, T. Global kinetic explorer: a new computer program for dynamic simulation and fitting of kinetic data. Anal Biochem 387, 20–29 (2009).

62. Gasteiger, E., et al. Protein Identification and Analysis Tools on the ExPASy Server. in The Proteomics Protocols Handbook (ed. Walker, J. M.) 571–607 (Humana Press, Totowa, NJ, 2005). doi:10.1385/1-59259-890-0:571.

63. Forloni, M., Liu, A. Y. & Wajapeyee, N. Megaprimer Polymerase Chain Reaction (PCR)-Based Mutagenesis. Cold Spring Harb Protoc 2019, pdb.prot097824 (2019).

64. Johnson, K. A., Simpson, Z. B. & Blom, T. FitSpace explorer: an algorithm to evaluate multidimensional parameter space in fitting kinetic data. Anal Biochem 387, 30–41 (2009).

65. Miao, Y., Feher, V. A. & McCammon, J. A. Gaussian Accelerated Molecular Dynamics: Unconstrained Enhanced Sampling and Free Energy Calculation. J. Chem. Theory Comput. 11, 3584–3595 (2015).

66. Pang, Y. T., Miao, Y., Wang, Y. & McCammon, J. A. Gaussian Accelerated Molecular Dynamics in NAMD. J Chem Theory Comput 13, 9–19 (2017).

67. Miao, Y. & McCammon, J. A. Gaussian Accelerated Molecular Dynamics: Theory, Implementation, and Applications. Annu Rep Comput Chem 13, 231–278 (2017).

68. Miao, Y., et al. Improved Reweighting of Accelerated Molecular Dynamics Simulations for Free Energy Calculation. J. Chem. Theory Comput. 10, 2677–2689 (2014).

69. Vanommeslaeghe, K., Raman, E. P. & MacKerell, A. D. Jr. Automation of the CHARMM General Force Field (CGenFF) II: Assignment of Bonded Parameters and Partial Atomic Charges. J. Chem. Inf. Model. 52, 3155–3168 (2012).

70. Y, A., Mp, M., O, I. & Dg, F. GAMESS as a free quantum-mechanical platform for drug research. Current topics in medicinal chemistry 12, (2012).

71. W, H., A, D. & K, S. VMD: visual molecular dynamics. Journal of molecular graphics 14, (1996).

72. Gz, G., et al. Mechanical signaling on the single protein level studied using steered molecular dynamics. Cell biochemistry and biophysics 55, (2009).

73. Skovstrup, S., David, L., Taboureau, O. & Jørgensen, F. S. A Steered Molecular Dynamics Study of Binding and Translocation Processes in the GABA Transporter. PLOS ONE 7, e39360 (2012).

74. Shen, M., et al. Steered molecular dynamics simulations on the binding of the appendant structure and helix-β2 in domain-swapped human cystatin C dimer. J Biomol Struct Dyn 30, 652–661 (2012).

75. Kumari, J. L. J., Sudan, R. J. J. & Sudandiradoss, C. Evaluation of peptide designing strategy against subunit reassociation in mucin 1: A steered molecular dynamics approach. PLOS ONE 12, e0183041 (2017).

76. Londhe, A. M., Gadhe, C. G., Lim, S. M. & Pae, A. N. Investigation of Molecular Details of Keap1-Nrf2 Inhibitors Using Molecular Dynamics and Umbrella Sampling Techniques. Molecules 24, 4085 (2019).

77. Bowman, J. D. & Lindert, S. Molecular Dynamics and Umbrella Sampling Simulations Elucidate Differences in Troponin C Isoform and Mutant Hydrophobic Patch Exposure. J Phys Chem B 122, 7874–7883 (2018).

78. Lindorff-Larsen, K., et al. Improved side-chain torsion potentials for the Amber ff99SB protein force field. Proteins 78, 1950–1958 (2010).

79. Abraham, M. J. & Gready, J. E. Optimization of parameters for molecular dynamics simulation using smooth particle-mesh Ewald in GROMACS 4.5. J Comput Chem 32, 2031–2040 (2011).

80. Pronk, S., et al. GROMACS 4.5: a high-throughput and highly parallel open source molecular simulation toolkit. Bioinformatics 29, 845–854 (2013).

81. Wang, Y.-L., Zhu, Y.-L., Lu, Z.-Y. & Laaksonen, A. Electrostatic interactions in soft particle systems: mesoscale simulations of ionic liquids. Soft Matter 14, 4252–4267 (2018).

82. Hess, B., Bekker, H., Berendsen, H. J. C. & Fraaije, J. G. E. M. LINCS: A linear constraint solver for molecular simulations. Journal of Computational Chemistry 18, 1463–1472 (1997).

83. Hsu, W.-T., Piomponi, V., Merz, P. T., Bussi, G. & Shirts, M. R. Alchemical Metadynamics: Adding Alchemical Variables to Metadynamics to Enhance Sampling in Free Energy Calculations. J Chem Theory Comput 19, 1805–1817 (2023).

84. Nava, M. Implementing dimer metadynamics using gromacs. J Comput Chem 39, 2126–2132 (2018).

85. Biswas, S. & Wong, B. Ab Initio Metadynamics Calculations Reveal Complex Interfacial Effects in Acetic Acid Deprotonation Dynamics. Preprint at 10.26434/chemrxiv.13647920.v1 (2021).

86. Sucerquia, D., Parra, C., Cossio, P. & Lopez-Acevedo, O. Ab initio metadynamics determination of temperature-dependent free-energy landscape in ultrasmall silver clusters. The Journal of Chemical Physics 156, 154301 (2022).

87. R, P. K. & L, R. A novel 7D-QSAR approach, combining QM based grid and solvation models to predict hotspots and kinetic properties of mutated enzymes: an enzyme engineering perspective. Insights in Enzyme Research https://doi.org/10.21767/2573-4466-C1-002 (2018) doi:10.21767/2573-4466-C1-002.

88. Kumar, P., Roopa L, K. & Sigamani, G. 7D QSAR based grid maps generated using quantum mechanic probes to identify hotspots and predict activity of mutated enzymes for enzyme engineering. Enzyme Engineering XXV https://dc.engconfintl.org/enzyme_xxv/127 (2019).

89. Sigamani, G., Kumar, P. & L, R. 7d-grid-ai technology: A technology that translates enzymes from a computer to business with limited lab experiments. Enzyme Engineering XXVII https://dc.engconfintl.org/enzyme_xxvii/50 (2023).

90. Melo, M. C. R., et al. NAMD goes quantum: an integrative suite for hybrid simulations. Nat Methods 15, 351–354 (2018).

91. Laio, A. & Parrinello, M. Escaping free-energy minima. Proceedings of the National Academy of Sciences 99, 12562–12566 (2002).

92. Valsson, O., Tiwary, P. & Parrinello, M. Enhancing Important Fluctuations: Rare Events and Metadynamics from a Conceptual Viewpoint. Annu Rev Phys Chem 67, 159–184 (2016).

93. Fiorin, G., Klein, M. L. & Hénin, J. Using collective variables to drive molecular dynamics simulations. Molecular Physics 111, 3345–3362 (2013).

94. Hostaš, J., Řezáč, J. & Hobza, P. On the performance of the semiempirical quantum mechanical PM6 and PM7 methods for noncovalent interactions. Chemical Physics Letters 568-569, 161–166 (2013).

95. Geu-Flores, F., Nour-Eldin, H. H., Nielsen, M. T. & Halkier, B. A. USER fusion: a rapid and efficient method for simultaneous fusion and cloning of multiple PCR products. Nucleic Acids Res 35, e55 (2007).

