## Supplementary Information for "Mechanism-Guided Engineering of Fluorinase Unlocks Efficient Nucleophilic Biofluorination"

**The PDF File includes:**

Materials and Methods
Supplementary Text
Figs. S1 to S31
Tables S1 to S4

### Materials and Methods

#### S1 Fluorinase expression and purification

pET28a(+)::*flA1_6xHis* plasmid encoding fluorinase from *Streptomyces* sp. MA37 with an N-terminal 6-histidine tag and a kanamycin resistance marker was transformed into *E.coli* BL21(DE3). The cells were grown in LB medium with 50 $\mu$g ml$^{-1}$ kanamycin at 37 °C and 115 rpm. Once they reached the exponential growth phase ($OD_{600}$ = 0.4-0.6), 0.5 mM IPTG was added to induce overexpression of FlA1, and the cells were grown at 20 °C and 115 rpm overnight. The cells were harvested by centrifugation (4,000 g, 15 mins, 4 °C), resuspended in purification buffer A (16 mM $K_2HPO_4$, 3.6 mM $KH_2PO_4$, 500 mM NaCl, 10 mM imidazole, pH 7.5), and frozen at -70 °C. DNase I (2 $\mu$g ml$^{-1}$) was added to the defrosted cells, which were then disrupted by sonication (sonic dismembrator Fisherbrand™ 705, Fisher Scientific, USA) and centrifuged (21,000 g, 1 h, 4 °C). The cell-free extract was loaded on Ni-NTA Superflow Cartridge loaded with $Ni^{2+}$ ions (Quiagen, Germany) in purification buffer A as the mobile phase. Metal affinity chromatography based on the strong interaction between the His-tag and $Ni^{2+}$ ions was used to purify the protein. The target enzyme was eluted when a 60 % gradient of purification buffer B (16 mM $K_2HPO_4$, 3.6 mM $KH_2PO_4$, 500 mM NaCl, 500 mM imidazole, pH 7.5) was reached. Collected fractions were transferred to 30 mM HEPES buffer (pH 7.8) with added salt (KF, NaF or NaCl of variable concentration) using PD-10 Desalting Columns (Cytiva, USA). SDS-PAGE (120 V, 80 mins) was used to verify the expression of fluorinase in the soluble and insoluble fraction of *E. coli* and the purity of the isolated protein. Enzyme concentration was determined using spectrophotometer DeNovix DS-11 by measuring absorbance at 280 nm with a known extinction coefficient calculated from the amino acid sequence using the ProtParam tool by ExPASy[62].

#### S2 SEC analysis of oligomeric states

The oligomeric state of purified proteins was analyzed by size-exclusion chromatography (SEC) using the ÄKTA FPLC system with Superdex 200 10/300 GL column. In the case of the wild-type FlA1, the system was equilibrated with 30 mM HEPES buffer (pH 7.8) with 75 mM KF. To characterize the cysteine mutants FlA1+C35A, FlA1+C44T and FlA1+C35A+C44T, 30 mM HEPES (pH 7.8) with 400 mM KF and 10 mM dithiothreitol (DTT) was used. 200 µl of the protein sample (~1 mg/ml) was injected onto the column and separated at a constant flow rate of 0.5 ml min$^{-1}$. The molecular weight was determined using low and high molecular weight calibration kits (GE Healthcare, Sweden).

#### S3 Temperature denaturation and aggregation studies

Uncle (Unchained Labs) was used to measure static light scattering (SLS) at 266 nm and barycentric mean fluorescence (BCM) of samples excited at 266 nm. The scattered light was detected at a 90° angle, assuming the same intensity in all directions (Rayleigh scattering) if the detected particles are small relative to the incident light. The signal was followed either in time during a 10-hour incubation at set temperatures (25 °C, 30 °C, 37 °C) or over a range of temperatures during heating of the sample from 15 °C to 95 °C at a 1 °C min$^{-1}$ scan rate. FlA1 (0.138-2.2 mg ml$^{-1}$) in 30 mM HEPES (pH 7.8) and 75 mM KF, with or without added DTT (0 – 20 mM), was used for the measurements.

S4 Site-directed mutagenesis (C35A, C44T, C35A+C44T)

Cysteine residues in FlA1 were mutated using the Megaprimer PCR-based mutagenesis method[63] with primers 5’-aagggcctgatgcacagcatcgcccccggtgtgaccgtggtggat-3’ for FlA1+C35A, 5’-accgtggtggatgtgacccacagc atgaccccgtgggac-3’ for FlA1+C44T, 5’-aagggcctgatgcacagcatcgcccccggtgtgaccgtggtggatgtgacccacagcatgacccc gtgggac-3’ for FlA1+C35A+C44T and 5’-gctagttattgctcagcgg-3’ as the reverse primer. Subsequently, the methodology described in S1 was used to express and purify the fluorinase variants.

S5 Steady-state kinetics of FlA1+C44T

The forward (fluorination) reactions were measured with 12 $\mu$M FlA1+C44T and a set of *S*-adenosylmethionine (SAM) (200, 400, 800 $\mu$M) and KF (25, 100, 400, 800 mM) concentrations (**Figures S5a-d**). To analyze substrate inhibition, the same experiments were carried out after 30-min preincubation of fluorinase with SAM (**Figure S5k**). To determine the contribution of the reverse reaction (defluorination) to the data, L-amino acid oxidase was added to the reactions to the final activity 2.13 mmol $l^{-1}$ $min^{-1}$ to block defluorination by consuming L-methionine (Met) (**Figures S5g-j**). The reverse reaction was measured with 25 $\mu$M FlA1 and a set of 5’-fluoro-5’-deoxyadenosine (5’-FDA) (400, 800, 1600 $\mu$M) and Met (800, 1600, 3200, 5000 $\mu$M) concentrations (**Figures S5e-f**). In all steady-state kinetic datasets, the samples were incubated for over 6 hours with shaking at 30 °C in 30 mM HEPES and 150 mM NaCl (pH 7.8). 50 µl samples were taken out of the reaction mixtures in 20-30 min intervals and mixed with 25 µl of 1.6 M sulphuric acid to terminate the reaction. Subsequently, 25 µl of 3.2 M sodium hydroxide was added to neutralize the samples, and the denatured enzyme was removed by centrifugation at 14,000 g and 4 °C for 3 mins. 80 µl of the supernatants were transferred to glass vials for quantification. RP-HPLC method using the Eclipse Plus C18 HPLC column was developed to detect and quantify SAM and 5’-FDA. The mobile phase consisted of 0.05% acetic acid with 5% acetonitrile (ACN), and elution was carried out by gradually increasing the ACN concentration. Absorbance at 260 nm was detected. Calibration standards were prepared in the same manner as the analysed samples by mixing with sulphuric acid and sodium hydroxide.

S6 Pre-steady-state burst of FlA1+C44T

46-58 µM FlA1+C44T and 800 µM SAM solutions were used for the quenched-flow measurements using the QFM rapid quench-flow instrument (Bio-Logic, France). For the experiment without preincubation, 800 mM KF was present in the SAM solution only (**Figure S5l**). In the setup where fluorinase was pre-incubated with fluoride, 400 mM KF was present in both the enzyme and substrate solution (**Figure S5m**). The reactions proceeded at 30 °C and pH 7.8 in 30 mM HEPES with 150 mM NaCl. 75 µl of enzyme and 75 µl of substrate were rapidly mixed to achieve final concentrations of 400 µM SAM, 400 mM KF, and 23-29 µM FlA1+C44T. Incubation of the mixture for variable reaction times (10 ms – 10 s) was followed by mixing with 100 µl of quencher (1.6 M sulphuric acid). 100 µl of 3.2 M sodium hydroxide were added to the collected samples for neutralization, and the denatured enzyme was removed by centrifugation at 14,000 g and 4 °C for 3 mins. 150 µl of the supernatants were transferred to glass vials, and the amount of 5’-FDA formed was quantified using RP-HPLC as described above.

### S7 Data analysis and statistics (FlA1+C44T kinetic mechanism)

All steady-state and pre-steady-state kinetic data were fit globally using the KinTek Explorer (KinTek, USA)[61] to pre-defined reaction models. During the data fitting, the program applied numerical integration of rate equations from the input models to fit the experimental data with simulated curves. This was achieved by searching a set of parameters using the Bulirsch–Stoer algorithm with an adaptive step size that produces a minimum $\chi^2$ value calculated using nonlinear regression based on the Levenberg-Marquardt method. To account for slight variations in the data, enzyme or substrate concentrations were allowed to vary within the interval of 10 % to make the best fits possible. A rigorous analysis of the variation of the kinetic parameters was accomplished by confidence contour analysis with FitSpace Explorer[64]. Each derived kinetic parameter was held fixed at varying values while the other constants were allowed to float to achieve the minimal $\chi^2$ value. An increase in $\chi^2$ as the fixed parameter diverges from the optimal value obtained from the fit implies that this parameter is well-defined. Setting a threshold of acceptable $\chi^2$ values ($\min\chi^2/\chi^2$ = 0.98) enabled the determination of a lower and upper bounds of each parameter.

Simple reaction models, such as the 5-step reaction mechanism of fluorinase described in **Figure 1c**, failed to yield satisfactory statistical fits. We explored a number of more complex reaction models, and finally, the model visualized in **Figure S6** could provide well-defined kinetic constants (**Table S2, Figure S7**).

### S8 DLS analysis of oligomeric states

Uncle (Unchained Labs) was used to measure dynamic light scattering (DLS) to determine the oligomeric state of FlA1+C44T during kinetic experiments. The enzyme of two different concentrations (29.7 $\mu$M and 7.43 $\mu$M) was analyzed at different concentrations of KF (0 – 800 mM). The total concentration of salt was adjusted to be the same in all samples by adding NaBr ($c_{\mathrm{KF+NaBr}}$ = 850 mM).

### S9 Gaussian Accelerated Molecular Dynamics (GaMD) studies

GaMD is a comprehensive molecular simulation technique within an enhanced sampling regime that accelerates protein conformational transitions between low-energy states. It achieves this by smoothing the potential energy surface using a harmonic boost potential that follows a Gaussian distribution, thereby aiding in the study of $F^-$ diffusion.[65–67]

Consider a system with NNN atoms positioned at $\vec{r} = \{\vec{r}1 \ldots\ldots \vec{r}N\}$. When the total system potential $V(\vec{r})$ is lower than a reference energy $E$, the new potential $V^*(\vec{r})$ of the system is calculated as

$$V^*(\vec{r}) = V(\vec{r}) + \Delta V(\vec{r})$$
$$\Delta V(\vec{r}) = \{1/2\, k(E - V(\vec{r}))^2, V(\vec{r}) < E\ 0, \qquad V(\vec{r}) \geq E$$

where $k$ represents the harmonic force constant. The two adjustable parameters, $E$ and $k$ are automatically set according to three principles of enhanced sampling.[68]

First, for any two arbitrary potential values $V_1(\vec{r})$ and $V_2(\vec{r})$ on the original energy surface, if $V_1(\vec{r}) < V_2(\vec{r})$, $\Delta V$ should be a monotonic function that that preserves the relative order of the biased potential values; i.e., ${V_1}^*(\vec{r}) < {V_2}^*(\vec{r})$. Second, if $V_1(\vec{r}) < V_2(\vec{r})$, the potential difference observed on the smoothed energy surface should be smaller than that of the original, in other words, ${V_2}^*(\vec{r}) - {V_1}^*(\vec{r}) < V_2(\vec{r}) - V_1(\vec{r})$. By combining these two criteria and applying them to the formula for $V^*(\vec{r})$ $and$ $\Delta V$ , we obtain:

$$V_{max} \leq E \leq V_{max} + 1/k$$

Where $V_{min}$ and $V_{max}$ represent the minimum and maximum potential energies of the system. For this equation to hold, $k$ must satisfy the condition $k \leq 1/(V_{max} - V_{min})$. By defining $k$ a $k \leq k_0/(V_{max} - V_{min})$, we ensure that $0 < k_0 \leq 1$. Third, the standard deviation (SD) of $\Delta V$ must be sufficiently small (indicating a narrow distribution) to enable accurate reweighting via cumulant expansion to the second order.

This is expressed as $\sigma_{\Delta v} = k\left(E - V_{avg}\right)\sigma_v \leq \sigma_0$ , where $V_{avg}$ and $\sigma_v$ are the average and SD of $\Delta V$, respectively, and $\sigma_0$ is a user-specified upper limit for accurate reweighting. When E is set to the lower bound $E = V_{max}$ according to equation 2, $k_0$ can be calculated as:

$$k_0 = min(1.0, k_0') = \left(1.0, \frac{\sigma_0}{\sigma_v} . \frac{V_{max} - V_{min}}{V_{max} - V_{avg}}\right)$$

Alternatively, when the threshold energy E is set to its upper bound $E = V_{max} + \frac{1}{k}$, $k_0$ is set to $k_0 = k_0'' \equiv \left(1 - \frac{\sigma_0}{\sigma_v}\right). \frac{V_{max} - V_{min}}{V_{avg} - V_{min}}$ if $k_0''$ is calculated between 0 and 1.

**System setup:** The 5'-FDA charges and molecular mechanics parameters were generated from cgenff server and SAM chargers were generated using GAMESS[69,70]. The *psfgen* plugin of VMD was used to generate psf files using CHARMM36 parameters[71]. The system was solvated and neutralised at 0.2 M NaF with the help of solvate plugin of VMD. The initial minimization, annealing and 20 ns of cMD equilibration of added boost potential were carried out at 310K and 1 atm pressure using NAMD2.13.

Particle-mesh Ewald summation method was used to compute electrostatic interactions with cut-off distance of 15 Å and a 2 fs integration time step was used. Different independent production GaMD simulations were run, lasting 1000 ns at the dual-boost and dihedral acceleration levels.

S10 Steered molecular Dynamics (SMD) studies

The SMD simulations were conducted to understand the entry and stabilization of $F^-$ ion in different paths when the $F^-$ ion was exposed to mechanical strain or rupture force, which cannot be achieved through standard MD simulations. The pulling simulations were implemented using GROMACS 2019.4 tool. Multiple paths leading to the active site residues (Ser158 and Thr80) from tunnel studies were considered. The pull velocity of 0.0005 $nm.ps^{-1}$ with the bias force constant of -8 $kJ\ mol^{-1}\ nm^{-2}$ was used. The SMD experimental protocol was maintained the same for E+$F^-$, E.SAM+$F^-$, and E-5'-FDA+$F^-$ where $F^-$ was constantly subjected to steered diffusion.[72–75]

Umbrella sampling (US) studies were conducted as an extension of SMD studies to estimate the energetics during the transfer of $F^-$ ion in the path towards active site. A series of reaction coordinates across the paths were chosen from the SMD studies and windows were constructed based on the distance between $F^-$ ion and catalytic residues. The path was discretized into multiple windows for every 0.5Å of the $F^-$ ion movement. SMD and umbrella sampling simulations were repeated 3 times for statistical significance.[76,77]

### S11 Metadynamics simulations for E·5'-FDA and F-· E·5'-FDA complexes

Molecular dynamics simulations were conducted on Apo protein using GROMACS 2019.4. The simulation was performed using Amber99sb force fields[78] with the water model TIP3P to mimic the natural environment around protein. TIP3P water model with the box volume of 1000 $nm^3$ was used to solvate the complex. The apoenzyme system was neutralized using 0.2 M concentration of $Na^+$ and $F^-$ ions. 3593 water molecules, 67 $Na^+$ ions, and 54 $F^-$ ions were added, which brings the overall number of atoms to 54653[79,80].

Energy minimization of all the systems was performed using the steepest descent algorithm with the convergence energy cut-off of 1000 kJ $mol^{-1}$ within 1000 steps. Equilibration was done using NVT and NPT ensembles. NVT was carried out at constant volume and temperature equilibrium and the NPT was carried out a constant pressure. The NVT and NPT equilibration steps were performed for 30 ns at 300 K. The short-range cut-off values of van der Waals were set to 1.0, and columbic interactions were calculated using PME (Particle Mesh Ewald) algorithm[79,81]. LINCS algorithm[82] was used for all bond length constraints and each time step of simulations was set to 2 fs.

Metadynamics simulations were conducted using plumed software coupled with GROMACS 2019.4. Well-equilibrated states of the enzyme-product complex in the presence and absence of $F^-$, taken from MD studies, were used for metadynamics-based studies. Collective variables for the enzyme-product complex were defined as CV1: Distance between the Center of Mass (COM) of adenine moiety of 5'-FDA ($FDA_{ade}$) and the COM of residues interacting with ($FDA_{ade}$) in Å. CV2: Distance between the COM of ribose-sugar moiety of 5'-FDA and the COM of residues Asp16 and Tyr77 in Å. Simulations were performed twice for 50 ns with a bias factor set to 8.0, the temperature set to 310 K, and the results were calculated accordingly. The free energy landscape was explored for the interaction of the substrate or product with the residues lining the active site[83–86].

### S12 Grid-Based Interaction Mapping of Molecular Probes with Enzyme-Substrate Ensembles for Predicting Mutation Hotspots and Substitutions

Enzyme-substrate conformations derived from both the diffusion pathway and the active-site structure preceding the entry of S-adenosylmethionine (SAM) were utilized in a grid-based mutation prediction protocol. Specifically, the conformational ensemble of the FlA1-$F^-$ complex within the active site, along with structures delineating the $F^-$ entry pathway, was extracted, aligned, and subsequently placed within a three-dimensional grid framework.

The grid was systematically constructed using grid points spaced at 2.0 Å resolution. At each grid point, molecular probes were iteratively positioned to simulate potential amino acid side chain interactions or solvent interactions. Probes representing amino acid side chains and water molecules were strategically aligned so that their geometric centers coincided precisely with each grid point, ensuring a standardized interaction framework. Pair Interaction Energy (PIE) values between probes and amino acid side chains and between probes and fluoride were computed using the Fragment Molecular Orbital (FMO) method. These PIE calculations assessed the energy differences between interacting and isolated molecular systems, thus quantitatively evaluating stability. Probes exhibiting higher PIE values indicated unfavorable, destabilizing interactions. Grid points with lower PIE values were systematically connected to adjacent points, forming networks. Smaller networks, characterized by higher cumulative PIE values, identified unstable regions within the protein, termed hotspots. Top hotspots identified through this process are residues A279, F13 and V162. Within these hotspots, residues located within a 3 to 5 Å radius were collected, and corresponding probes were recorded from the N- to C-terminal direction. These N- to C-terminal sequences became primers used to identify suitable substitutions for specific hotspots. Initial query patterns comprising three probes derived from closely placed residues of the protein within the hotspot region were searched against a predefined internal database populated with probe sequences derived from well-characterized housekeeping proteins and other structural proteins. Additional probes from the query region were incrementally added in N- and C-terminal directions to refine the search, with the closest contacts added incrementally. Database matches were filtered iteratively, retaining only those probe sequences showing exact incremental matches. From the retained database matches, the best matches were ranked based on the highest alignment score and most favorable cumulative energy. The highest-ranking probe sequence from the database was used to identify the actual amino acids corresponding to the matched database sequence. The primary rationale behind this method is that similar probes exhibit significantly matched energy profiles, thus the initial three-probe alignment serves as a direct substitution reference.

The database was built using a specific collection of housekeeping proteins and proteins with defined structural characteristics. The distinctive aspect of this method involves placing the protein within a grid and systematically positioning probes at each grid point to calculate interaction energies. These energies were used to delineate regions of varying sizes, with larger regions representing greater stability. The region with the lowest energy was selected as the core residue, initiating a detailed search for van der Waals contacts within a 3–5 Å radius. Probe sequences for the initial three residues were recorded in all possible combinations, and corresponding residues were stored. Additional residues within this defined region were incrementally assessed to verify the extendibility of the initial trimeric probe match against the comprehensive probe database.[87–89].

### S13 Site-directed mutagenesis (A279R, W50F)

For introduction of the mutation A279R, the primers 5’-gttgcgggccaggctcag-3’ and 5’cgcgccagcctggcctatccgtac-3’ were used for amplification of the template (pET28, carrying a N-terminal 6-His wild-type FlA1 or FlA1+C44T). Accordingly, for mutation W50F, the primers 5’-acggggtcatgctgtgggtcacatc-3’ and 5’-accccgttcgacgtcgaggagggcgcc-3’ were used.

After amplification, the amplicons were treated with KLD reaction mix (NEB Biolabs) according to the manufacturer's specification and used for transformation. Subsequently, the methodology described in Chapter S1 was used to express and purify the fluorinase variants.

### S14 Mass photometry

The analysis of the fluorinase oligomeric state was carried out by mass photometry using Refeyn TwoMP instrument (Refeyn Ltd., UK). Stock solutions (2 μM) of the fluorinase variants were prepared by dilution to a condition used in the kinetic experiments (30 mM HEPES buffer pH 7.5). Dilution series of protein concentrations between 31.25 and 1000 nM were prepared for each variant in (i) the buffer alone, (ii) buffer + 75 mM KF, and (iii) buffer + 75 mM KF + 800 μM SAM. For each measurement, 10 μL of the appropriate condition was pipetted to the cassette well to focus the laser signal and to obtain background. Next. 10 μL of sample was added and count acquisition was carried out for 1 min. The resulting ratiometric contrast was converted to the molecular weight using a calibration curve of bovine serum albumin (BSA) and Immunoglobulin G (IgG) standards. The histograms of the binding event counts were fitted to the sum of five Gaussian distributions described by **Equation 1** using the NumPy python package (**Figure S14**).

$$y = A/(\sigma\sqrt{2\pi})\ \exp(-1/2\ ((x-\mu)/\sigma)^2\ )$$

**Equation 1**

Where A is the amplitude, σ is the standard deviation (the width of the curve), and μ is the mean (the center of the peak). The means were fixed to $M_w$ values corresponding to the 1-, 3-, 6-, 9-, and 12-mers, whilst the upper and lower boundaries for sigma values were set to 45 and 5, respectively, to narrow the fitting parameter space. Areas under the peak were computed by numerical integration using the trapezoidal rule implemented via the np.trapz function of NumPy, and normalized to the sum of all areas to obtain a relative fraction of species.

### S15 Data analysis and statistics (oligomerization dynamics)

The equilibrium titration data recorded using mass photometry were fit globally using KinTek Explorer (KinTek, USA)[61], a dynamic kinetic simulation program that allowed multiple data sets to be fit simultaneously to a single model. Data fitting used numerical integration of rate equations from an input model (**Scheme 1**) searching a set of parameters (**Table S3**) applying the Bulirsch–Stoer algorithm with an adaptive step size that produces a minimum $\chi^2$ value calculated by using nonlinear regression based on the Levenberg-Marquardt method. Residuals were normalized by sigma value for each data point. Since it is unlikely that the trimer is formed via third-order collision, sequential reaction monomer (E) -> dimer (2E) -> trimer (3E) has been applied for monomer to trimer transition (**Scheme 1**, where E is a monomer, 2E dimer, 3E trimer, 6E hexamer, HO higher oligomer and AGG aggregated fluorinase, $k_i$ and $k_{-i}$ are forward and reverse rates for the step i, respectively). The numerical simulation of the titration data was performed under the assumption of rapid equilibrium, which was achieved by setting the forward rates ki locked at a sufficiently high value of 1 000 $s^{-1}\ \mu M^{-1}$ when integrating over time 1 s. This made it possible to directly calculate the equilibrium association constant for the i-th step, $K_i$, from the fitted values of the reverse rates k-i using the relation $K_i = 1\,000/k_{-i}$. Since no transient increase in the dimer concentration was observed during the mass photometry measurement, and the decrease in monomer concentration was directly proportional to the increase in the amount of trimer, it is not possible to accurately determine separate parameters of the second step. The overall equilibrium association constant for monomer to trimer conversion $K_{a,3E} = 1\,000/k_{-1}$ can only be obtained by fitting the parameter $k_{-1}$ with the locked value $k_{-2} = 0$ s$^{-1}$. Beyond the formation of the prevalent trimer ($K_{a,3E} = 1\,000/k_{-1}$) and hexamer ($K_{a,6E} = 1\,000/k_{-3}$), higher association complexes and aggregates are generated by the progressive addition of trimers ($k_{agg}$ and $k_{-agg}$, **Scheme 1** in gray). The steps leading to further polymerization/aggregation are characterized by the equilibrium association constant $K_{a,agg}$, expressed as $K_{a,agg} = 1\,000/k_{-agg}$, where $k_{-agg}$ represents the rate constant for dissociation of high oligomers and aggregates (**Scheme 1**).

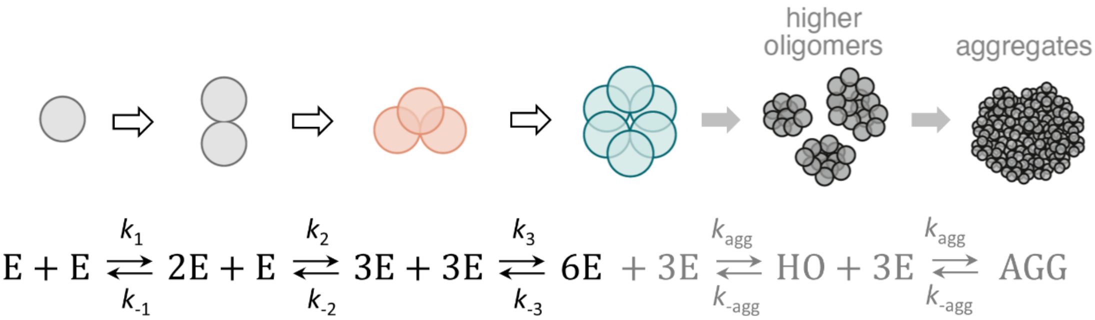

**Scheme 1**

The standard error (s.e.) of the fitted parameters was calculated from the covariance matrix during nonlinear regression. The output observable signal (**Figure S15**) was defined as the fraction of a particular oligomeric state, e.g., the monomer fraction f(E) is described in **Equation 2**, the dimer fraction f(2E) in **Equation 3**, and so on after the aggregated fraction f(AGG) is described in **Equation 6**. In addition to s.e. values, a more rigorous analysis of the variation of the kinetic parameters was accomplished by confidence contour analysis using FitSpace Explorer (KinTek Corporation, USA)[64]. In these analyses (**Figure S16**), the lower and upper limits for each parameter were derived (**Table S3**) from the confidence contour obtained from setting $\chi^2$ threshold at 0.98.

f(E) = E/(E+2*2E+3*3E+6*6E+9*HO+12*AGG)
**Equation 2**
f(3E) = (3*3E)/(E+2*2E+3*3E+6*6E+9*HO+12*AGG)
**Equation 3**
f(6E) = (6*6E)/( E+2*2E+3*3E+6*6E+9*HO+12*AGG)
**Equation 4**
f(HO) = (9*HO)/( E+2*2E+3*3E+6*6E+9*HO+12*AGG)
**Equation 5**
f(AGG) = (12*AGG)/( E+2*2E+3*3E+6*6E+9*HO+12*AGG)
**Equation 6**

### S16 Steady-state kinetics of FlA1 variants

Activity assays were carried out at 37 °C in 600 µL reactions containing 50 mM HEPES (pH 7.8), 75 mM KF, 1 µM FlA1 variant, and 1.56-800 µM SAM. Before adding SAM to initiate the reaction, the reaction mix was pre-heated for 5 mins at 37 °C. 90 μL samples were removed at 2, 5, 10, 15, 20 and 30 mins, and immediately incubated at 95 °C for 5 mins to terminate the reaction by enzyme inactivation and SAM degradation. After incubation on ice, the denatured enzyme was removed by centrifugation, and the supernatant was used for HPLC analysis of 5'-FDA concentration.

### S17 Data analysis and statistics (steady-state kinetics of wild-type and engineered FlA1 variants)

The analytical fitting was performed using OriginPro 2023, version 10.0.0.154 (OriginLab Corporation, USA). The global numerical integration analysis was performed using KinTek Explorer (KinTek, USA)[61], a dynamic kinetic simulation program that allowed multiple data sets to be fit simultaneously to a single model. The simultaneous fitting of the mass photometry and reaction kinetic data used numerical integration of rate equations from an input model searching a set of parameters applying the Bulirsch–Stoer algorithm with an adaptive step size that produces a minimum $\chi^2$ value calculated by using nonlinear regression based on the Levenberg-Marquardt method. Residuals were normalized by sigma value for each data point. The standard error (s.e.) of the fitted parameters was calculated from the covariance matrix during nonlinear regression. In addition to s.e. values, a more rigorous analysis of the variation of the kinetic parameters was accomplished by confidence contour analysis using FitSpace Explorer (KinTek Corporation, USA)[64]. In these analyses, the lower and upper limits for each parameter were derived from the confidence contour obtained from setting $\chi^2$ threshold at 0.98.

### S18 Hybrid QM/MM Simulations combined with Metadynamics

The enzyme-substrate complexes were initially prepared by embedding them in a cubic box with a 12 Å solute-solvent separation margin in all dimensions, using the QwikMD[90] program integrated into VMD. To maintain electroneutrality, $Na^+$ and $F^-$ ions were added to achieve a salt concentration of 0.2 M. The CHARMM36 force field was employed to parameterize the protein topology using the psfgen and autopsf modules, while solvation was handled using the TIP3P water model. Short-range non-bonded interactions were treated with a 12.0 Å cutoff, while long-range electrostatics were computed using the Particle-Mesh Ewald (PME) method. The r-RESPA multiple time step scheme was applied, with short-range interactions updated every step and long-range interactions every two steps, using a 2-fs integration time step.

System equilibration began with a 1000-step energy minimization using the conjugate gradients method, followed by heating to 300 K via the Langevin thermostat (collision coefficient of 1 $ps^{-1}$). The pressure was maintained at 1 atm using a barostat.

For QM/MM simulations, the QM region was defined to include Asp16, Thr80, Ser158, SAM and $F^-$. The charge of each QM region was kept between +1 and −1 to ensure effective semi-empirical QM calculations. Subsequent steps included 1000 steps of minimization and 10,000 steps of simulated annealing for equilibration. The equilibrated QM/MM structure then served as the initial input for QM/MM metadynamics simulations, an enhanced sampling technique that applies a history-dependent bias potential along selected collective variables (CVs), which correspond to specific reaction coordinates such as bond distances or dihedral angles[91,92].

The simulations were performed at 300 K and 1 bar under periodic boundary conditions, with a 0.5-fs time step ensuring stability over 1000 ps of simulation time. Using the colvars module in NAMD 2.13[93]. Gaussians with a height of 0.2 kcal/mol were deposited along the CV at each step, with a width of 1°, to construct the metadynamics bias potential. The production phase of the hybrid QM/MM simulations was conducted using the PM7 semi-empirical QM method in conjunction with the CHARMM36 force field, ensuring that the total charge remained between +1 and -1[94].

<u>S19 Metadynamics simulation of multimeric complexes</u>

Well-tempered metadynamics simulations is a technique where multiple collective variables are chosen, of which for one or both the potential will be added periodically by adding the repulsive potential of Gaussian shape at the chosen collective variable location. The fluorinase hexamer stabilization in the wild-type and the A279R were examined using well-tempered metadynamics simulations. The simulations were conducted on the hexamer complex as well as the trimer complex to understand the dynamic behaviour of both.

The GROMACS 2019.4 simulation engine patched with PLUMED 2.9.0 was used to conduct the simulations. The simulations were performed using AMBER99sb force fields with the water model TIP3P to mimic the natural environment with a box volume of 1000 nm3 to solvate the wild-type and A279R E-S complex.

<u>S20 Experimental design for simulation of Timer complexes</u>

In the trimer simulation box, for wild-type and A279R, the system was neutralized using a 0.15 mM concentration of NaF. In the wild-type simulation box, 22,975 water molecules, 98 Na+ ions, and 83 $F^-$ ions were added to the system along with the enzyme-substrate complex, which makes a total number of 82,519 atoms in the simulation setup. In the case of the A279R simulation box, 22,798 water molecules, 94 $Na^+$ ions, and 82 $F^-$ ions were added to the system along with the enzyme-substrate complex, which makes a total number of 82,025 atoms.

<u>S21 Experimental design for simulation of Hexamer complexes</u>

In the hexamer simulation box, for wild-type and A279R, the system was neutralized using a 0.15 mM concentration of NaF ions. In the wild-type simulation box, 36,881 water molecules, 169 Na+ ions, and 138 $F^-$ ions were added to the system along with the enzyme-substrate complex, which makes a total number of 1,37,775 atoms in the simulation setup. In the case of the A279R simulation box, 38,540 water molecules, 166 Na+ ions, and 142 $F^-$ ions were added to the system along with the enzyme-substrate complex, which makes a total number of 1,42,838 atoms.

For trimer and hexamer conditions, energy minimisation of the system was performed using the steepest descent algorithm with the convergence energy cut-off of 1000 kJ/mol within 1000 steps. Equilibration was done using NVT and NPT (N=constant number, V=constant volume, T=constant temperature, P=constant pressure). The NVT and NPT equilibration steps were performed for 30 ns at 300 k. The short-range cut-off values of van der Waals were set to 1.0, and columbic interactions were calculated using the PME (Particle Mesh Ewald) algorithm. LINCS algorithm was used for all bond length constraints and 2fs was set for each time step of simulations.

### S22 Metadynamics simulations

The well-tempered metadynamics simulations were conducted for both trimer and hexamer complexes with a precisely defined CVs (CV1 vs CV2) which are explained below. The CV1 was constructed with the COM of distance between the interface residues which are present in N1, N2 and N3 vs the COM of distance between the interface residues of N1′, N2′ and N3′ (Supplemental **Fig. S25A**). The residues which are present in the interface of the trimers are K237, D241, VL243, L244, P245, F246, E247, N262, I263, R277, P285, Y286 and N287. The CV2 was constructed by using the amino acid residues present in the interface of chain N1′ and chain N3′. The distance between the Centre of Mass (COM) of chain N1′ residues vs the distance between COM of chain N3′ residues defined the CV2 (Supplemental **Fig. S25B**). The residues such as H211, P212, F213, P252, T253, F254, A255, I260, G261, N278, A279, A280, S281, Y284 from chain N1′ and residues such as G18, T48, P49, W50, D51, V52, E53, P78, A79, G81, T82, T83, P154 and T155 from chain N3′ was selected for constructing CV2 (Supplemental **Fig. S25C**). The well-tempered metadynamics was conducted for the above-explained CV1 vs CV2, and the FES Graphs were plotted as a function of Gaussian free energy obtained for the COM distance of CV1 vs CV2.

The well-tempered metadynamics simulation for the trimer was conducted with a bias factor of 6.0 and temperature of 300k with the minimum Gaussian bin height of 1kJ/mol, Sigma of 0.35 nm, collective variable stored for every 20 steps and the Gaussian bias potential is deposited to the system for every 500 steps for both collective variables. The simulation was conducted for 100ns, and the coordinates were saved for every 2ps of the simulations.

In the case of hexamer simulations, the wild-type and mutant were simulated with a bias factor of 8.0 and temperature of 300k with the minimum Gaussian bin height of 1kJ/mol, Sigma of 0.35 nm, collective variable stored for every 20 steps and the Gaussian bias potential is deposited to the system for every 500 steps for the collective variables, coordinates were saved for every 2ps of the simulations.

### S23 Expression of mtnN

The 5'-methylthioadenosine/S-adenosylhomocysteine nucleosidase (mtnN/pfs) was amplified from the genome of *E. coli* DH10α with the primer pair 5'-AACATAUGAAAATCGGCATCATTGG-3'and 5'-AATTCGUCAATGGTGGTGGTGATGATGGCCATGTGCAAGTTTCTGC-3'. In parallel, the plasmids p23T7(ref) was amplified with the primer pair 5'-ACGAATUCGAGCTCGGTACCCGGGG-3' and 5'-ATATGTUTTTCCTCCTCCTAGGTCCTGGGGT. The fragments were cloned through USER cloning [95]. Transformational and protein purification was analogous to the methodology with the fluorinase.

### S24 Validation of *in silico* predicted kinetics with high substrate concentrations

Fluorinase activity assays were carried out 8-well PCR stripes containing each 100 µL reactions with 30 mM HEPES (pH = 7.8), 0.01, 0.05, 0.1, 0.5 or 1 M NaF, 0.5 µM fluorinase (wild type or W50F+A279R) and 10, 25, 50, 100, 250 or 500 µM SAM (New England Biolabs). If indicated, 0.5 µM of MtnN was supplemented. Then, the PCR stripes were sealed and incubated for 2 h at 37°C followed by 5 min at 95°C, and finally cooled to 10°C in a thermocycler (Bio-Rad). The stripes were then centrifuged for 30 min at 2,500×g, and 90-µL supernatant aliquots were transferred to 96 -well plate (conical), which were sealed with a silicone lid for HPLC analysis.

5'-FDA and SAM were quantified following absorbance at 230 nm using a Zorbax C18 column (3.5µm×4.6×100mm, Agilent) connected to a HPLC system (Dionex Ultimate 3000, ThermoFisher) with the following gradient (at an 1 mL/min flow rate): 5-12% solvent B in 1.5 min, 12% solvent B for 1 min, 12–30% solvent B in 2 min and 30-70% B in 1.5 min [solvent A: 0.05%(v/v) acetic acid in water; solvent B: acetonitrile]. A six-point calibration curve (R2 >0.99) was prepared with 0.5-50 µM 5'-FDA, which was chemically synthesized as previously described[30]. Similarly, SAM and adenine were quantified in the reactions and corrected for impurities and non-enzymatic degradation during the reaction.

### S25 Nucleotide sequences

*fla_MA37*

6xHisTag, TEV site, **ORF**, **Fla**, **Mutations:**

**W50F:** TGG271TTC

**C44T:** TGC190ACC

**A279R:** GCC898CGC

**ATGGGCAGCAGCCATCATCATCATCATCACAGCAGCGGCGAGAATCTTTATTTTCAGGGC**
**CATGCCGCCAACGGCAGCCAGCGCCCGATCATCGCGTTCATGTCCGACCTGGGCACCACCG**
**ACGACAGCGTGGCGCAGTGCAAGGGCCTGATGCACAGCATCTGCCCCGGTGTGACCGTGG**
**TGGATGTGTGCCACAGCATGACCCCGTGGGACGTCGAGGAGGGCGCCCGTTACATCGTGG**
**ACCTGCCGCGTTTCTTCCCGGAGGGCACCGTCTTCGCCACCACCACCTATCCCGCCACCGG**
**CACCACCACCCGCAGCGTGGCCGTGCGTATCCGCCAGGCCGCCAAAGGCGGCGCCCGTGGC**
**CAGTGGGCCGGCAGCGGCGACGGCTTCGAACGTGCCGACGGCAGCTACATCTACATCGCC**
**CCGAACAACGGCCTGCTGACCACCGTGCTGGAGGAACACGGCTATATCGAGGCCTACGAG**
**GTGACCAGCACCAAGGTGATCCCGGCGAACCCGGAGCCGACCTTCTACAGCCGCGAAATG**
**GTGGCCATCCCGTCCGCCCACCTGGCCGCCGGCTTCCCGCTGGCCGAGGTGGGCCGTCGTC**
**TGGATGACAGCGAGATCGTCCGTTTCCACCGCCCCGCCGTCGAGATCTCCGGCGAAGCCCT**
**GAGCGGCGTGGTGACCGCCATCGACCACCCGTTCGGCAACATCTGGACCAACATCCACCGT**
**ACCGACCTGGAAAAGGCCGGCATCGGCCAGGGCAAACACCTGAAGATCATCCTGGACGAC**
**GTGCTGCCGTTCGAAGCCCCGCTGACCCCCACCTTCGCCGACGCCGGCGCCATCGGCAACA**
**TCGCCTTCTACCTGAACAGCCGCGGCTATCTGAGCCTGGCCCGCAACGCCGCCAGCCTGGC**
**CTATCCGTACAACCTGAAAGCCGGCCTGAAGGTGCGCGTGGAGGCCCGTTGA**

*mtnN_Ecol*

*T7Pr*, *rbs*, **ORF** 6xHisTag,

*TACGACTCACTATAGTAAGGAGA*AAATAATTTTGTTTAACTTTAACACCCCAGGACCTAGGAGGA
GGAAAAACAT**ATGAAAATCGGCATCATTGGTGCAATGGAAGAAGAAGTTACGCTGCTGCGT**
**GACAAAATCGAAAACCGTCAAACTATCAGTCTCGGCGGTTGCGAAATCTATACCGGCCAACT**
**GAATGGAACCGAGGTTGCGCTTCTGAAATCGGGCATCGGTAAAGTCGCTGCGGCGCTGGGT**
**GCCACTTTGCTGTTGGAACACTGCAAGCCAGATGTGATTATTAACACCGGTTCTGCCGGTGG**
**CCTGGCACCAACGTTGAAAGTGGGCGATATCGTTGTCTCGGACGAAGCACGTTATCACGACG**
**CGGATGTCACGGCATTTGGTTATGAATACGGTCAGTTACCAGGCTGTCCGGCAGGCTTTAA**
**AGCTGACGATAAACTGATCGCTGCCGCTGAGGCCTGCATTGCCGAACTGAATCTTAACGCTG**
**TACGTGGCCTGATTGTTAGCGGCGACGCTTTCATCAACGGTTCTGTTGGTCTGGCGAAAATC**
**CGCCACAACTTCCCACAGGCCATTGCTGTAGAGATGGAAGCGACGGCAATCGCCCATGTCTG**
**CCACAATTTCAACGTCCCGTTTGTTGTCGTACGCGCCATCTCCGACGTGGCCGATCAACAGT**
**CTCATCTTAGCTTCGATGAGTTCCTGGCTGTTGCCGCTAAACAGTCCAGCCTGATGGTTGAG**
**TCACTGGTGCAGAAACTTGCACATGGCCATCATCACCACCACCATTGA**

### Supplementary text

S26 *In silico* mutant design

#### *Diffusion of fluoride into the apoenzyme, the E·SAM complex and the E·5'-FDA complex*

SMD and umbrella sampling simulation studies were used to study the diffusion of F- into the apoenzyme, the E·SAM complex, and the E·5'-FDA complex. F- diffusion into the apoenzyme is relatively straightforward and energetically favorable. Conversely, in the presence of SAM, F- doesn't reach the active site, but instead forms a stable hydrogen bond with non-catalytic amino acids Thr155 and Ser269 (**Figure S9**). This correlates well with the ordered sequential mechanism of fluorinase, which requires fluoride to bind as the first substrate. In the presence of 5'-FDA in the active site, the absence of the methionine moiety increased the likelihood of F- approaching the active site. As seen from the kinetic analysis, such binding of fluoride to the enzyme-product complex disables 5'-FDA release from the active site, causing the inhibitory effect of fluoride.

#### *Inhibition of the enzyme-product complex by fluoride occurs through a bridging interaction*

A metadynamics study of the E·5'-FDA and F-·E·5'-FDA complexes was carried out (**Figure S10**) to gain insight into the mechanism of fluoride inhibition. In the case of E·5'-FDA, the sampled conformations at 5-7Å from Asp16 likely correspond to the release of the product. The increased sampling in CV2 indicates that the product release is facilitated by the interactions of the ribose sugar rather than the adenine. If a new fluoride diffuses into the product-bound active site (forming F-·E·5'-FDA), it has a probability of forming a hydrogen bonding bridge between 5'-FDA and the active site environment. Specifically, the ion has a chance to bind to the catalytic Ser and Thr residues and forms a hydrogen bonding network with the ribose-sugar moiety of the product at the 2' and 3' hydroxyl groups (**Figure S11**). This bridging interaction hyperstabilizes the complex of fluorinase and 5'-FDA, and is likely responsible for inhibiting the release of the product.

S27 Dynamics of fluorinase oligomerization

The equilibrium association constants for the formation of a trimer ($K_{a,3E}$), a hexamer ($K_{a,6E}$), and higher oligomers ($K_{a,agg}$) are reported in **Table S3** and **Figure S17**. The equilibrium parameters provide insight into the binding affinities and stability of intermediates at different stages of the oligomerization process, as well as how mutations influence these processes. The comparison between buffer conditions and the presence of substrates shows that substrate presence significantly affects the oligomerization behaviour of all variants. The dynamic nature of the fluorinase oligomerization makes it difficult to directly compare how individual mutations impact the final population of each oligomeric state by inspecting these constants alone. To better illustrate the overall impact of mutations on the distribution of individual oligomeric forms, we used the numerical model to simulate the equilibrium distribution of oligomeric forms (**Figure S18**) under the conditions used for the steady-state kinetic experiments.

For the free enzyme, only the A279R mutation increased the efficiency of trimer formation compared to the wild type, while all other mutations resulted in a decrease in the $K_{a,3E}$ value (**Figure S17a**). However, in the presence of substrates, the conversion from monomer to trimer ($K_{a,3E}$) was closely similar for all tested variants, including wild type and mutants. Interestingly, when fluoride alone was added, the rate of the initial oligomerization steps increased for both wt and mutants, suggesting that charged ligands might lower the barrier for the initial dimer formation. The exception was A279R, which showed reduced efficiency in the first oligomerization step ($K_{a,3E}$) when both fluoride and SAM were present.

The A279R variant uniquely showed a significant increase in the efficiency of hexamer formation. Especially in the presence of both substrates, there was an increase of $K_{a,6E}$ by two orders of magnitude (**Figure S17b**, please note that the y-axis is on a logarithmic scale). Under these conditions, the hexamer became the dominant form for the A279R variant (**Figure S18c**). This effect was fully transferred to the double mutant W50F+A279R. In both variants, the A279R substitution significantly increases the efficiency of trimer assembly into hexamers ($K_{a,6E}$); however, at the same time, the mutation increases the ability to form higher-order oligomers, nonamers and dodecamers, which may contribute to protein aggregation ($K_{a,agg}$). W50F enhances the formation of trimers, in the presence of both substrates, whereas in the presence of fluoride only, it forms mostly hexamers.

Compared to the wild type, all introduced mutations generally reduce the formation of higher oligomers, decreasing the enzyme's tendency to aggregate (**Figure S18**). This effect is especially pronounced in the presence of substrates (**Figure S18c**). The most potent inhibition of high oligomer formation, particularly in the presence of substrates, was observed for the anti-aggregation mutation C44T. This mutation reduced the $K_{a,agg}$ value (**Figure S17c**), leading to the most significant decrease in the formation of unpreferred higher oligomers (**Figure S18c**). When combined with the W50F+A279R mutations, the resulting triple mutant (C44T+W50F+A279R) showed both anti-aggregation properties and enhanced hexamer formation, with the most substantial effect occurring in the presence of reaction substrates.

If an enzyme's active site involves multiple monomeric subunits, the substrate may simultaneously interact with more than one subunit, potentially influencing the enzyme's oligomeric state. Engineered mutations designed to enhance substrate binding could promote the formation of specific oligomeric assemblies, such as highly active hexamers, as seen in the case of fluorinase. This concept introduces an interesting paradigm in protein engineering, wherein controlling oligomerization through the optimization of enzyme-substrate interactions can lead to enhanced catalytic activity.

### S28 Steady-state kinetics of wild-type and engineered variants of FlA1

*Conventional data fitting*

The concentration dependence of initial velocities, derived from a linear fit of raw kinetic data (**Figure S19**), was subsequently analyzed using the conventional fitting to the hyperbolic steady-state Michaelis-Menten model (**Figure S20A**). Although this hyperbolic model initially appeared to capture the general trend of the data, a closer examination of the residuals revealed significant systematic deviations, indicating that the simple Michaelis-Menten model did not fully account for the observed kinetics. In particular, kinetic data for substrate concentrations up to 200 μM SAM displayed a clear substrate inhibition pattern (**Figure S20B**). At higher substrate concentrations, however, the data trended more closely with a hyperbolic model, suggesting a shift in kinetic behavior. These observations imply a mixed kinetic profile, likely due to the presence of two enzyme species with distinct kinetic characteristics. When the data were analysed using a complex model accounting for two distinct enzymatic species (**Figure S20C**), the fit closely matched the experimental data, as evidenced by the residuals, which were small and randomly distributed.

Kinetic analysis revealed that two distinct forms of the enzyme exhibit similar turnover numbers ($k_{cat}$) but differ significantly in their Michaelis constant ($K_M$). This finding is consistent with previous studies on fluorinase engineering, which demonstrated that the isolated trimeric and hexameric forms of the enzyme differ primarily in their $K_M$ values, 206 ± 34 and 3.7 ± 0.7 μM, respectively, while exhibiting nearly identical $k_{cat}$ values, 0.20 ± 0.01 and 0.22 ± 0.01 $min^{-1}$[39]. To differentiate the individual contributions in mixed kinetics, where both trimeric and hexameric forms may coexist, it is crucial to determine their relative proportions under the specific experimental conditions. To address this, we studied the oligomeric behavior of fluorinase using mass photometry (**Figure S14**). First, we analyzed how the enzyme behaves in its free state and combined this information with catalytic data for a global analysis. Subsequently, we applied the concentration estimates for the trimer and hexamer, determined by mass photometry under specified reaction conditions, in the analytical fitting to get refined estimates of kinetic parameters (**Figure S20D**). However, the free enzyme oligomeric data alone could not explain the observed kinetics, and we couldn't find a unified model that connected the structural and functional observations. To resolve this, we examined how the substrate affects the enzyme's oligomeric state. This deeper analysis revealed the complex behavior of fluorinase and finally allowed us to develop a unified kinetic model that explains both its structural changes and functional activity, including its mutants employing global numerical integration.

*Global numerical analysis*

Although analytical fitting provides valuable insights into enzymatic mechanisms, it has notable limitations. These challenges include difficulties in accurately estimating parameters due to approximations and the accumulation of errors across multiple fitting steps. Additionally, this approach does not enable the clear identification of enzyme states, such as distinguishing between trimeric and hexameric forms. The assignments used in the analysis described in **Figure S20D** were based on previously reported data[39]. Moreover, analytical fitting is insufficient for resolving the global kinetic relationships between functional and structural observations.

To overcome these limitations, we implemented global numerical fitting to analyze reaction kinetics alongside structural data obtained from mass photometry. This approach preserves all raw data, allowing for the extraction of additional information from kinetic curves, which is crucial for accurate parameter estimation, particularly under non-linear conditions. Moreover, it considers the active enzyme population participating in the reaction. Notably, for the fluorinase reaction under the tested conditions, the active enzyme concentration was

found to differ significantly from the total enzyme added (**Figure S19**). This is primarily due to the fluoride concentration being below the enzyme-fluoride dissociation constant ($[F^-] < K_{d,F}$) and also due to the incorrect-order binding of SAM to the enzyme. Collectively, the results from analytical fitting, mass photometry analysis, and observations of the active site titration effect were integral in defining the minimal kinetic model (**Figure S21A**). This model enabled rigorous numerical analysis through the simultaneous fitting of both kinetic and structural data directly in its raw form (**Figure S21B-D**). The confidence contour analysis (**Figure S21E**) indicates that most of the obtained parameter estimates (**Figure S21F**) are well constrained by the data involved in the global analysis. However, for the trimer reaction, it was not possible to determine a unique solution for separate $k_{cat}$ and $K_M$ values. This is already evident from the high values of the standard errors. Systematic confidence contour analysis further indicated that these two parameters strongly correlate, and only their ratio, $k_{cat}/K_M$, could be accurately determined.
Subsequently, we performed the same global analysis for the mutant variants (**Figure S22**). Statistical analysis revealed correlations between $k_{cat}$ and $K_M$ for both trimeric and hexameric forms in some mutants. To ensure accurate comparisons, the final numerical analysis applied $k_{cat}/K_M$ ratio for both trimer and hexamer catalytic efficiency, providing a precise and consistent comparison across all tested variants (**Table S4**). Similarly, the natural correlation between $K_{d,F}$, and $K_{i,SAM}$, which define the active enzyme fraction, was addressed by fitting their ratio, $K_{i,SAM}/K_{d,F}$, to enhance model accuracy and interpretability. Recognizing standard error analysis can yield overly optimistic uncertainty estimates for fitted parameters, we have additionally provided rigorous confidence contour analysis (**Figure S23**). As a result, the lower and upper confidence limits were determined and used for interpreting the kinetic results, avoiding reliance on potentially misleadingly low standard errors.

### S29 *In silico* characterization of mutants

#### *Metadynamics simulations of Hexamer Complexes*

The hexamer is a complex simulation box with a higher number of atoms in the simulation box; an average of 1,40,000 atoms were present in the simulations. The CVs were defined to study the events which could occur during the formation of a hexamer and stabilize the trimer. The CV for the well-tempered metadynamics was selected based on the contacting regions of the trimer-trimer interface in the hexamer complex. The contacting residues, which are present in this region, stabilise the hexamer complex, multiple salt-bridges between residues such as K237-D247, R277-D241 and many other VdW interactions were identified. Based on the interactions and the interface where trimers become hexamer this region was selected for defining the CVs.

To dissect the assembly process, individual chains of each trimer were tracked: chains A, B, and C (trimer-1) were mapped as N1, N2, N3; and chains D, E, F (trimer-2) as N1′, N2′, N3′. The planar angle between these trimers (N1–N3 vs N1′–N3′) was calculated to monitor conformational transitions (**Supplementary Fig. S26a**).

Well-tempered metadynamics simulations revealed that the A279R mutant adopts a more stable hexamer conformation compared to the wild type. Conformational transitions during the simulation indicated distinct hexamer assembly pathways, with A279R showing faster and more stable trimer association. Root Mean Square Deviation (RMSD) analysis confirmed reduced fluctuations in A279R trimers relative to wild-type (**Supplementary Fig. S26b**). A279R showed a distinct angle–distance trajectory (**Supplementary Fig. S26c**), reflecting a unique, non-overlapping pathway toward hexamer formation. A plot of distance vs. time (**Supplementary Fig. S26d**) further illustrated these differences: in the wild-type, N1 and N1′ associate first (event E1), while in A279R, simultaneous association of N1/N2 with N1′/N2′ was observed. The graphs indicate the distance of association of N2 with N2′ in A279 and wild-type.

The wild-type simulation shows 3 different events that occurred during the hexamer association, where Event 1 (E1) happens with the first chain N1 coming in contact with the N1′. Residues such as K237, D242, V243, L244, P245, E247, R299 from N1 and K237, L244, P245, F246, E247, R277, and Y286 from N1′ were found within the contacting range. E1 was mapped on the FES, at the CV1=49.988Å and CV2=10.061 Å with the free energy value of -38.973kcal/mol. Once N1 and N1′ were in contact, Event 2 (E2) was observed with a longer time interval as the N2 and N2′ chains had to obtain an energetically favourable path to make contact. E2 is a crucial conformation attained during the simulation which facilitates the hexamer stability. The association of N2 and N2′ was observed with the similar residues observed in the N1 and N1′ interactions as the interface residues are similar across all chains, V243 from N2 and Y286 from N2′ residues were observed to make the initial contact, followed by D241 and K237 of N2 forming a crucial salt bridge with R277, and E247 of N2′, respectively for the stabilization of the N2 and N2′. E2 state was slightly better energetically, compared to E1, as it is closer to forming the hexamer complex with CV1 = 30.417 Å and CV2 = 10.010 Å and free energy value of -48.7846 kcal/mol. The final event observed in the wild-type was Event 3 (E3), where N3 forms a stable interaction with the N3′ and forms a stable hexamer complex. The initial interacting residues were similar to N1-N1′, and the final complex was found with 2 different salt bridges across each of the interacting regions of the hexamer. The final hexamer complex was observed in the E3 event where CV1 = 10.358 Å and Cv2 = 9.061 Å with the free energy of -93.928 kcal/mol. Across the hexamer formation events, the overall common contacting residues from all the inter-trimer chains were found to be K237, D241, D242, V243, L244, P245, F246, E247, N262, I263, R277, Y286, E297, and R299. A279R shows different or unique ways of forming hexamer associations when compared to the wild-type. The events observed in the A279R hexamer simulations occurred much faster with an energetically favourable path due to the conformational transitions which were observed in A279R. The enhanced stability in A279R is attributed to stronger inter-chain interactions. Specifically, R279 of N1 chain forms salt bridges with D51 and E53 of N2 chain, stabilizing the trimer. This lowers the energy barrier for hexamer assembly. Additional stabilizing contacts between R279 and residues W50, I260, and N278, along with inter-chain salt bridges to N3, further reinforce the dimer and trimer interfaces. Together, these interactions support a more stable and efficient assembly pathway for A279R, facilitating rapid and energetically favourable hexamer formation, and provide plausible explanation of additive effects observed with A279R+W50F mutant (**Supplementary Figure S27**).

*Metadynamics simulation of Trimer complexes*

The CVs are important factors of the metadynamics, which delineate the stability and function of amino acids in the given range of CVs. The trimer well-tempered simulation was conducted by selecting 2 different CVs. The contacting regions between two chains, which are crucial to maintain the interchain stability were mapped, crucial residues which stabilizes the dimeric interface were selected for constructing the CV. The CV1 was constructed by using the amino acid residues present in the interface of chain N1′ and chain N2′. The distance between the Centre of Mass (COM) of chain N1′ residues vs the distance between COM of chain N2′ residues defined the CV1. The residues such as H211, P212, F213, P252, T253, F254, A255, I260, G261, N278, A279, A280, S281, Y284 from chain N1′ and residues such as G18, T48, P49, W50, D51, V52, E53, P78, A79, G81, T82, T83, P154, and T155 from chain N2′ was selected for constructing CV1. CV2 was defined more precisely in the active site, The COM of distance between the catalytic residues such as S158 and T80 of N2′, the F- which is bound to these catalytic residues Vs the distance of COM of SAM was considered for defining the CV2. The well-tempered metadynamics was conducted for the above-explained CV1 vs CV2, and the FES Graphs were plotted as a function of Gaussian free energy obtained for the COM distance of CV1 vs CV2.

The well-tempered metadynamics simulation was conducted on wild-type and A279R trimer and hexamer complexes to extrapolate the stability of the complexes across the simulations. The simulations conducted on the trimer complex showed higher stability in the A279R mutant. The FES graphs of the wild-type and A279R show different minima obtained by the wild-type which is energetically not favourable as compared to the A279R (**Supplementary Figure S28a,b**). Well-tempered metadynamics simulations were performed to analyze the stability of wild-type and A279R complexes, where the wild-type and A279R mutant Gaussian energy was taken at the lowest minima points observed on the FES plots. The wild-type had a Gaussian energy of -53.63 kcal/mol at CV1 of 0.9981 and CV2 of 0.6745, while the A279R mutant found with more favourable energy of -70.44 kcal/mol at when CV1 is 0.9901 and CV2 is 0.6069. These findings indicate the unique stable conformations of the trimers, of which the mutant prefers an energetically more favourable minima. The minima observed in the wild-type are much enhanced in the A279R with much lower energy indicating stable conformation. The RMSD graphs of the backbone of the protein extracted for the trimers also indicate the stable conformation of the A279R trimer complex compared to the wild-type simulation (**Supplementary Fig. S28c**).

### S30 Application of fluorinase in optimized conditions

To validate the optimal process conditions predicted by our model, the activity of wild-type fluorinase and W50F+A279R mutant was assayed at varied substrate concentrations (**Supplementary Figure S29a**,**b**). Both fluorinase variants displayed the highest activity at high substrate concentration: 0.5 mM SAM and 1000 mM NaF for the wild-type, and 1 mM SAM and 1000 mM NaF for W50F+A279R. The results align closely with the estimates obtained from the kinetic model (**Figure 3b**), which predicted maximum normalized productivity of 0.6-0.8 $min^{-1}$ at 500 mM $F^-$ and 0.1-1 mM SAM. While the wild-type fluorinase reached the predicted productivity (0.59 $min^{-1}$), the mutant showed lower productivity (0.4 $min^{-1}$) than expected. Based on the predictions from our mechanistidc model, we hypothesised that the productivity could be increased by depleting the product 5'-FDA from the reaction mixture **(Figure 3e)**. Therefore, we added *S*-adenosylhomocysteine nucleosidase (MtnN), which converts 5'-FDA into adenine and 5'-fluoro-5'-deoxyribose ($K_m$ of 15.2 ± 0.7 µM and $k_{cat}$ of 5.79 ± 0.5 $min^{-1}$ for 5'-FDA), thereby recapitulating the downstream steps of the metabolic pathway. Through addition of equal molar amounts of fluorinase (0.5 µM) and MtnN (0.5 µM), accumulation of 5'-FDA should be omitted. Indeed, the productivity of both fluorinases increased dramatically (**Supplementary Figure S29c**,**d**). The wild-type fluorinase reached a productivity of 5 $min^{-1}$, while the mutant reached 8.3 $min^{-1}$. Interestingly, in contrast to the reaction without 5'-FDA consumption, lower $F^-$ concentration were favorable for high productivity. This observation agrees with the predicted fluorinase behaviour under 5'-FDA consumption and suppression of inhibition by SAM (no E·SAM) (**Figure 3e**). Due to the unexpectedly high productivity, SAM was completely depleted in several reactions after 2 h (**Supplementary Figure S29d**). Additionally, up to 13 µM 5'-FDA accumulated in several reactions (**Supplementary Figure S30**). Against this background, we modified the reaction conditions to optimize productivity: 500 mM NaF, 0.5 mM SAM, 1 µM MtnN. To avoid depletion of the substrates, the reaction time was decreased to 45 min. Under these conditions, the wild-type fluorinase has a productivity of ~3.7 ± 0.49 $min^{-1}$, while the mutant W50F+A279R exhibited a normalized productivity of 12.5 ± 2.1 $min^{-1}$ and a yield of the halogenated product 244 µM after 45 min (**Supplementary Figure S31**). The productivities observed in our experiments with optimized conditions dramatically surpassed previously reported activity of with the wild-type fluorinases of 0.16 $min^{-1}$ (0.8 mM SAM and 0.075 M KF)[22] and 0.26 $min^{-1}$ (0.8 mM SAM and 0.2 M KF)[26], measured at low substrate concentrations that mimic natural conditions.

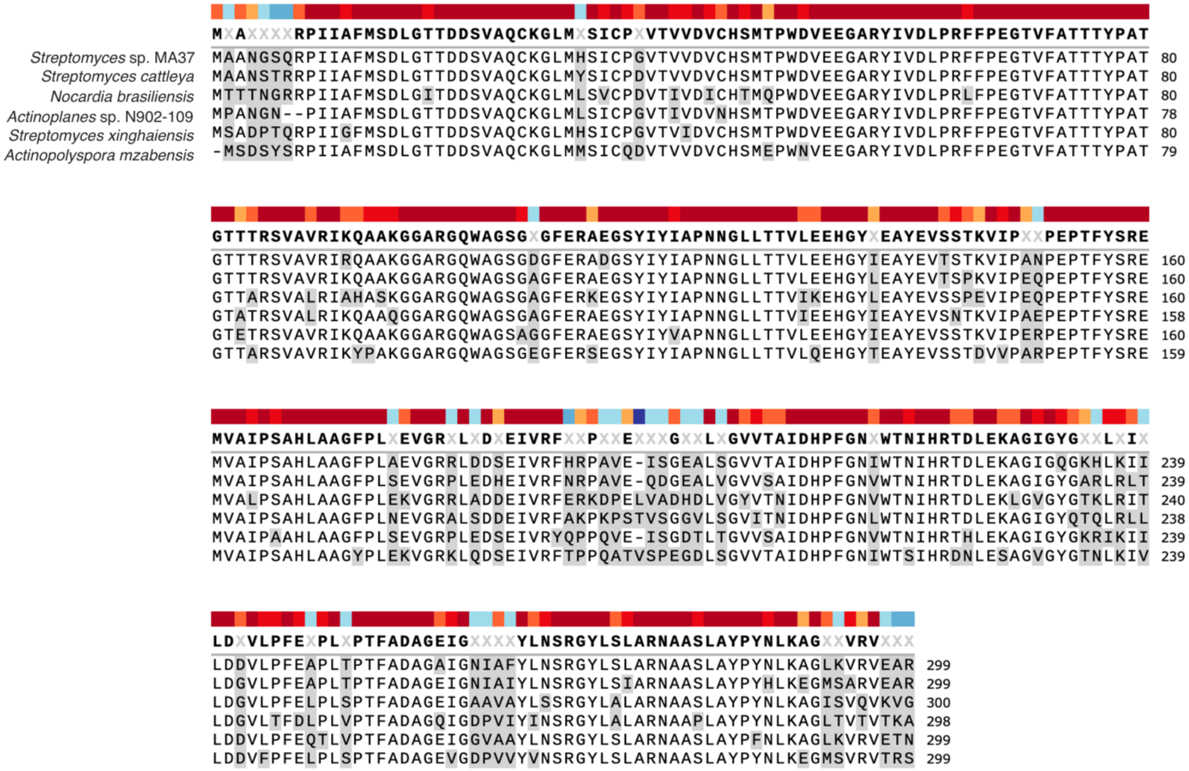

**Fig. S1.**

**Multiple sequence alignment of several bacterial fluorinases**. Sequences were imported from UniProt: W0W999 (*Strepotomyces* sp. MA37), Q70GK9 (*Streptomyces cattleya*), W8JNL4 (*Nocardia brasiliensis*), R4LHX8 (*Actinoplanes* sp. N902-109), A0A068VNW5 (*Streptomyces xinghaiensis*) and A0A1G9FQX8 (*Actinopolyspora mzabensis*) into SnapGene, and aligned using MUSCLE multiple sequence alignment tool. Consensus residues are shown with a threshold of >50% (86.3% amino acids). Residues with a lower consensus are highlighted in grey.

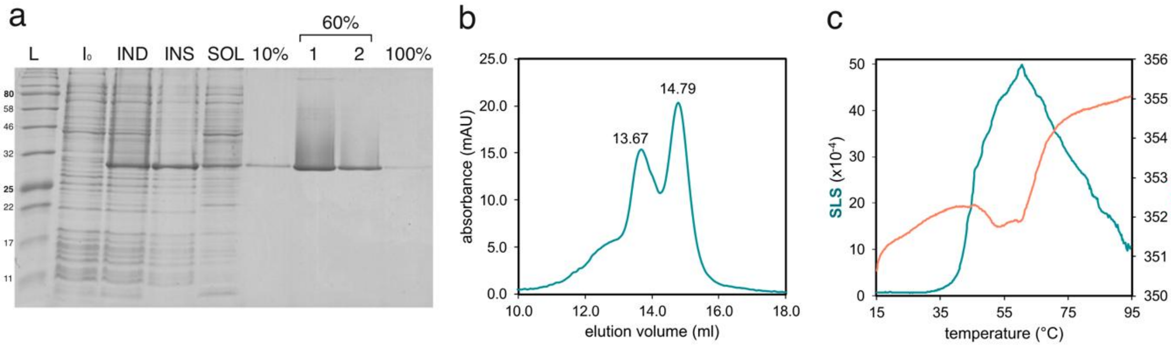

**Fig. S2.**

**Biochemical characterization of FlA1.** a) SDS-PAGE analysis of whole cell lysate before induction of FlA1 ($M_W$ = 34.4 kDa) expression ($I_0$), whole cell lysate after FlA1 induction with 0.5 mM IPTG (IND), the insoluble fraction of the lysate (INS), the soluble fraction of the lysate (SOL) and fractions from FlA1 purification (10%, 60% and 100% or purification buffer B). Only 29 ± 9 % of FlA1 was produced in the soluble fraction. Fluorinase eluted in the second fraction of the 60% gradient was used for kinetic experiments. b) Chromatogram from analytical size-exclusion chromatography of FlA1 (1 mg ml$^{-1}$, 200 $\mu$l) in 30 mM HEPES (pH 7.8), 75 mM KF. Hexamers were eluted at 13.67 ml (experimental Mw = 183.2 kDa, theoretical Mw = 206 kDa) and trimers at 14.79 ml (experimental Mw = 95.23 KDa, theoretical Mw = 103 kDa). The third peak at a lower retention volume indicates the presence of aggregates. c) Temperature dependence of FlA1 (2.1 mg ml$^{-1}$) aggregation followed by static light scattering (SLS) and barycentric mean fluorescence (BCM) in 30 mM HEPES, 75 mM KF (pH 7.8). In the case of non-aggregating proteins, the BCM value typically increases at higher temperatures because tryptophan residues become more exposed to the polar solvent during unfolding, leading to a red shift of the spectra to longer wavelengths. However, in the case of fluorinase, instead of the characteristic sigmoidal curve, a blue shift to shorter wavelengths was observed at ~45 °C. Simultaneously, the scattering signal increased, indicating the formation of bigger particles, such as aggregates.

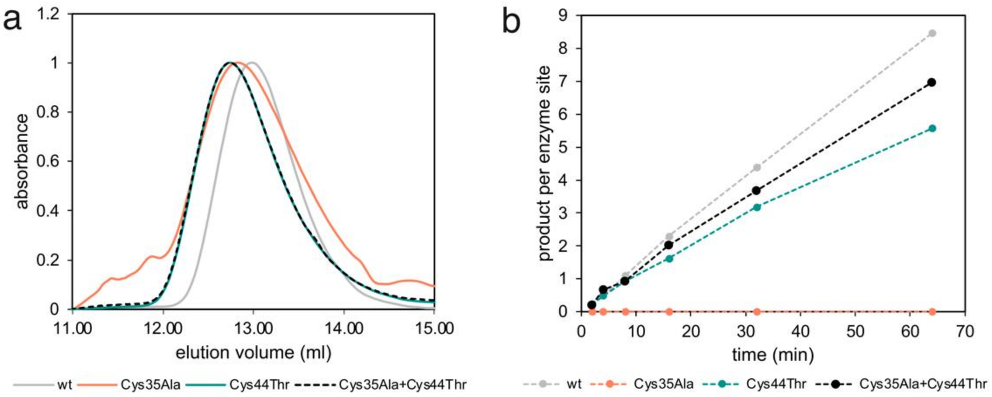

**Fig. S3.**

**Characterization of fluorinase variants with mutated cysteines. a)** size exclusion chromatography confirmation of the oligomeric state of FlA1 wild-type (wt) and its variants (1 mg ml$^{-1}$) measured in 30 mM HEPES, 400 mM KF, and 10 mM DTT (pH 7.8). The elution volumes correspond to hexameric fluorinase. **b)** activity of fluorinases variants (6 $\mu$M) measured with 800 $\mu$M SAM and 400 mM KF in 30 mM HEPES (pH 7.8), 10 mM DTT, at 25 °C. HPLC signal from 5'-FDA was used as the readout.

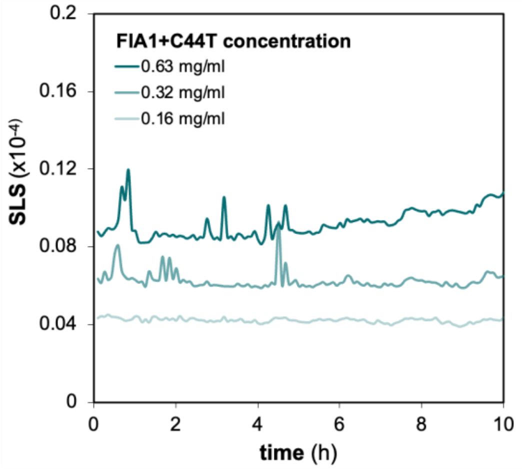

**Fig. S4.**

**Aggregation of the stabilized FlA1+C44T mutant at 30 °C.** FlA1+C44T of different concentrations was incubated at 30 °C in 30 mM HEPES, 75 mM KF (pH 7.80).

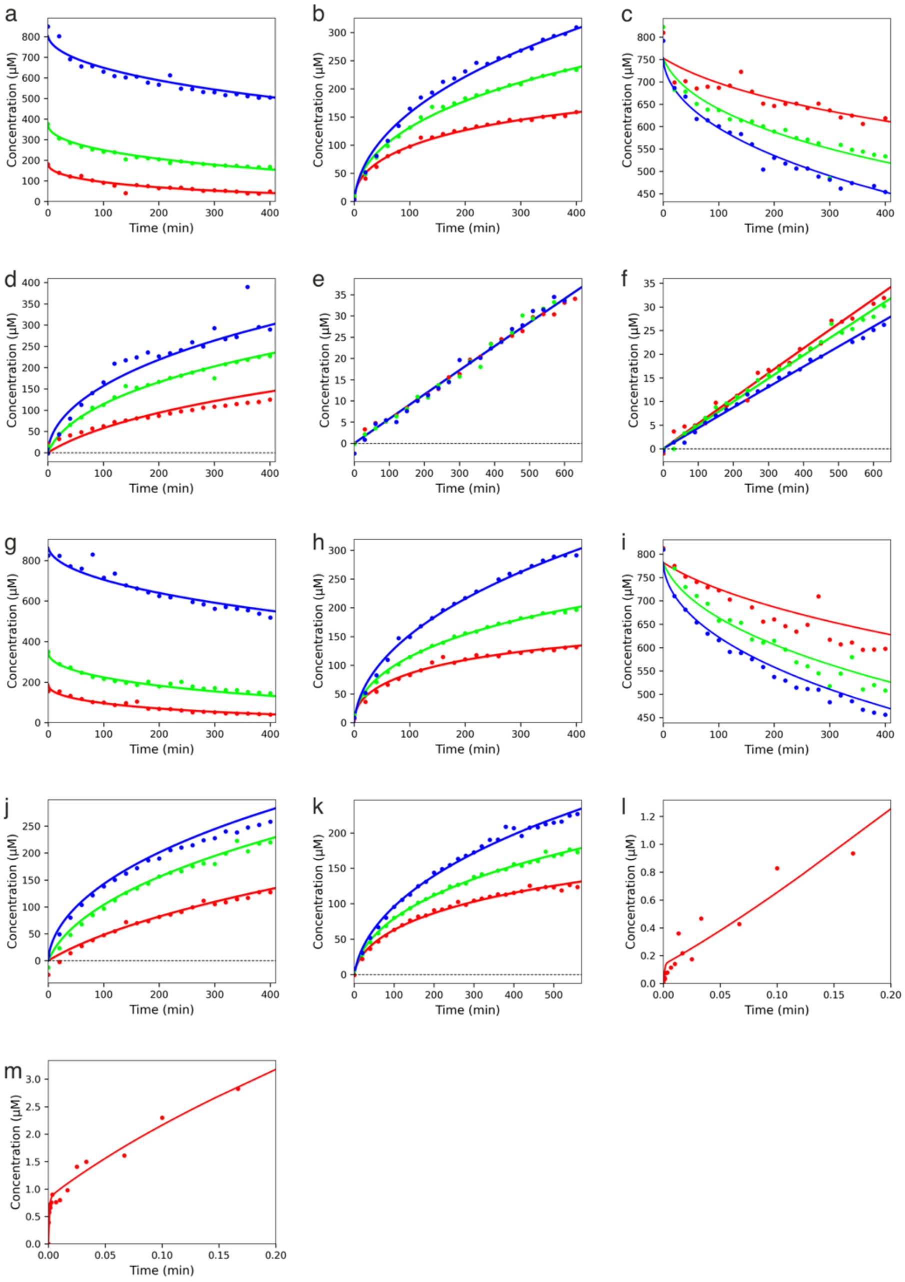

**Fig. S5.**

**Global analysis of FlA1+C44T kinetic datasets.** The lines represent the best fit and correspond to the kinetic mechanism in **Figure 3** of the main text. All experiments were conducted in 30 mM HEPES and 150 mM NaCl (pH 7.8). **a)** Kinetics of 12 $\mu$M FlA1+C44T with concentration series of SAM (800, 400, 200 $\mu$M) and 400 mM KF followed by SAM consumption. **b)** Kinetics of 12 $\mu$M FlA1+C44T with concentration series of SAM (800, 400, 200 $\mu$M) and 400 mM KF followed by 5'-FDA formation. **c)** Kinetics of 12 $\mu$M FlA1+C44T with concentration series of KF (25, 100, 800 mM) and 800 $\mu$M SAM followed by SAM consumption. **d)** Kinetics of 12 $\mu$M FlA1+C44T with concentration series of KF (25, 100, 800 mM) and 800 $\mu$M SAM followed by 5'-FDA formation. **e)** Kinetics of 25 $\mu$M FlA1+C44T in the reverse direction (defluorination) with concentration series of 5'-FDA (400, 800, 1600 $\mu$M) and 5 mM L-Met followed by SAM formation. **f)** Kinetics of 25 $\mu$M FlA1+C44T in the reverse direction (defluorination) with concentration series of L-Met (800, 1600, 3200 $\mu$M) and 1.6 mM 5'-FDA followed by SAM formation. **g)** Kinetics of 12 $\mu$M FlA1+C44T with the reverse reaction blocked by L-amino acid oxidase (2.13 mmol $l^{-1}$ $min^{-1}$), with concentration series of SAM (800, 400, 200 $\mu$M) and 400 mM KF followed by SAM consumption. **h)** Kinetics of 12 $\mu$M FlA1+C44T with the reverse reaction blocked by L-amino acid oxidase (2.13 mmol $l^{-1}$ $min^{-1}$), with concentration series of SAM (800, 400, 200 $\mu$M) and 400 mM KF followed by 5'-FDA formation. **i)** Kinetics of 12 $\mu$M FlA1+C44T with the reverse reaction blocked by L-amino acid oxidase (2.13 mmol $l^{-1}$ $min^{-1}$), with concentration series of KF (25, 100, 800 mM) and 800 $\mu$M SAM followed by SAM consumption. **j)** Kinetics of 12 $\mu$M FlA1+C44T with the reverse reaction blocked by L-amino acid oxidase (2.13 mmol $l^{-1}$ $min^{-1}$), with concentration series of KF (25, 100, 800 mM) and 800 $\mu$M SAM followed by 5-'FDA formation. **k)** Kinetics of 12 $\mu$M FlA1+C44T preincubated with SAM (800, 400, 200 $\mu$M) for 30 mins, with 400 mM KF followed by 5'-FDA formation. **l)** pre-steady-state burst of 29 $\mu$M FlA1+C44T with 400 $\mu$M SAM and 400 mM KF, followed by 5'-FDA formation. **m)** pre-steady-state burst of 29 $\mu$M FlA1+C44T preincubated with 400 mM KF, with 400 $\mu$M SAM, followed by 5'-FDA formation.

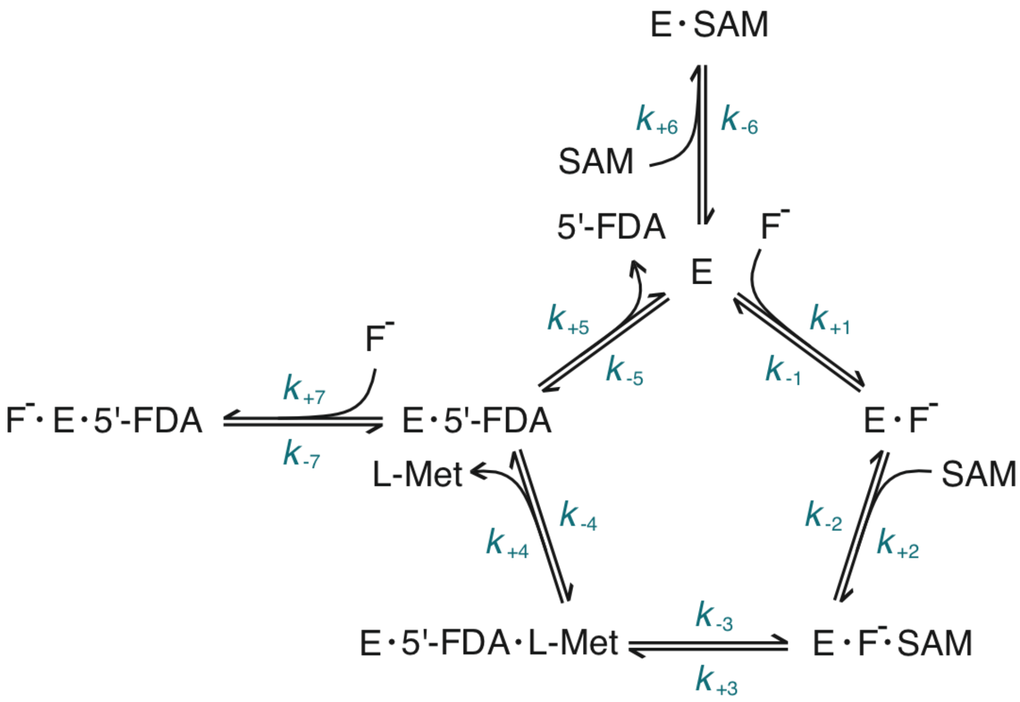

**Fig. S6.**

Input model for fitting kinetic datasets of FlA1+C44T. E – enzyme (FlA1+C44T).

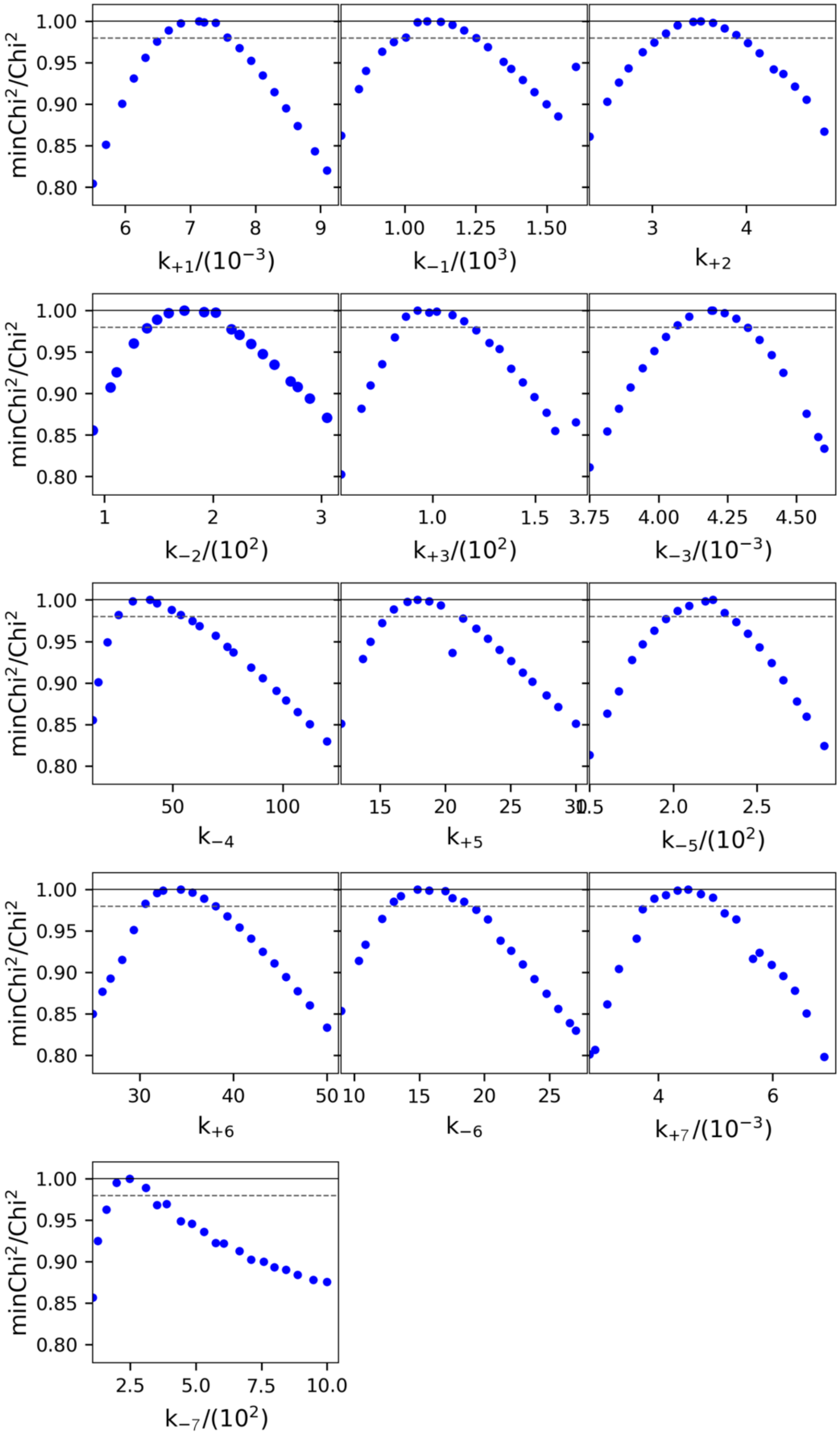

**Fig. S7.**

FitSpace 1D confidence contour analysis of the derived kinetic constants. $K_4 = 10000/k_{-4}$.

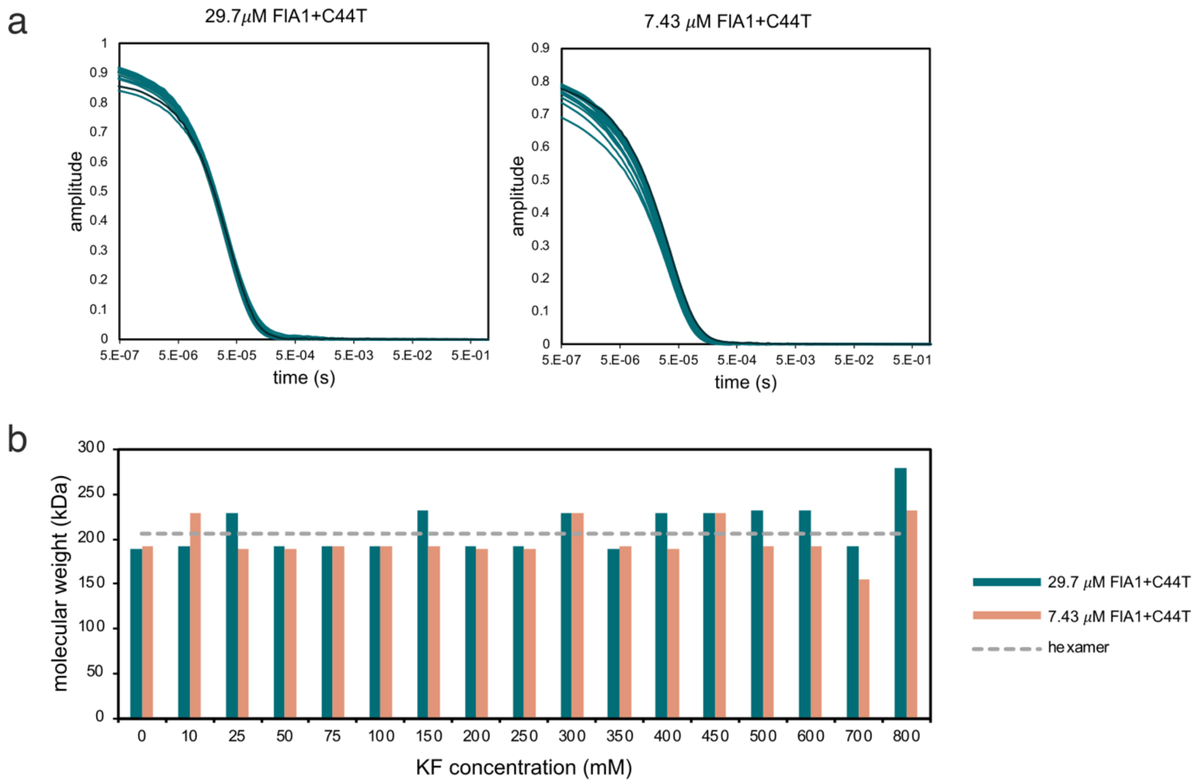

**Fig. S8.**

**Oligomeric state of FlA1+C44T during kinetic experiments measured by DLS. a)** Correlation functions. **b)** Calculated molecular weight at varying concentrations of KF. All samples of FlA1+C44T were hexameric (206 kDa). Two enzyme concentrations were analyzed in 30 mM HEPES (pH 7.8) with varying concentrations of KF. NaBr was present in all samples to keep the total salt (KF+NaBr) concentration at 850 mM.

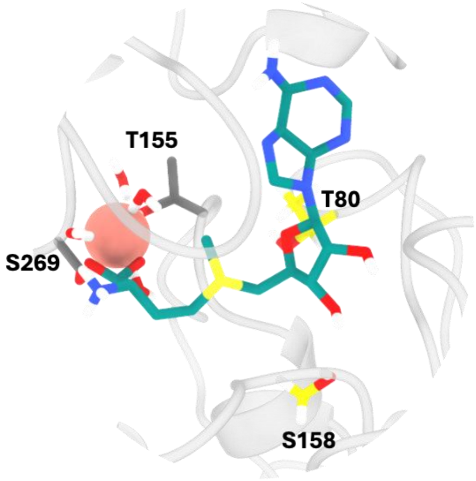

**Fig. S9.**

$F^-$ is bound to a non-catalytic Ser-Thr pair (grey sticks) instead of the catalytic pair (yellow sticks).

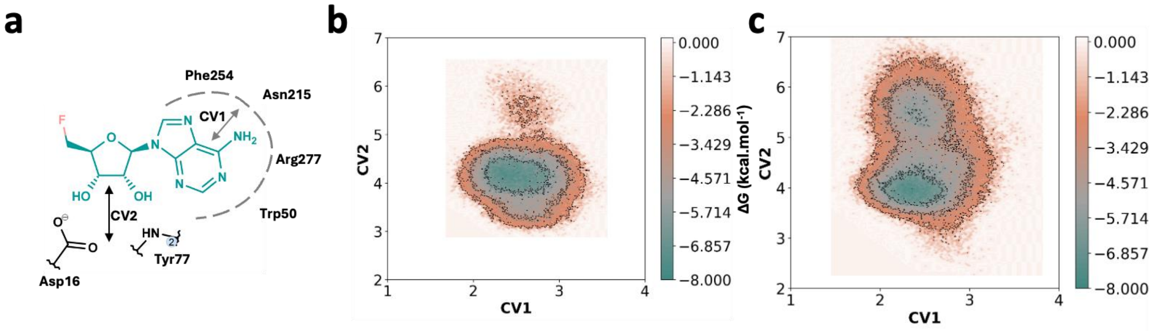

**Fig. S10.**

**Metadynamics study of FlA1 inhibition by fluoride. a)** CV1: distance between the COM of $FDA_{ade}$ and the COM of residues interacting with $FDA_{ade}$ in Å. CV2: Distance between the COM of ribose-sugar of FDA and the COM of residues Asp16 and Tyr77 in Å. Free energy landscape exploring the potential energy wells formed in the conformational sampling derived from metadynamics study of SAM in the active site of the **b)** E·5'-FDA and **c)** $F^-$·E·5'-FDA complexes.

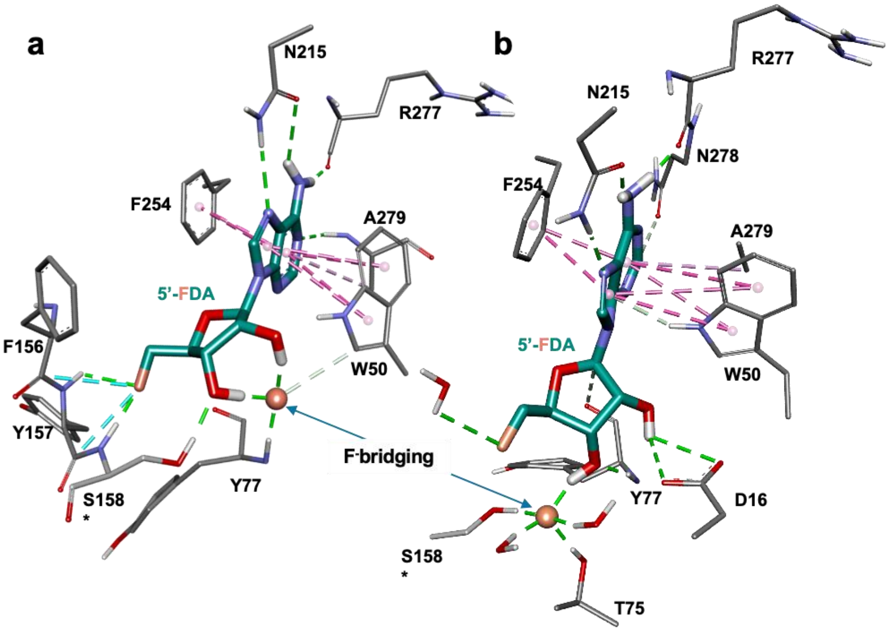

**Fig. S11.**

**Representation of a bridge interaction formed between enzyme residues, $F^-$, and 5'-FDA in the inhibited $F^-$·E·5'-FDA complex.** The blue arrow indicates the hydrogen bond bridge that is formed between the $F^-$, nearby residues in the active site, hydroxyl of the sugar moiety of the product, and in one case with two water molecules. **a)** and **b)** are two snapshots derived from two different simulations to depict the $F^-$ bridging concept. Product FDA, interacting residues, and catalytic residue Ser are represented in pink, grey, and green sticks, respectively. $F^-$ is represented as ca yan sphere. Hydrogen bonds are presented as green dashed lines. Hydrophobic interactions are depicted as pink dashed lines.

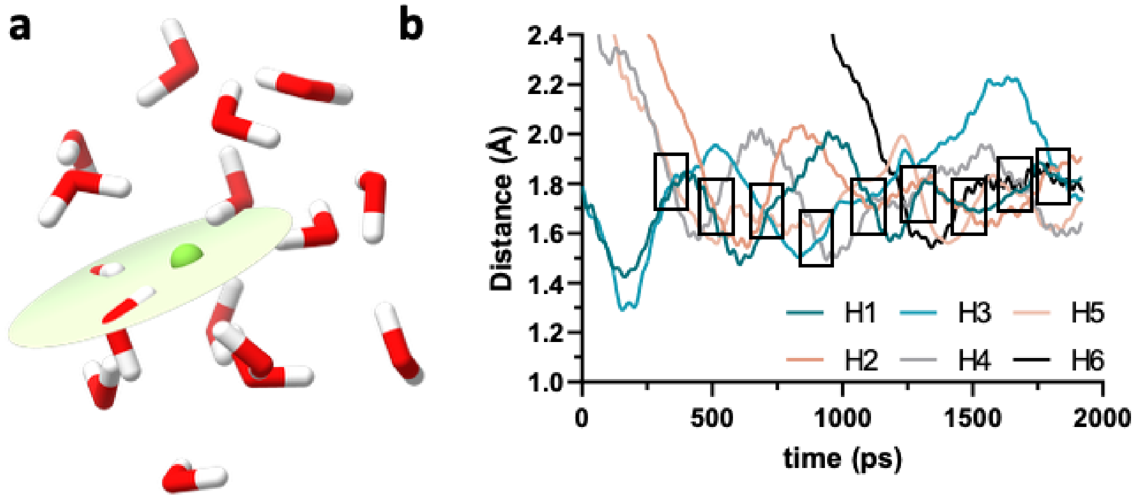

**Fig. S12.**

**Planarity of water shell formed during $F^-$ diffusion**. **a)** The $F^-$ interacts with water molecules through H-bonding and these water molecules lie in a plane. **b)** Convergence of hydrogen bonding distances of different water molecules with the $F^-$. Highlighted rectangles indicate events where a different water molecule shows interactions with $F^-$**.**

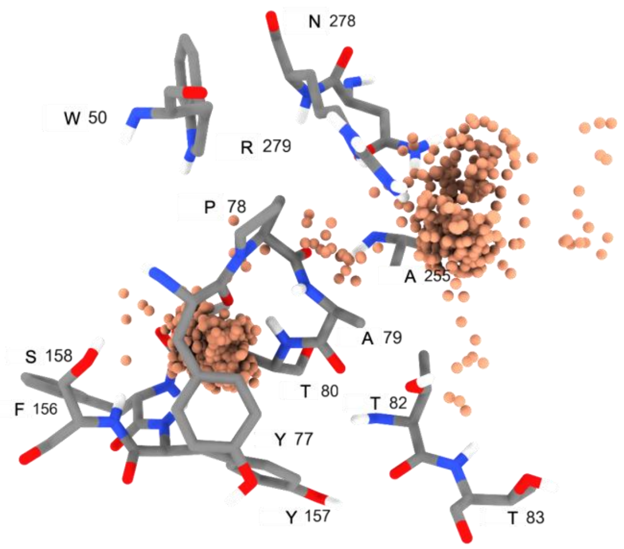

**Fig. S13.**

Residues of the prime cluster (grey sticks) interacting with $F^-$ (orange spheres).

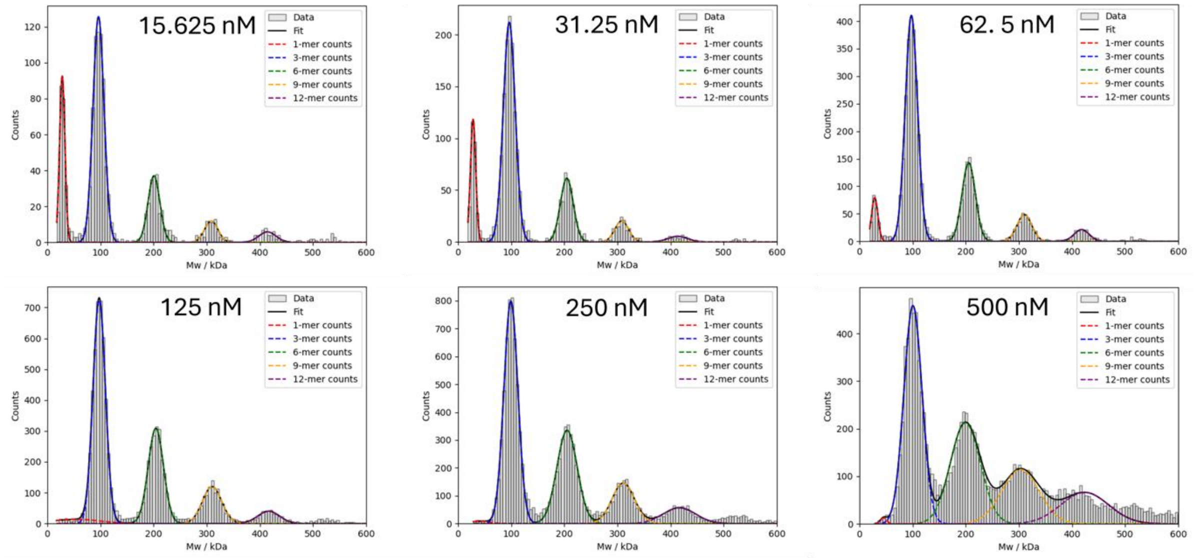

**Fig. S14.**

**Mass photometry analysis of fluorinase oligomerization.** The histograms of the binding event counts fitted to the sum of five Gaussian distributions described by **Equation 1** using the NumPy python package. The example of the data was recorded for the wild-type fluorinase at concentrations ranging from 15.625 to 500 nM in 30 mM HEPES buffer, pH 7.5, 20°C. For clarity, the species labelled as 9-mers in the figure legend represent higher oligomeric (HO) assemblies, defined operationally by their apparent mass in the mass photometry analysis, whereas the 12-mer population marks the transition to non-specific aggregate formation rather than a discrete functional oligomer.

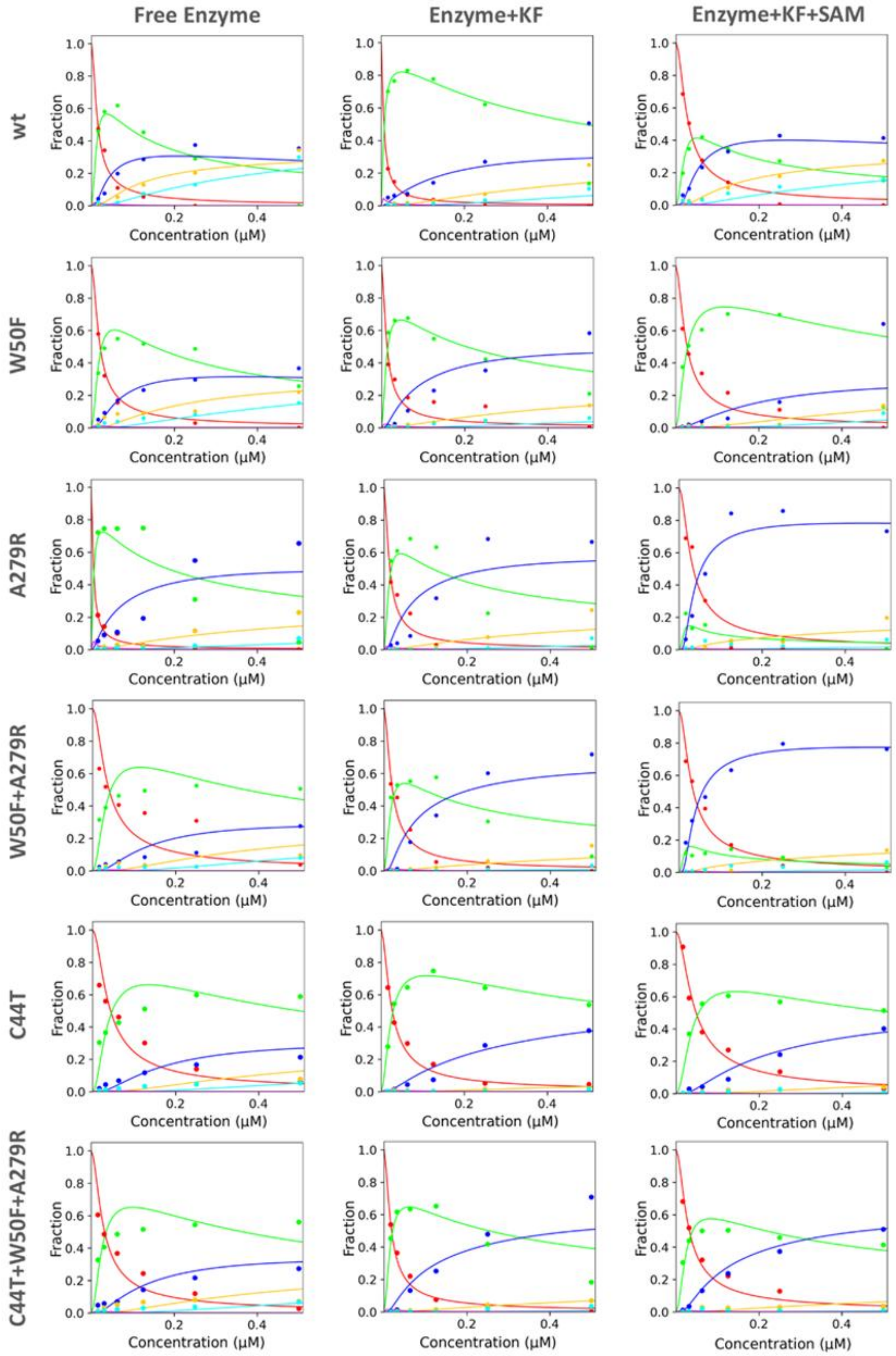

**Fig. S15.**

**Global numerical analysis of fluorinase oligomerization.** Mass photometry data (dots) for wild-type and mutants (W50F, A279R, W50F+A279R, C44T, C44T+W50F+A279R) were fitted to a model of oligomeric states (lines: monomer-red, trimer-green, hexamer-dark blue, nonamer-yellow, dodecamer- turquoise). Experiments were conducted under various conditions: buffer (30 mM HEPES, pH 7.5, 20°C), 75 mM KF (presence of fluoride substrate), and 75 mM KF + 800 μM SAM (presence of both substrates). The concentration range of the fluorinase was 0.0156-0.5 μM.

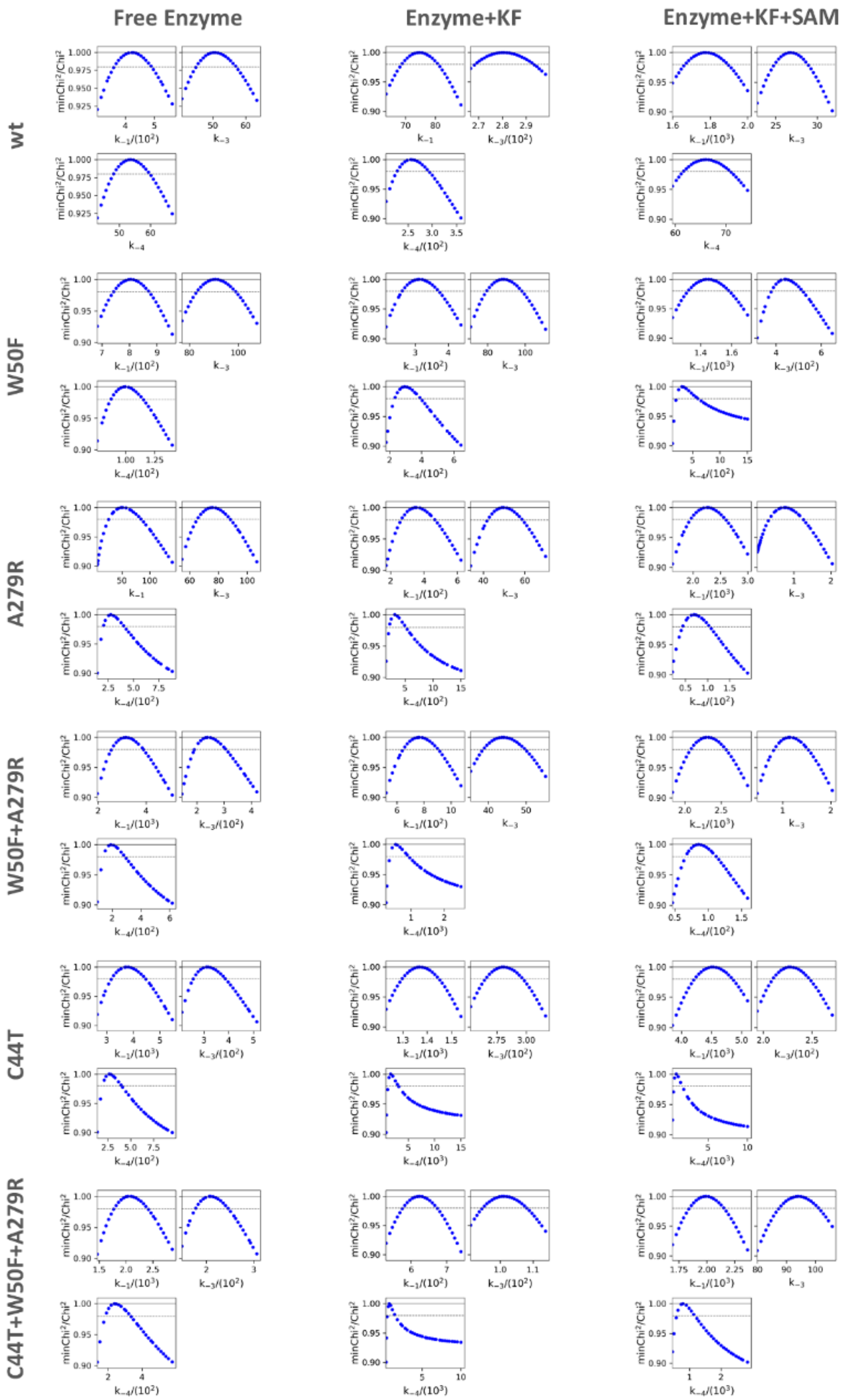

**Fig. S16.**

**Confidence contour analysis.** Individual figures represent the dependence of the error on the respective parameter while varying all other parameters to achieve the best fit. In every case, the dashed line shows the $\chi^2$ threshold (0.98) used to establish confidence intervals, as reported in **Table S3**. Experiments were conducted for the free enzyme in buffer (30 mM

HEPES, pH 7.5, 20°C), the enzyme in the presence of fluoride substrate (75 mM KF), and in the presence of both substrates (75 mM KF + 800 µM SAM).

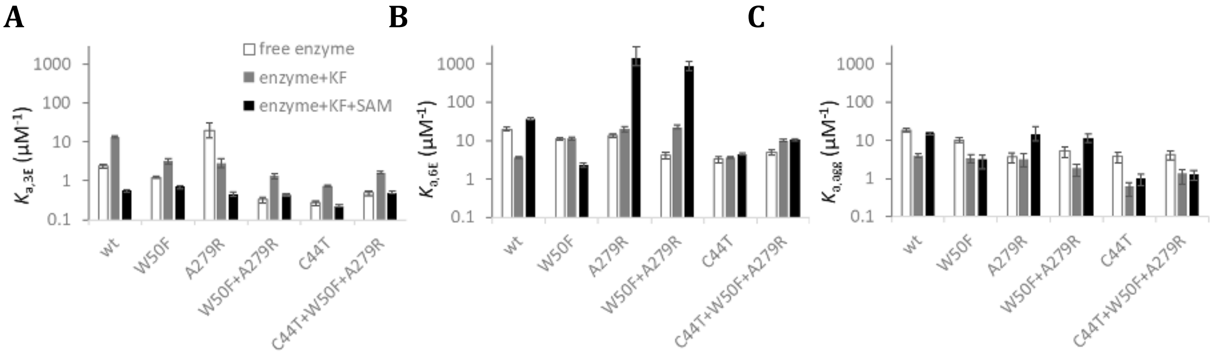

**Fig. S17.**

**Comparison of the equilibrium constants for wild-type and mutants.** The equilibrium association constants for (A) monomer to trimer transition $K_{a,3E}$, (B) formation of a hexamer $K_{a,6E}$, and (C) formation of higher oligomers $K_{a,agg}$. Error bars represent the confidence intervals (lower and upper limits) of the parameters determined by confidence contour analysis with a $\chi^2$ threshold of 0.98. Experiments were conducted for free enzyme in buffer (30 mM HEPES, pH 7.5, 20°C), the enzyme in the presence of fluoride substrate (75 mM KF), and in the presence of both substrates (75 mM KF + 800 µM SAM).

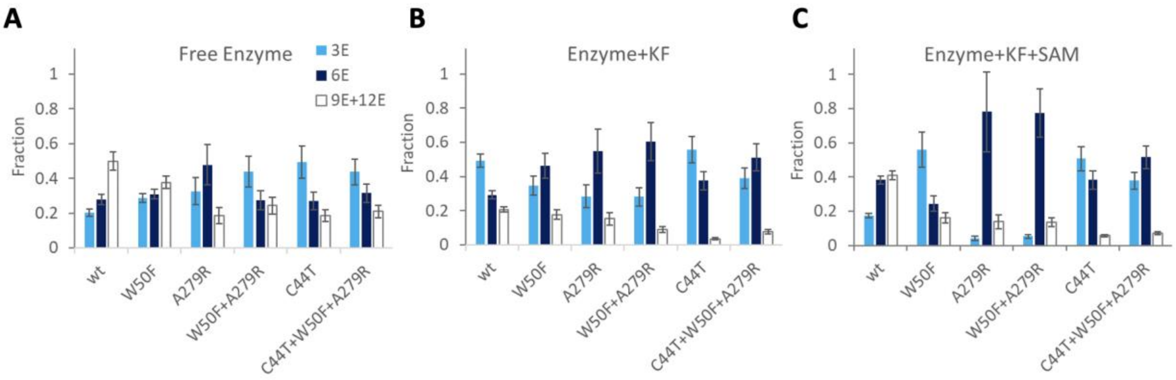

**Fig. S18.**

**Simulation of fluorinase oligomerization.** The distribution of individual oligomeric states, trimer (3E), hexamer (6E), and higher oligomers (HO+AGG), was simulated for the free enzyme in 30 mM HEPES buffer, pH 7.5, 20°C (A), the enzyme in the presence of 75 mM KF (B), and in the presence of both substrates 75 mM KF and 800 μM SAM (C) at enzyme concentration 0.5 μM used for steady-state kinetic experiments. Error bars represent the confidence intervals.

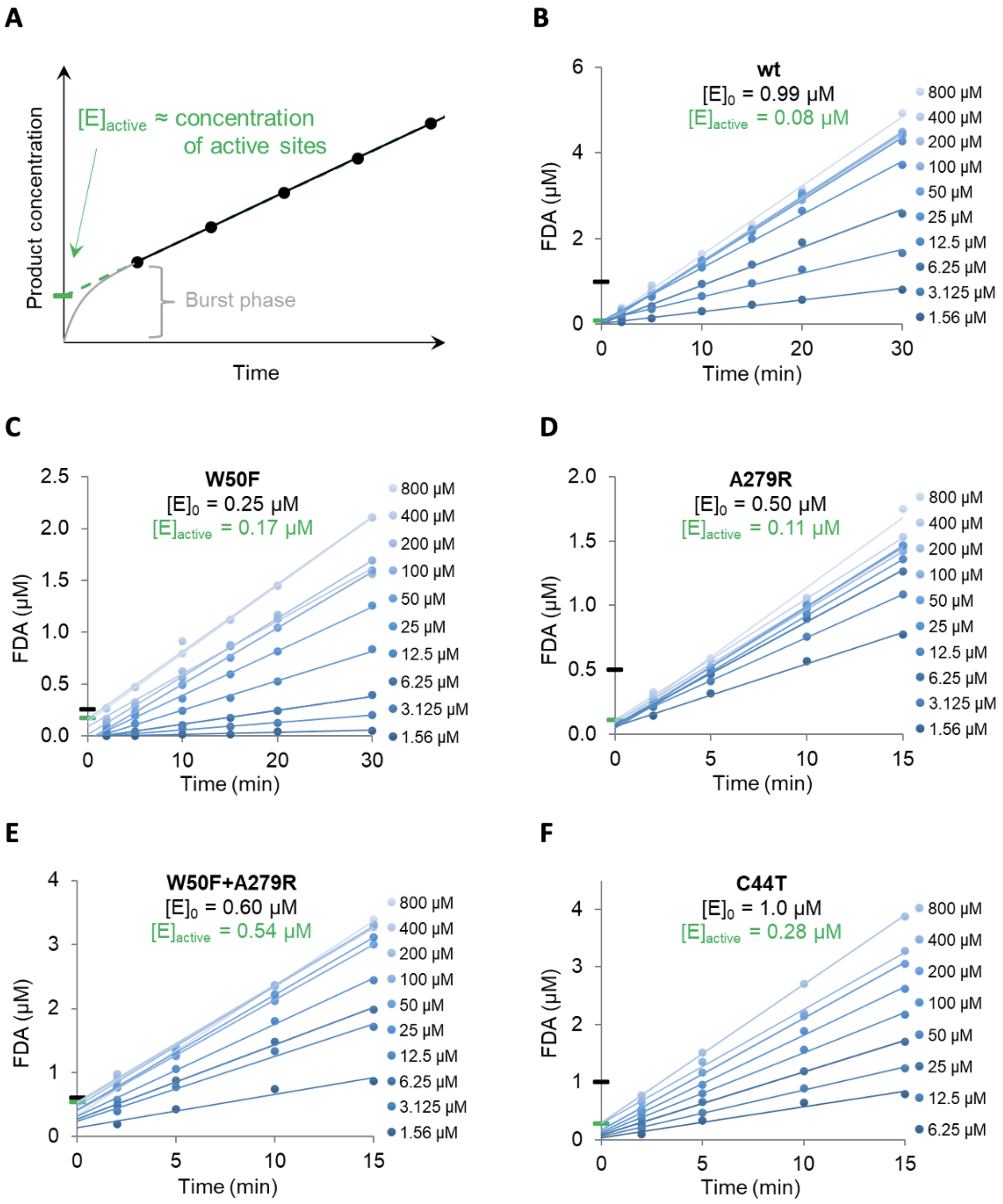

**Fig. S19.**

**Analytical fitting of the steady-state kinetic data of wild-type FlA1.** The kinetic data was collected from 1.56 to 800 $\mu$M of SAM, 75 mM KF in 50 mM HEPES, pH 7.8 at 37 °C. Solid lines represent linear fits to the raw data (average of three replicates). The velocity parameters derived from these fits have been utilized for subsequent modeling and secondary fitting (**Figure S20**). The y-intercept of the activity data (**A**) indicates the concentration of active enzyme sites participating in the reaction (green mark), even when the transient kinetic phase (the burst phase) is not observable in the steady-state kinetic data. The black marks indicate the initial FlA1 concentration added to the reaction $[E]_0$ = 0.99, 0.25, 0.50, 0.60 and 1.0 $\mu$M of wt (**B**), W50F (**C**), A279R (**D**), W50F+A279R (**E**) and C44T (**F**), respectively, the green markers denote the largest intercept obtained during the linear fit of the activity data indicating the fraction of active enzyme $[E]_{active}$ under SAM saturation.

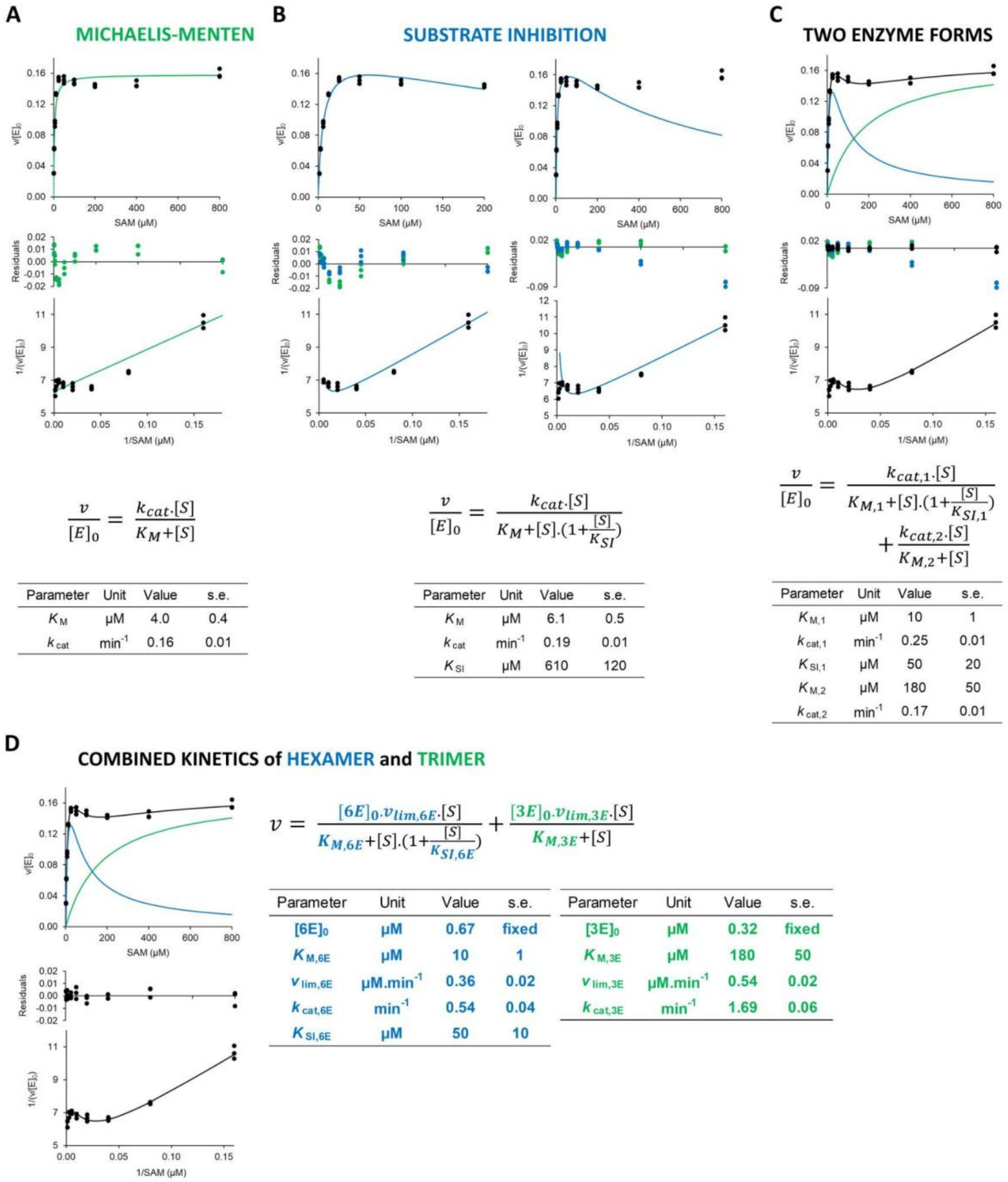

**Fig. S20.**

**Analytical fitting of the steady-state kinetic data of wild-type FlA1.** The kinetic data was collected with 1.56-800 $\mu$M SAM, 1 $\mu$M FlA1 and 75 mM KF in 50 mM HEPES, pH 7.8 at 37 °C. (**A**) The best fit is shown for hyperbolic Michaelis-Menten, (**B**) substrate inhibition at first 200 µM (left) and full kinetic data (right), and (**C**) two-enzyme form kinetics.For each model type, the fit of the reaction rate dependence on substrate concentration (top panel), the residuals (middle panel), and the corresponding double-reciprocal Lineweaver–Burk plot (bottom panel) are shown. Data points show the three replicated measurements. The equations used and the corresponding parameter estimates are shown below each fit. (**D**) Refined estimates of kinetic parameters when concentration for the trimer and hexamer obtained by mass photometry was included in the analytical fitting. Each datapoint represents the average of three replicated measurements.

A

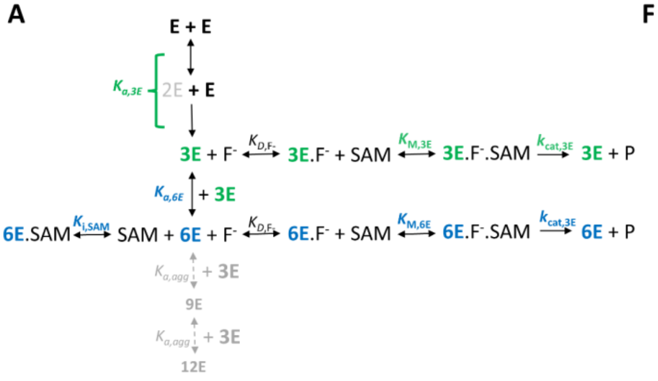

F

| Parametr | Unit | value ± s.e. | lower limit | upper limit |
|---|---|---|---|---|
| $K_{a,3E}$ | $\mu M^{-1}$ | 0.56 ± 0.03 | 0.51 | 0.62 |
| $K_{a,6E}$ | $\mu M^{-1}$ | 34.6 ± 1.7 | 30.6 | 38.8 |
| $K_{a,agg}$ | $\mu M^{-1}$ | 14.0 ± 0.8 | 12.7 | 15.2 |
| $K_{m,3E}$ [a] | μM | 128 ± 98 | n.d. | n.d. |
| $k_{cat,3E}$ [a] | $min^{-1}$ | 2.0 ± 1.5 | n.d. | n.d. |
| $k_{cat,3E}/K_{m,3E}$ | $\mu M^{-1}.min^{-1}$ | 0.016 ± 0.001 | 0.015 | 0.017 |
| $K_{m,6E}$ | μM | 9.9 ± 0.7 | 9.8 | 12.1 |
| $k_{cat,6E}$ | $min^{-1}$ | 1.5 ± 0.1 | 1.4 | 1.8 |
| $k_{cat,6E}/K_{m,6E}$ [b] | $\mu M^{-1}.min^{-1}$ | 0.15 ± 0.02 | n.d. | n.d. |
| $K_{D,F}$ | mM | 72 ± 6 | 68 | 78 |
| $K_{i,SAM}$ | μM | 1.16 ± 0.01 | 1.10 | 1.22 |

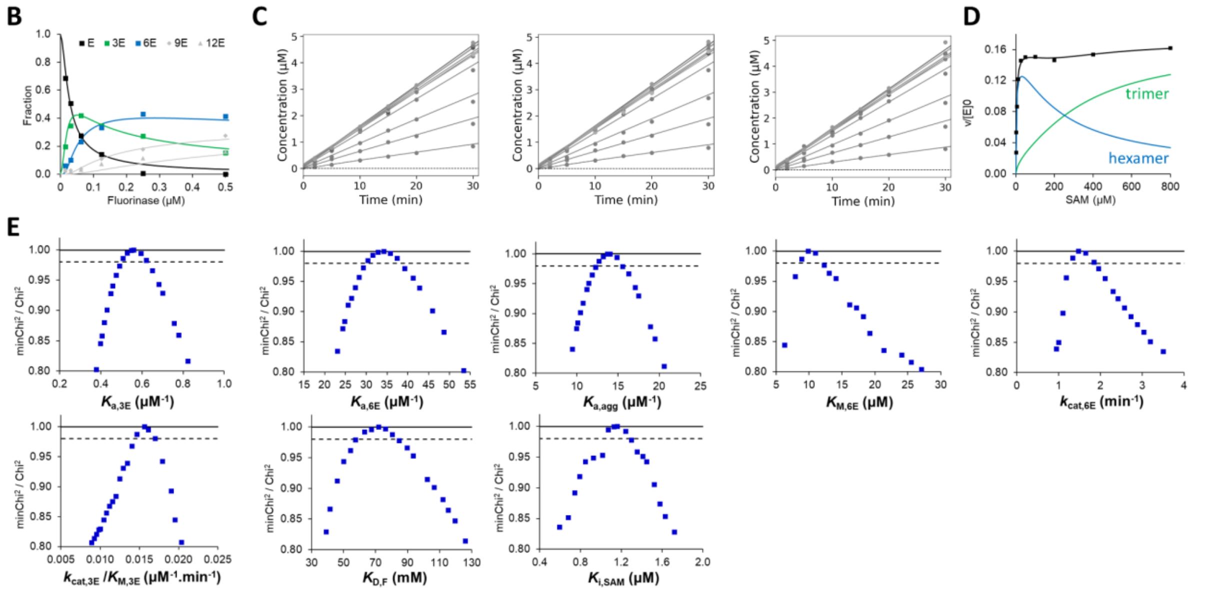

**Fig. S21.**

**Global analysis of steady-state kinetics and oligomerization data for the wild-type FlA1.** The input kinetic model (**A**) depicts enzyme species in various oligomeric states E (monomer), 3E (trimer), 6E (hexamer), HO and AGG (higher oligomers and aggregates) with association constants $K_{a,3E}$ (trimer), $K_{a,6E}$ (hexamer), and $K_{a,agg}$ (higher-order oligomers leading to aggregation). $K_M$ is the Michaelis constant, and $k_{cat}$ is the turnover number. $K_{i,SAM}$ represents the dissociation constant of the inhibitory enzyme-SAM complex. $K_{d,F}$ is the dissociation constant of the active enzyme-fluoride complex. Mass photometry data (**B**) were collected in 30 mM HEPES, pH 7.5, at 20 °C, with 75 mM KF and 800 μM SAM. Raw kinetic data (**C**) collected in triplicate across 1.56–800 μM SAM with 75 mM KF in 50 mM HEPES, pH 7.8, at 37 °C. The SAM concentration dependence of initial rates (**D**), data points show an average of triplicate measurements. Solid lines in **B**, **C** and **D** represent the global fit. Confidence contour analysis (**E**) shows parameter error dependence while varying other parameters for the best fit. The dashed line indicates the $\chi^2$ threshold (0.98) used to define confidence limits. Panel (**F**) summarizes parameter estimates, with standard errors (s.e.) calculated from the covariance matrix during nonlinear regression, upper and lower confidence limits obtained from the statistical analysis shown in **E.** [a]The $k_{cat}$ and $K_M$ for the trimer along with their respective standard errors (**F**) determined during the initial global fit. Due to the strong correlation between these individual parameter values, the secondary analysis focused on their ratio, $k_{cat}/K_M$, to enable a more robust and reliable parameter estimation. [b]The specificity constant ($k_{cat}/K_M$) for the hexamer was calculated from fitted $K_M$ and kcat values, with standard errors propagated to estimate the error in $k_{cat}/K_M$.

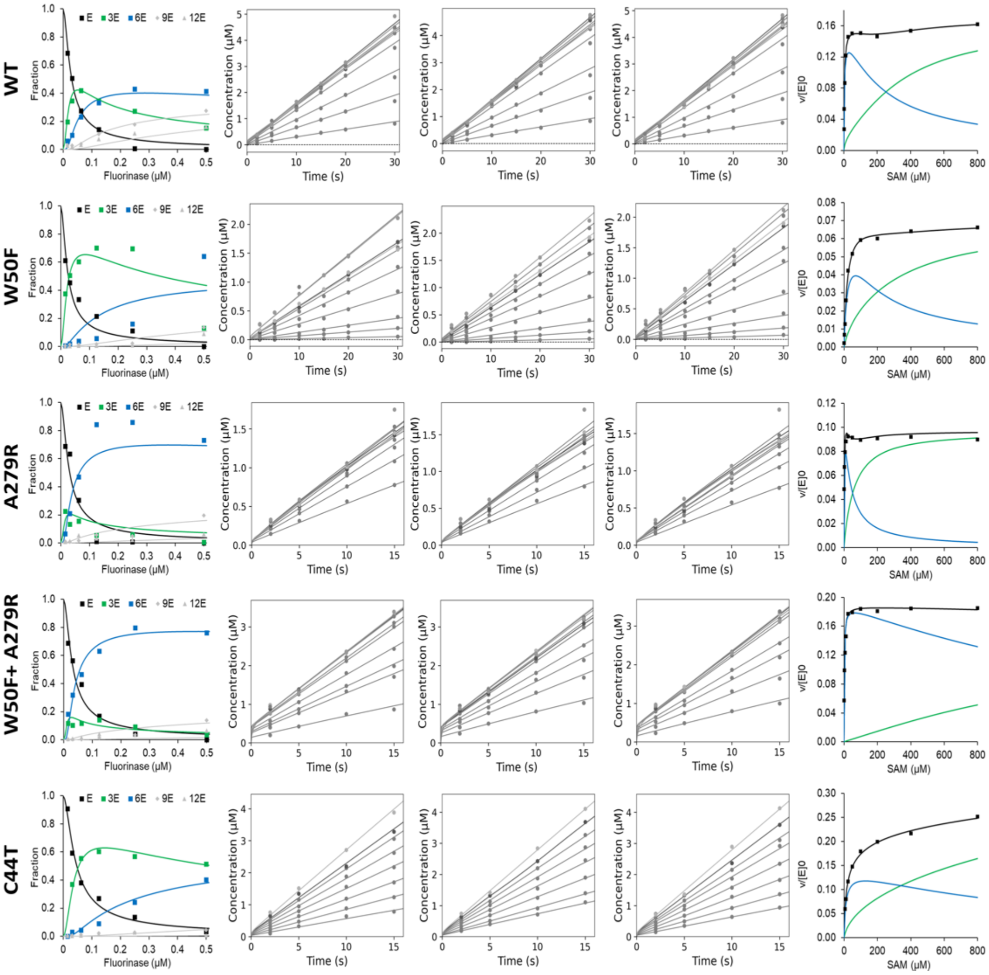

**Fig. S22.**

**Global analysis and comparison of wild-type and engineered variants of FlA1.** Mass photometry data (left) were collected in 30 mM HEPES, pH 7.5, at 20 °C, with 75 mM KF and 800 μM SAM. Raw kinetic data (middle) collected in triplicate across 1.56–800 μM SAM with 75 mM KF in 50 mM HEPES, pH 7.8, at 37 °C. The SAM concentration dependence of initial velocity (right), data points show an average of triplicate measurements. Solid lines represent the global fit.

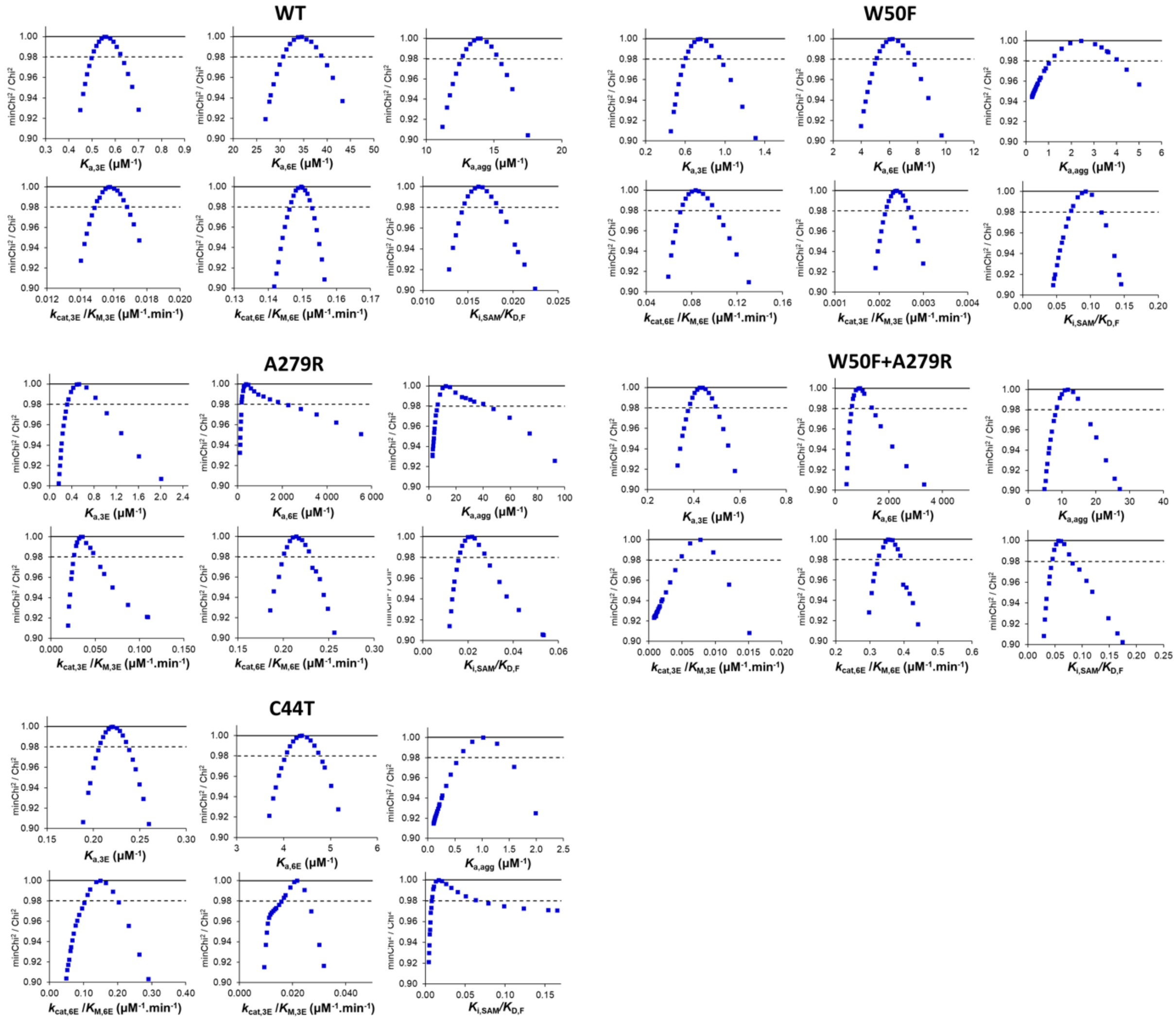

**Fig. S23.**

**Confidence contour analysis.** Individual figures represent the dependence of the error on the respective parameter while varying all other parameters to achieve the best fit. In every case, the dashed line shows the $\chi^2$ threshold (0.98) used to establish confidence intervals, as reported in **Table S4**.

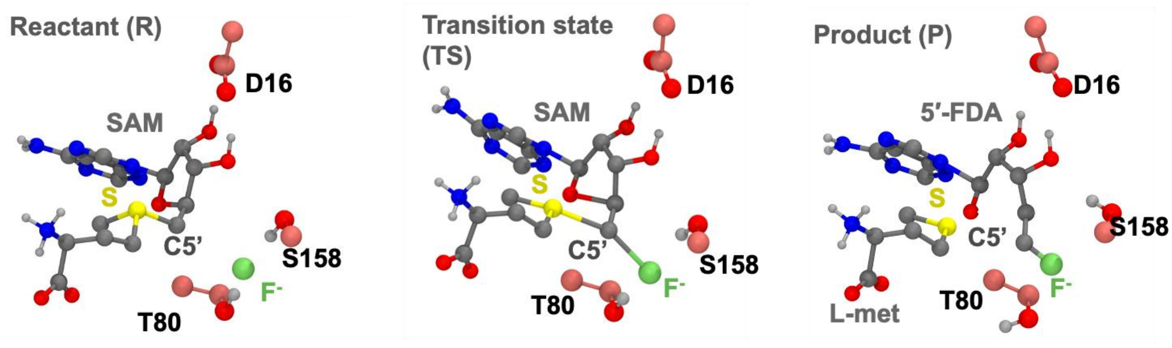

**Fig. S24.**

Schematic representation of the fluorinase reaction mechanism, showing the catalytic itinerary from the reactant state (R, left), where SAM and $F^-$ are positioned in the active site, through the transition state (TS, middle), where partial bond formation and breaking occur, to the product state (P, right), where the fluorinated product is formed (5’-FDA) and by product (L-methionine) are formed. Key active-site residues D16, T80, and S158 facilitate the reaction.

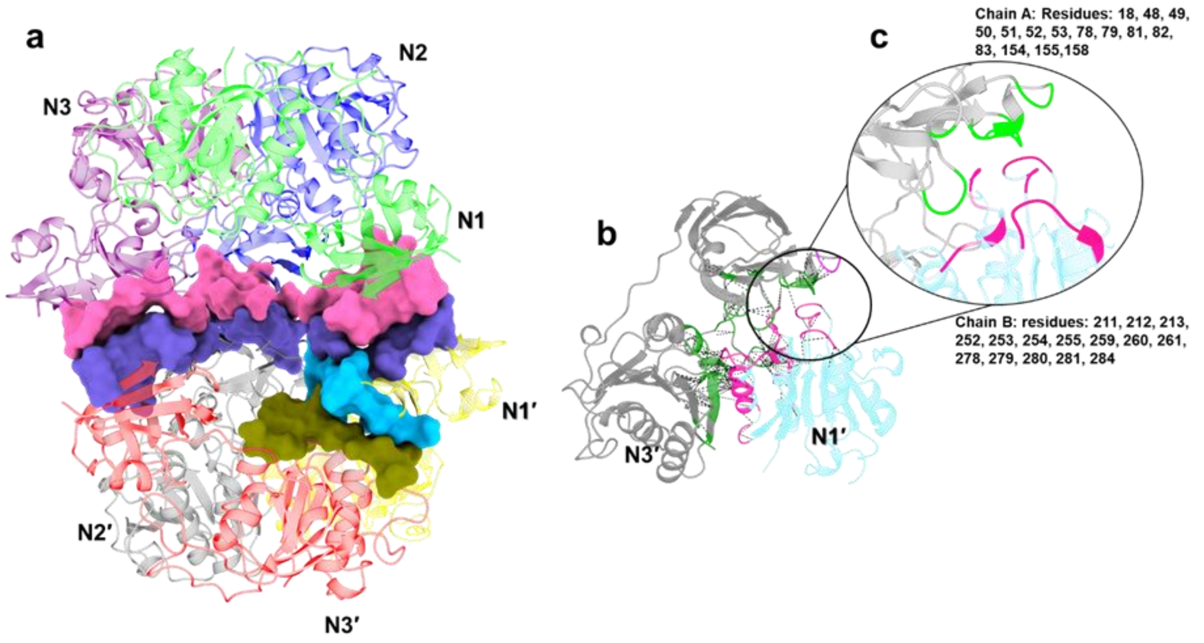

**Fig. S25.**

**a)** Interface region between trimers selected for constructing CV in hexamer simulations. **b)** Three-dimensional structure of dimer interface between N1′ and N3′ in a trimer. **c)** The residues of dimeric region selected for constructing CVs.

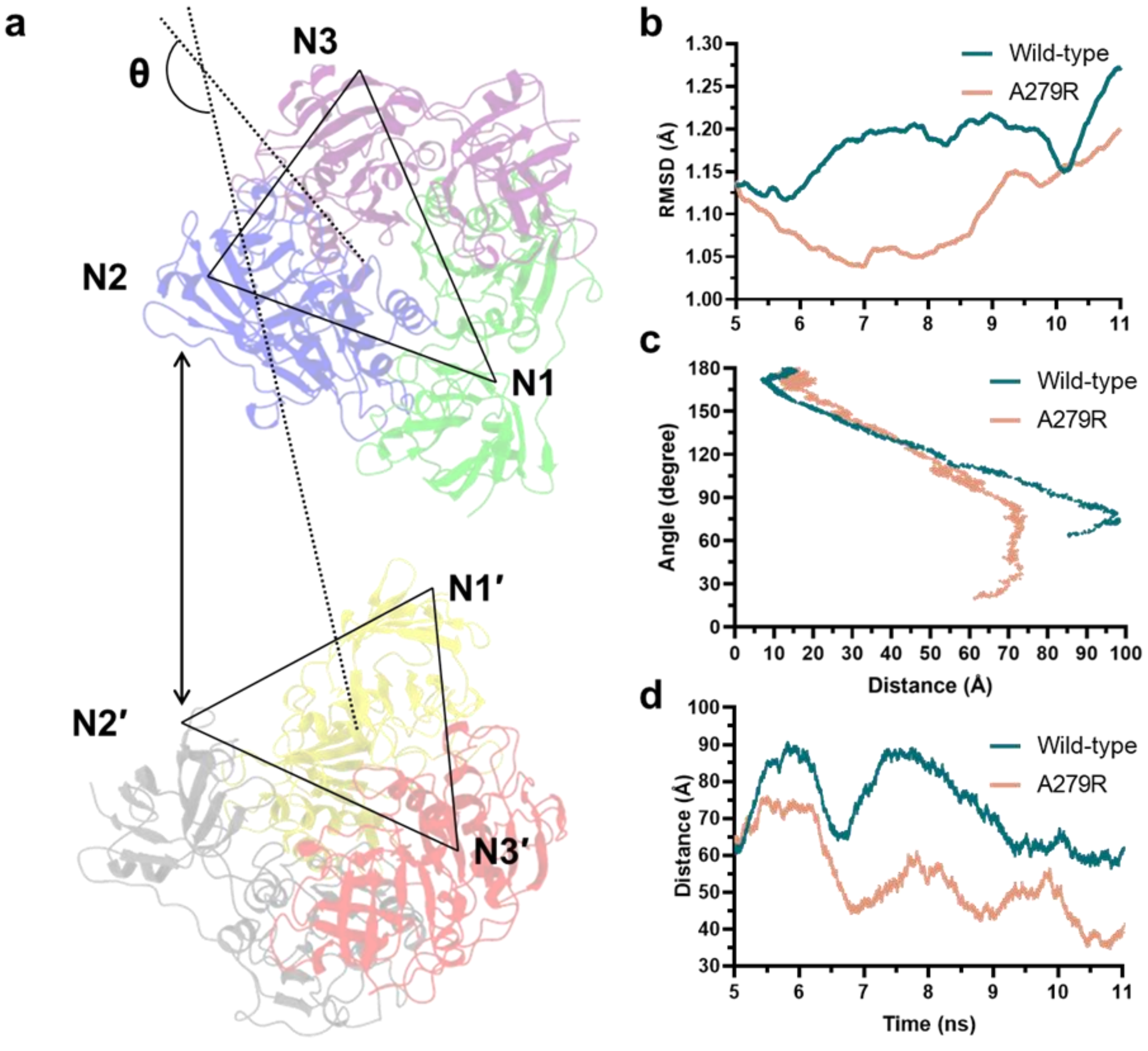

**Fig. S26.**

**a)** A schematic representation of calculation of planar angle ‘θ’ between the normal of the planes derived from the trimers. In each chain, the Cα atom of Val243 was considered as a fixed point. A plane was defined between the Cα atoms of V243 in chains N1, N2, and N3. Similarly, a second plane was defined between the Cα atoms of V243 in chains N1′, N2′, and N3′. **b)** RMSD plot comparing trimer stability in wild-type and A279R. R279 of N1 forms a stabilizing salt bridge with E53 of N3. Additional interactions with W50, I268, and N280 further stabilize R279 and reinforce the dimeric interface. **c)** Pathway of hexamer assembly plotted as distance vs. planar angle. **d)** Distance over time graph highlighting event E1 in both wild-type and A279R which is highlighted with the double-sided arrow in a.

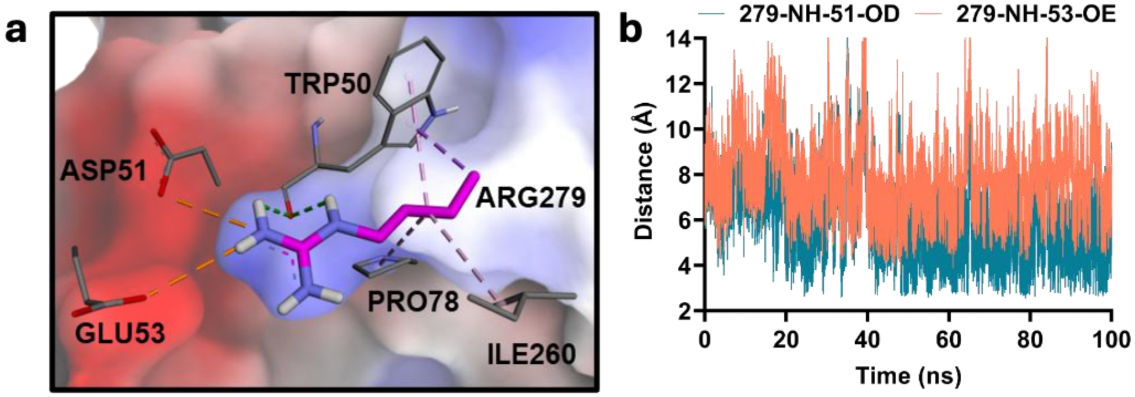

**Fig. S27.**

The metadynamics simulations shows R279 of N1 forming a prominent salt-bridge with D51 and E53 residues of N2. a) The electrostatic surface representation overlaid on the interactions of Arg with nearby residues such as W50, D51, E53, P78, and I260, b) a distance graph indicating the stable distance between R279NH1 and NH2 atoms with OD1 and OD2 of D51 (Blue lines) and OE1 and OE2 of E53 (Red lines) obtained from the extended well-tempered metadynamics simulations where it forms prominent salt-bridges. The R279 side chain is also stabilized by the backbone interactions with W50.

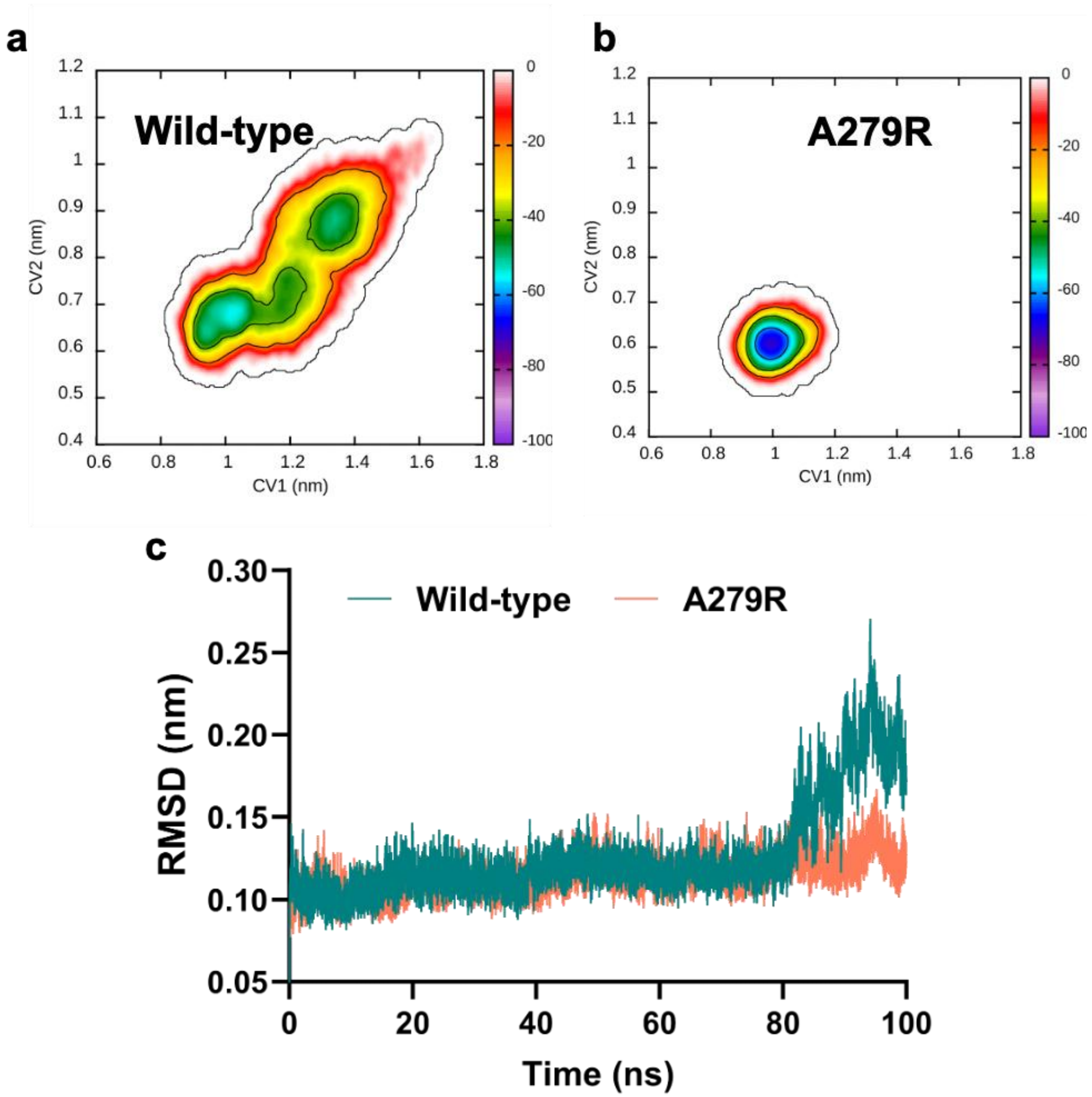

**Fig. S28.**

FES graph obtained from the well-tempered metadynamics simulation was conducted on **a)** wild and **b)** A279R trimer. **c)** RMSD calculated for wild and A279R trimer complexes.

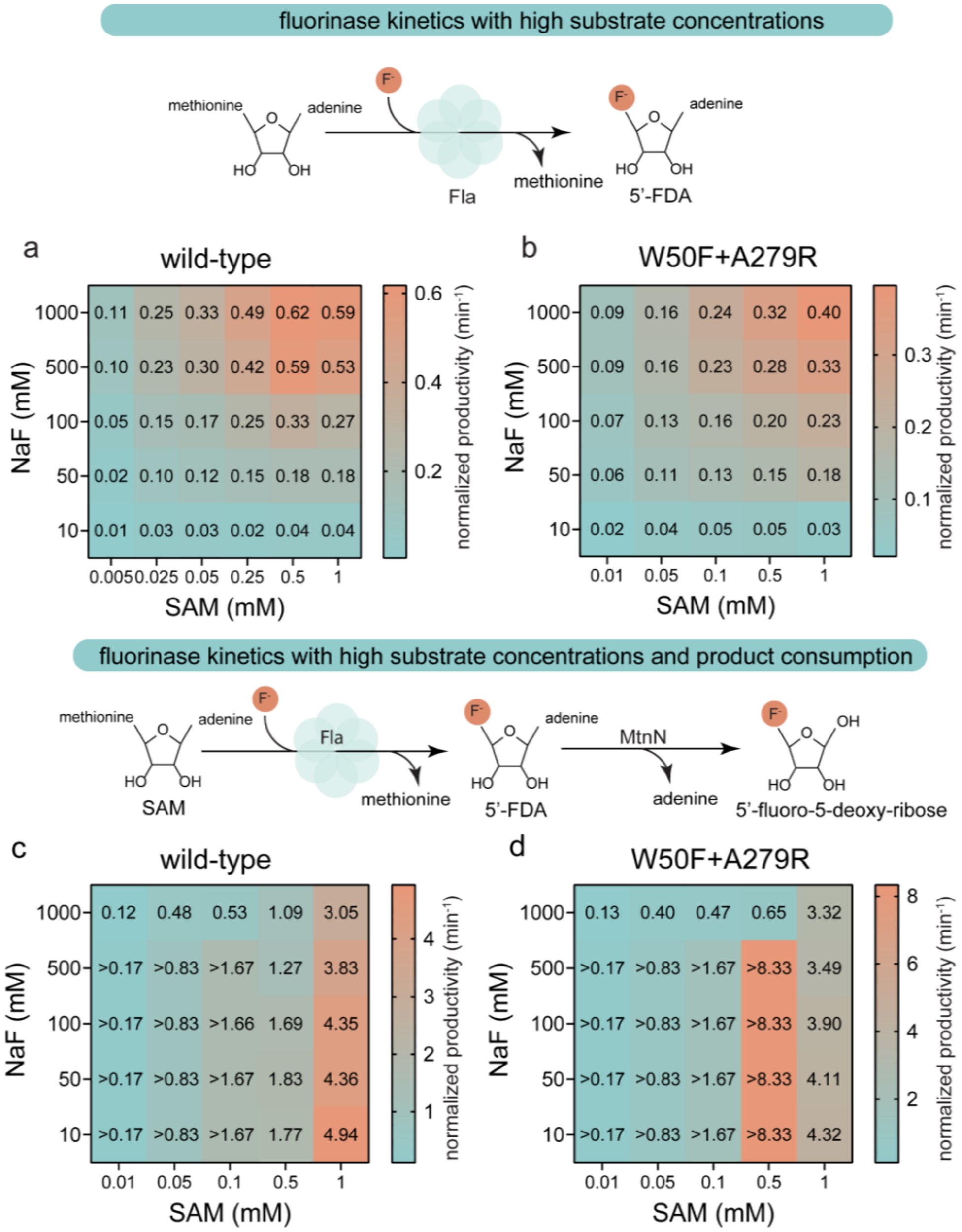

**Fig. S29.**

**Experimental validation of the predicted enzyme productivity.** Enzyme productivity was evaluated by measuring product formation at the 2-hour end-point. Values are shown as normalized productivity, defined as product formed per unit of enzyme and per unit time. For coupled reactions (FLa1 + MtnN), adenine formation was used as a proxy for product formation. The kinetic data were collected from 0.01 to 1 mM of SAM, 0.5 $\mu$M FlA1, 0.5 $\mu$M MtnN, 10-1000 mM NaF in 30 mM HEPES, pH 7.8 at 37 °C. An "*" indicates total substrate consumption after 2 h. The wild-type (**a**) and W50F+A279R mutant (**b**) fluorinase show the highest productivity with the highest substrate concentrations. The nucleosidase, MtnN, hydrolyses 5'-FDA to adenine and 5-fluoro-5-deoxy-ribose. In the coupled reaction of the fluorinase and MtnN, the wild-type shows the highest productivity (~5 min$^{-1}$) at 10 mM NaF and 1 mM SAM (**c**). While the mutant has a >8.33 min$^{-1}$ (complete substrate conversion) with

0.5 mM SAM and 10-500 mM NaF. For both variants 1000 mM NaF shows inhibitory effect on the reaction in comparison to 500 mM.

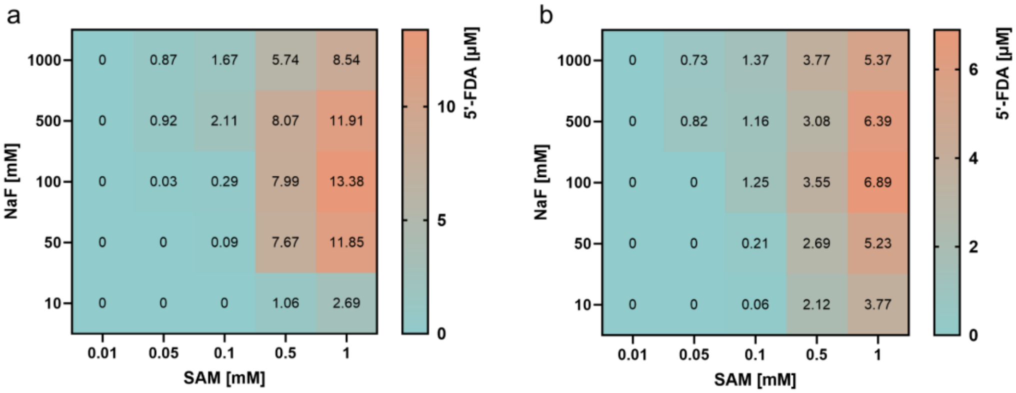

**Fig. S30.**

5’-FDA remaining in coupled reaction with wild-type (a) and W50F+A279R mutant (b).

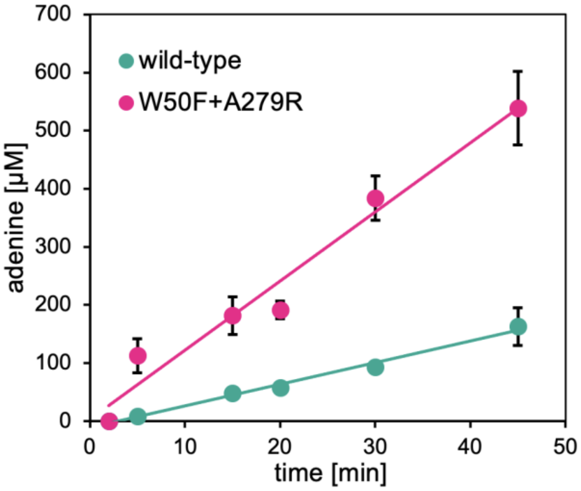

**Fig. S31.**

Adenine production of the wild-type and W50F+A279R fluorinase under optimal conditions. 500 mM NaF, 0.5 mM SAM, 1 µM MtnN, 0.5 µM Fla1, 37 °C, 30 mM HEPES (pH = 7.8)

**Table S1.**

Amino acids most suitable for substitution of cysteine redisues in the flurinase sequence predicted by HotSpot Wizard[32].

| residue | location | frequencies of amino acids | recommended substitution |
|---|---|---|---|
| C27 | buried, second shell of the active site | Cys (48%), Met (39%), Gly (6%) | C27M |
| C35 | surface, with no contacts | Cys (57%), Ala (27%), Asn (6%) | C35A |
| C44 | surface, interface of two units | Thr (42%), Ser (27%), Cys (27%) | C44T |

**Table S2.**

**Kinetic parameters of the FlA1+C44T reaction mechanism.** Constants $k_{+1}$, $k_{+2}$, $k_{-4}$, $k_{-5}$, $k_{+6}$ and $k_{+7}$ are displayed in units μM$^{-1}$ min$^{-1}$. Constants $k_{-1}$, $k_{-2}$, $k_{+3}$, $k_{-3}$, $k_{+4}$, $k_{+5}$, $k_{-6}$ and $k_{-7}$ are displayed in units min$^{-1}$. Lower and upper boundaries were calculated from the FitSpace confidence contour analysis by setting the min$\chi^2/\chi^2$ limit to 0.98. $k_{+4}$ was fixed at 10000 (⊗) and was used with $k_{-4}$ to calculate the dissociation constant ($K_4 = k_{+4}/k_{-4}$).

| reaction step | | value | lower boundary | upper boundary |
|---|---|---|---|---|
| E + $F^-$ = E·$F^-$ | $k_{+1}$ | 0.00713 | 0.00652 | 0.00786 |
| | $k_{-1}$ | 1080 | 1040 | 1210 |
| E·$F^-$ + SAM = E·$F^-$·SAM | $k_{+2}$ | 3.43 | 3.15 | 3.77 |
| | $k_{-2}$ | 174 | 149 | 203 |
| E·$F^-$·SAM = E·5’-FDA·Met | $k_{+3}$ | 102 | 86.9 | 115 |
| | $k_{-3}$ | 0.00419 | 0.00395 | 0.00446 |
| E·5’-FDA·Met = E·5’-FDA + Met | $k_{+4}$ | ⊗10000 | | |
| | $k_{-4}$ | 39.7 | 31.8 | 49.6 |
| E·5’-FDA = E + 5’-FDA | $k_{+5}$ | 17.1 | 15.3 | 19.6 |
| | $k_{-5}$ | 219 | 202 | 230 |
| E + SAM = E·SAM | $k_{+6}$ | 32.5 | 31.9 | 36.9 |
| | $k_{-6}$ | 17 | 13.1 | 185 |
| E·5’-FDA + $F^-$ = $F^-$·E·5’-FDA | $k_{+7}$ | 0.00435 | 0.00393 | 0.00664 |
| | $k_{-7}$ | 199 | 188 | 311 |

**Table S3.**

**The equilibrium constants obtained by global fit.** The best-fit estimates were obtained by nonlinear regression based on numerical integration of the rate equations. The standard error (± s.e.) was calculated from the covariance matrix during nonlinear regression. Confidence intervals (lower and upper limits) of the parameters were obtained by confidence contour analysis for χ2 threshold of 0.98. Experiments were conducted for the free fluorinase in buffer (30 mM HEPES, pH 7.5, 20°C), the fluorinase in the presence of fluoride substrate (75 mM KF), and in the presence of both substrates (75 mM KF + 800 µM SAM).

| | free enzyme | | | | enzyme+KF | | | | enzyme+KF+SAM | | | |
|---|---|---|---|---|---|---|---|---|---|---|---|---|
| | value | s.e. | lower | upper | value | s.e. | lower | upper | value | s.e. | lower | upper |
| **wt** | | | | | | | | | | | | |
| $K_{a,3E}$ ($\mu M^{-1}$) | 2.37 | 0.47 | 2.05 | 2.59 | 13.4 | 1.6 | 12.4 | 14.5 | 0.563 | 0.046 | 0.529 | 0.599 |
| $K_{a,6E}$ ($\mu M^{-1}$) | 19.7 | 3.2 | 17.6 | 22.0 | 3.56 | 0.22 | 3.42 | 3.72 | 37.3 | 3.6 | 35.0 | 40.0 |
| $K_{a,agg}$ ($\mu M^{-1}$) | 18.7 | 2.6 | 16.8 | 20.8 | 3.92 | 0.66 | 3.46 | 4.41 | 15.2 | 1.3 | 14.2 | 16.2 |
| **W50F** | | | | | | | | | | | | |
| $K_{a,3E}$ ($\mu M^{-1}$) | 1.24 | 0.14 | 1.15 | 1.35 | 3.22 | 0.72 | 2.73 | 3.83 | 0.698 | 0.085 | 0.633 | 0.758 |
| $K_{a,6E}$ ($\mu M^{-1}$) | 11.0 | 1.3 | 10.1 | 12.0 | 11.3 | 1.6 | 10.2 | 12.6 | 2.28 | 0.48 | 1.96 | 2.58 |
| $K_{a,agg}$ ($\mu M^{-1}$) | 10.1 | 1.7 | 8.8 | 11.4 | 3.41 | 1.04 | 2.58 | 4.26 | 3.29 | 2.12 | 1.77 | 4.12 |
| **A279R** | | | | | | | | | | | | |
| $K_{a,3E}$ ($\mu M^{-1}$) | 19.6 | 15.1 | 12.5 | 30.6 | 2.82 | 1.08 | 2.16 | 3.72 | 0.444 | 0.080 | 0.391 | 0.510 |
| $K_{a,6E}$ ($\mu M^{-1}$) | 13.2 | 2.9 | 11.5 | 15.1 | 20.1 | 5.0 | 17.1 | 23.3 | 1 351 | 1 059 | 862 | 2732.2 |
| $K_{a,agg}$ ($\mu M^{-1}$) | 3.68 | 1.76 | 2.62 | 4.59 | 3.18 | 1.88 | 2.04 | 4.39 | 14.5 | 7.5 | 9.8 | 22.6 |
| **W50F+A279R** | | | | | | | | | | | | |
| $K_{a,3E}$ ($\mu M^{-1}$) | 0.326 | 0.072 | 0.261 | 0.379 | 1.31 | 0.28 | 1.11 | 1.52 | 0.437 | 0.055 | 0.394 | 0.478 |
| $K_{a,6E}$ ($\mu M^{-1}$) | 4.18 | 1.17 | 3.28 | 4.88 | 22.7 | 3.9 | 20.2 | 25.7 | 870 | 363 | 676 | 1 161 |
| $K_{a,agg}$ ($\mu M^{-1}$) | 5.26 | 2.05 | 3.58 | 6.58 | 1.88 | 1.07 | 1.13 | 2.35 | 11.7 | 3.7 | 8.9 | 14.6 |
| **C44T** | | | | | | | | | | | | |
| $K_{a,3E}$ ($\mu M^{-1}$) | 0.272 | 0.049 | 0.228 | 0.307 | 0.731 | 0.051 | 0.694 | 0.769 | 0.221 | 0.020 | 0.207 | 0.2358 |
| $K_{a,6E}$ ($\mu M^{-1}$) | 3.24 | 0.78 | 2.65 | 3.76 | 3.56 | 0.25 | 3.38 | 3.75 | 4.39 | 0.46 | 4.07 | 4.76 |
| $K_{a,agg}$ ($\mu M^{-1}$) | 3.80 | 1.53 | 2.54 | 4.76 | 0.620 | 0.356 | 0.347 | 0.775 | 1.02 | 0.54 | 0.65 | 1.27 |
| **C44T+W50F+A279R** | | | | | | | | | | | | |
| $K_{a,3E}$ ($\mu M^{-1}$) | 0.492 | 0.084 | 0.422 | 0.552 | 1.61 | 0.18 | 1.50 | 1.72 | 0.503 | 0.047 | 0.465 | 0.538 |
| $K_{a,6E}$ ($\mu M^{-1}$) | 4.85 | 0.97 | 4.07 | 5.65 | 9.90 | 0.98 | 9.26 | 10.50 | 10.6 | 1.0 | 9.9 | 11.4 |
| $K_{a,agg}$ ($\mu M^{-1}$) | 4.24 | 1.36 | 3.13 | 5.29 | 1.37 | 0.95 | 0.70 | 1.72 | 1.28 | 0.50 | 0.88 | 1.59 |

**Table S4.**

**Kinetic parameters for wild-type and engineered variants of FlA1.** The table presents best-fit estimates of kinetic parameters derived through nonlinear regression based on the numerical integration of rate equations. The standard error (± s.e.) was calculated using the covariance matrix obtained during the regression process. Confidence intervals (lower and upper bounds) for the parameters were determined through confidence contour analysis with a $\chi^2$ threshold of 0.98. Experiments were conducted under the following conditions: 50 mM HEPES buffer, pH 7.8, at 37 °C.

| | | wild type | | | | W50F | | | | A279R | | | |
|---|---|---|---|---|---|---|---|---|---|---|---|---|---|
| parametr | unit | value | ± s.e. | lower limit | upper limit | value | ± s.e. | lower limit | upper limit | value | ± s.e. | lower limit | upper limit |
| $K_{a,3E}$ | $\mu M^{-1}$ | 0.56 | ± 0.03 | 0.51 | 0.62 | 0.75 | ± 0.08 | 0.60 | 0.94 | 0.5 | ± 0.2 | 0.3 | 0.8 |
| $K_{a,6E}$ | $\mu M^{-1}$ | 34.6 | ± 1.7 | 30.6 | 38.8 | 6.2 | ± 0.5 | 5.1 | 7.3 | 380 | ± 30 | 170 | 1830 |
| $K_{a,agg}$ | $\mu M^{-1}$ | 14.0 | ± 0.1 | 12.7 | 15.2 | 2.4 | ± 0.6 | 1.2 | 3.0 | 12.4 | ± 4.1 | 6.4 | 38.0 |
| $k_{cat,3E}/K_{m,3E}$ | $\mu M^{-1}.min^{-1}$ | 0.016 | ± 0.001 | 0.015 | 0.017 | 0.0024 | ± 0.0002 | 0.0022 | 0.0027 | 0.036 | ± 0.006 | 0.026 | 0.048 |
| $k_{cat,6E}/K_{m,6E}$ | $\mu M^{-1}.min^{-1}$ | 0.150 | ± 0.001 | 0.146 | 0.152 | 0.08 | ± 0.02 | 0.07 | 0.10 | 0.21 | ± 0.05 | 0.07 | 0.27 |
| $K_{i,SAM}/K_{d,F}$ | μM | 0.016 | ± 0.001 | 0.015 | 0.017 | 0.09 | ± 0.01 | 0.07 | 0.10 | 0.021 | ± 0.005 | 0.016 | 0.027 |

| | | W50F+A279R | | | | C44T | | | |
|---|---|---|---|---|---|---|---|---|---|
| parametr | unit | value | ± s.e. | lower limit | upper limit | value | ± s.e. | lower limit | upper limit |
| $K_{a,3E}$ | $\mu M^{-1}$ | 0.44 | ± 0.03 | 0.38 | 0.49 | 0.22 | ± 0.01 | 0.21 | 0.24 |
| $K_{a,6E}$ | $\mu M^{-1}$ | 870 | ± 120 | 620 | 1360 | 4.4 | ± 0.2 | 4.1 | 4.7 |
| $K_{a,agg}$ | $\mu M^{-1}$ | 11.8 | ± 0.1 | 8.5 | 15.3 | 1.0 | ± 0.2 | 0.7 | 1.3 |
| $k_{cat,3E}/K_{m,3E}$ | $\mu M^{-1}.min^{-1}$ | 0.008 | ± 0.001 | 0.005 | 0.010 | 0.022 | ± 0.002 | 0.016 | 0.024 |
| $k_{cat,6E}/K_{m,6E}$ | $\mu M^{-1}.min^{-1}$ | 0.35 | ± 0.01 | 0.33 | 0.39 | 0.15 | ± 0.03 | 0.11 | 0.19 |
| $K_{i,SAM}/K_{d,F}$ | μM | 0.061 | ± 0.001 | 0.045 | 0.077 | 0.017 | ± 0.009 | 0.008 | 0.063 |

**Movie S1.**

**Diffusion of fluoride into the active site**. Catalytic residues (T80, S158) are depicted as orange sticks, and other active site residues in teal sticks. Only water molecules and residues in the active site that directly interact with the fluoride are displayed for clarity.

**Movie S2.**

**Elucidated Reaction mechanism** for formation of 5’FDA from SAM, showing the Reactant (R), Transition State (TS), and Product (P).

**Movie S3.**

**Formation of Hexamer complex** in **A)** A279R mutant, **B)** wild-type. Chains are coloured differently to distinguish between the subunits of the hexamer.
